# Development of a hybrid Bayesian network model for predicting acute fish toxicity using multiple lines of evidence

**DOI:** 10.1101/750935

**Authors:** S. Jannicke Moe, Anders L. Madsen, Kristin A. Connors, Jane M. Rawlings, Scott E. Belanger, Wayne G. Landis, Raoul Wolf, Adam D. Lillicrap

**Author notes:** Corresponding author: S. Jannicke Moe. **Abbreviations** AFT: acute fish toxicity; BN: Bayesian network; CV: coefficient of variation; EC50: effect concentration for 50% of the test individuals; ECHA: European Chemicals Agency; FET: fish embryo toxicity; LC50: lethal concentration for 50% of the test individuals; QSAR: quantitative structure–activity relationship; REACH: registration, evaluation, authorisation and restriction of chemicals; WOE: weight of evidence.

## Abstract

A Bayesian network was developed for predicting the acute toxicity intervals of chemical substances to fish, based on information on fish embryo toxicity (FET) in combination with other information. This model can support the use of FET data in a Weight-of-Evidence (WOE) approach for replacing the use of juvenile fish. The BN predicted correct toxicity intervals for 69%-80% of the tested substances. The model was most sensitive to components quantified by toxicity data, and least sensitive to components quantified by expert knowledge. The model is publicly available through a web interface. Further development of this model should include additional lines of evidence, refinement of the discretisation, and training with a larger dataset for weighting of the lines of evidence. A refined version of this model can be a useful tool for predicting acute fish toxicity, and a contribution to more quantitative WOE approaches for ecotoxicology and environmental assessment more generally.

**Highlights:**

- A Bayesian network (BN) was developed to predict the toxicity of chemicals to fish
- The BN uses fish embryo toxicity data in a quantitative weight-of-evidence approach
- The BN integrates physical, chemical and toxicological properties of chemicals
- Correct toxicity intervals were predicted for 69-80% of test cases
- The BN is publicly available for demonstration and testing through a web interface

## 1. Introduction

Weight-of-evidence (WOE) is a term commonly used about the assembly, weighting, and integration of multiple pieces of evidence for environmental assessment and decision-making. However, the term has been used in multiple ways, including metaphorical, theoretical and methodological, without consensus about its meaning (Weed, 2005). Historically, environmental risk assessors have relied upon narrative and qualitative approaches such as expert knowledge, or limited quantitative methods such as direct scoring to integrate multiple lines of evidence (Linkov et al., 2009). These historical methods lacked transparency and reproducibility. A review of WOE methods applied for ecological risk assessment (Linkov et al., 2009) showed that only 9 out of 44 published studies had used a quantitative approach (scoring, indexing or other type of quantification). Therefore, WOE approaches have been criticised for being too vague, intransparent and subjective (Linkov et al., 2016). Nevertheless, methods are needed for integrating evidence and assess their uncertainty to support decision-making for environmental protection. These processes would benefit from more quantitative and rigorous approaches to WOE.

More systematic frameworks for WOE approaches in scientific assessments have recently been proposed, for example by the European Chemicals Agency (ECHA, 2016), by the European Food Safety Authority (EFSA Scientific Committee et al., 2017) and by the US Environmental Protection Agency (USEPA, 2016) to assist ecological assessment. The basic steps in these frameworks are to (1) assemble evidence, (2) to weight the evidence by assigning scores depening on e.g. relevance, strength and reliability, and (3) to intergrate the evidence (weigh the body of evidence). Suter et al. describe how the US EPA framwork can be used to infer qualitative properties (2017a) as well as quantitative estimates (2017b). Use of such frameworks can increase the consistency and rigor of WOE practices and provide greater transparency than ad hoc and narrative-based approaches.

The weight-of-evidence was first proposed as a Bayesian statistical approach (Good, 1960). In Bayesian statistics, a model is based on updating prior beliefs in (probabilities of) a hypothesis after evaluating of evidence in order to achieve a posterior belief. In this context, the WOE is defined as the logarithm of the Bayes factor, which is calculated as the ratio of the posterior odds to the prior odds (Good, 1985). In this paper, we return to the Bayesian origin of the WOE term and develop a Bayesian network to combine lines of information in a probabilistic model to support risk assessment of chemicals.

Environmental risk assessment is a process for quantifying the probability of an adverse effect from a chemical exposure. Traditionally, acute toxicity data from three trophic levels (i.e., fish, invertebrates, algae) are required for risk assessments. However, ethical considerations and recent EU regulation have required that the number of animals used for toxicity testing is kept at a minimum, and that the use of vertebrate species (such as fish) should be avoided as far as possible (OECD, 1996). Therefore, various alternative methods have been developed and proposed to replace the use of live fish in toxicity testing (Lillicrap et al., 2016), including predictive toxicity modelling (Luechtefeld et al., 2018).

Legislative authorities in some geographic regions have encouraged the use of fish embryos, which are in this context considered as a non-protected life stage, over more developed life stages such as juvenile fish. The test of fish embryo toxicity (FET; OECD method 236) (OECD, 2013) has therefore been proposed and evaluated as an alternative to using juvenile fish for testing acute fish toxicity (AFT; OECD method 203) (OECD, 1992) (Busquet et al., 2014). Previous studies show a good correlation of FET with the standard acute fish toxicity (AFT) test (Belanger et al., 2013; Rawlings et al., 2019). Nevertheless, it has been found that the fish embryo toxicity test alone is currently not sufficient to replace the acute fish toxicity data as required by the European REACH regulation (Regulation, Evaluation, Authorisation and Restriction of Chemicals) (Sobanska et al., 2018). However, the authors have suggested that “the test may be used within weight-of-evidence approaches together with other independent, relevant, and reliable sources of information”.

To this end, we have developed a Bayesian network (BN) model for integrating fish embryo toxicity data with other information to predict the acute toxicity to (juvenile) fish for any given chemical substance. A Bayesian network is a probabilistic graphical model over a set of random variables. Bayesian networks are commonly used in environmental modelling (Aguilera et al., 2011; Barton et al., 2012; Landuyt et al., 2013). For example, there are numerous of BN models developed for assessment and management of water quality (Barton, 2014; Borsuk et al., 2012; Borsuk et al., 2004; Moe et al., 2019; Moe et al., 2016). BN modelling has been applied more rarely within ecotoxicology and ecological risk assessment, but recent publications have demonstrated the applicability of BN modelling in these fields (Graham et al., 2019; Landis et al., 2017; Landis et al., 2019; Lehikoinen et al., 2015). Compared to traditional qualitative WOE approaches, the network structure and probabilistic framework of BN provide advantages by capturing the impacts of multiple sources of quantifiable uncertainty on predictions of ecological risk (Carriger et al., 2016). Another example of applying a BN model for a quantitative WOE and testing strategy is provided by Jaworska et al. (2015).

Here, we develop a BN model for integrating four lines of evidence in a quantitative framework: (1) information on physical and chemical properties of the substance, (2) toxicity data for chemically related substances, (3) toxicity data for other species (crustaceans and algae) and (4) fish embryo toxicity data. By implementing this WOE model in a Bayesian network, we aim to meet the demands for making WOE approaches more transparent, structured and quantitative (Linkov et al., 2016).

The purpose of this paper is to describe the development, parameterisation and evaluation of the BN model. The paper will also present an online web interface to the model. More detailed information on this model’s performance for specific chemical substances and endpoints, and the implications for current legislation and guidelines for toxicity testing, are addressed by Lillicrap et al. (2019). Feedback from researchers, regulators or other potential users to this first BN version will be used for improving future versions of the model, with the aim of making it a useful tool in a WOE approach for predicting acute fish toxicity.

## 2. Materials and methods

### 2.1 Data

An expanded version of the Threshold Database (Rawlings et al., 2019) was obtained from Procter & Gamble and used to construct the BN. This database contains acute freshwater toxicity values for fish (AFT), fish embryos (FET), algae, and invertebrates for 237 chemical substances. The toxicity values are measured as EC50 (Effect Concentration for 50% of the test individuals) for non-lethal effects, and as LC50 (Lethal Concentration of 50% of the test individuals) for lethal effects. The dataset used for development of this model, which will be referred to as “our dataset”, contained the following toxicity data:

- Algae: 264 values of EC50 based on OECD test no. 201 acute algal growth inhibition tests (OECD, 2006).
- *Daphnia*: 1164 values EC50 based on OECD test no. 202 *Daphnia sp*. acute immobilisation tests (OECD, 2004), using crustaceans of the genus *Daphnia* (daphnids).
- Juvenile fish: 1459 LC50 values based on OECD test no. 203 acute fish toxicity test (OECD, 1992).
- Fish embryo: 541 LC50 values based on OECD test no. 236 Fish embryo test (OECD, 2013).

In addition to the data obtained from laboratory assays, modelled acute fish LC50 values were calculated using quantitative structure-activity relationships (QSARs) for each chemical, including the US EPA Ecological Structure Activity Relationship (ECOSAR 1.11) models and the Danish QSAR database (extracting values for Leadscope and SciQSAR SARs). Data were then averaged for each chemical, resulting in 152 QSAR values representing modelled toxicity to juvenile fish. More details on QSAR model selection, equation acceptability criteria and results can be found in (Lillicrap et al., 2019).

### 2.2 BN model objective and structure

A BN model consists of a directed, acyclic graph with nodes representing the random variables and arrows (archs) representing conditional probability distributions (Kjærulff and Madsen, 2008). Each node has a probability distribution conditional on its parents in the graph. The probability distributions quantify the strengths for the dependence relations defined by the structure of the graph. The nodes are typically defined by limited number of discrete states (categories or intervals) which are quantified by a prior probability distribution. New evidence is combined with the prior probabilities to calculate posterior probability distributions, using Bayes’ rule from 1763 (see e.g. Linkov et al., 2016).

The objective of the BN model presented here is to predict the acute toxicity of a chemical substance to juvenile fish, corresponding to the interval of LC50 values from the AFT assay (OECD, 1992), by integrating toxicity data from testing of fish embryos (OECD, 2013) with other relevant physical, chemical and toxicological information for the given substance. The BN model is structured along the four lines of evidence (Figure 1), representing different types of information for a given chemical substance for which a user wants to predict the acute fish toxicity. The parameterisation of all lines is described in Supplementary material (Tables S.1-S.21), while more details on the assumptions are provided by Lillicrap et al. (2019).

**Figure 1.**
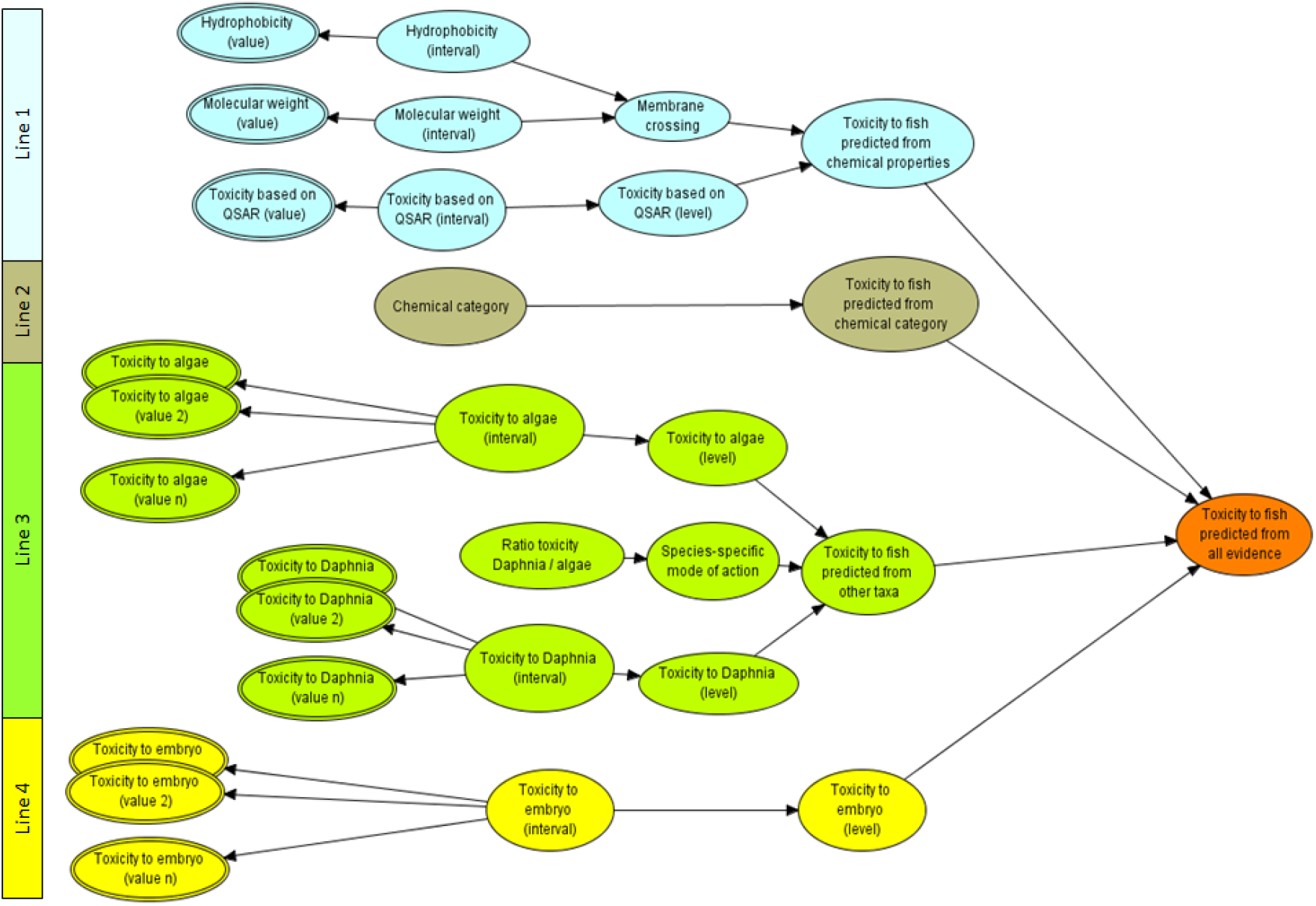
The main structure of the BN model: the four lines of evidence for predicting the interval of acute toxicity of a substance to fish. Nodes with double outline represent continuous value nodes (e.g. “Toxicity to algae (value 1)”), while nodes with single outline are interval or categorical nodes. All of the left-most nodes are input nodes for a given substance. For simplicity, only three continuous input nodes are shown for each of algae, *Daphnia* and embryo. “Value n” denotes the maximum number of input variable nodes in the web interface (currently 10).

- Line 1: Physical and chemical properties of the substance. The selected nodes quantify the size (molecular weight, g/mol), hydrophobicity (octanol water partitioning coefficient, logK_ow_) and modelled toxicity to juvenile fish based on the structure (QSAR) of the substance. Substances with a low molecular size and low hydrophobicity (i.e. high solubility) are assumed to have a high ability to cross a biological membrane. For a substance that is modelled via QSAR to be toxic (low LC50 value) and have a high ability of membrane crossing, the predicted toxicity to juvenile fish from this line of evidence will have highest probability of the higher toxicity intervals.
- Line 2: Chemical category of the substance. This line of evidence is based on existing data on toxicity of substances to juvenile fish, aggregated by chemical category. The 237 chemicals used to parametrize this model were assigned to 42 different chemical categories (Table S.11), based on model output ECOSAR and further refined by expert knowledge (S. Belanger).
- Line 3: Toxicity to other taxa, representing lower trophic levels: crustaceans (*Daphnia magna* or *D. pulex*) or unicellular algae. This line of evidence also considers whether the substance has a species-specific (or more generally, taxon-specific) mode of action by examining the ratio of *Daphnia* and algal toxicity. If the ratio of toxicity to *Daphnia* vs. algae is above 2 or below 0.5, then the substance’s mode of action is assumed to be specific to one of these taxa (e.g. algacidal), and therefore to be less toxic to fish. Conversely, if the toxicity ratio is between 0.5 and 2, then the toxicity to fish is assumed to be approximately equal to *Daphnia* and algae, and a toxicity at the same level (but with more uncertainty) is assumed for fish. The full details of these assumptions are described by Lillicrap et al. (2019).
- Line 4: Toxicity to fish embryo. The measured toxicity of a substance to fish embryos is used directly as the fourth line of evidence.

The development and application of this BN is comparable to a weight-of-evidence approach (e.g., Suter et al., 2017a), with the assignment of conditional probabilities to different variables within in a line of evidence in the BN corresponding to setting weights to pieces of evidence in a WOE. For example, assignment of low weight to a piece of evidence can be obtained by setting wide probability distributions (representing high uncertainty or variability) in the relation from this node to its child node. Within a line of evidence (e.g. Line 3 “Toxicity to other taxa”, Figure 1), the calculation of posterior probabilities for the last child node of this line (“Toxicity to fish predicted from other taxa”) accumulates the weights given to all parent nodes in this line.

When using the BN to predict the toxicity to juvenile fish from all lines of evidence for a chemical of concern, the weighing of the total evidence for each hypothesis in a WOE (Suter et al., 2017a) can correspond to calculating the posterior probability of each toxicity level (“very low”, “low” etc.) in the BN. For example, given two lines of evidence with posterior probability distributions both centered around “high toxicity”, the line with the more narrow probability distribution will typically contribute more to the posterior probability of “high toxicity” in the final child node (“Toxicity to fish predicted from other taxa”). In a WOE approach, this would correspond to this line of evidence having a higher weight for the hypothesis “high toxicity to juvenile fish” than the line with a wider probability distribution.

The BN was implemented in the software HUGIN Researcher version 8.7, developed by HUGIN EXPERT A/S (http://www.hugin.com). The model is available through a web interface (described in Section 3), which is published on the demonstration web site http://demo.hugin.com/FET. The aim this web interface is to facilitate feedback for further improvement of the model, as recommended by Marcot (2017).

### 2.3 Model parametrization

#### 2.3.1 Node types and discretization

BN models are usually constructed with discrete nodes with a low number of states (categories or intervals) (Kjærulff and Madsen, 2008). An area of recent interest and progress is the development of continuous BN models (Qian and Miltner, 2015), where quantitative variables are not discretised into intervals but instead are represented by equations or statistical distributions, or hybrid BNs (e.g., Aguilera et al., 2010), which contain both discrete and continuous nodes (Marcot and Penman, 2019). For our model, categorical nodes with few (5) states would be the most convenient for parameterization of conditional probability tables in cases where expert knowledge was required. On the other hand, continuous nodes would be preferable to optimise the use of the continuous input values (e.g. measured toxicity values) and their variability. Consequently, this model is a hybrid BN with both discrete and continuous nodes (Figure 1). We let interval nodes represent the true, but unknown toxicity of a substance. The interval nodes correspond to the categorical nodes, but with two additional (more extreme) states (Table 1). The true toxicity was considered to cause the observed values, therefore the toxicity observations were modelled by continuous nodes as children (realisations) of the interval nodes. The parameterisation of all nodes, i.e. priors, conditional probabilities and conditional density functions, is described in Supplementary material (Tables S.1 to S.21), following the recommendations for good practice in BN modelling (Chen and Pollino, 2012).

**Table 1.** Overview and description of nodes in the BN model. The sources of information for probability distributions are described in Section 2. “Proportions” refers to count of observations in our dataset. For more details, see Supplementary material.

| Node group | Node name | Node type | No. of states | Source of probability distribution | In-/output in web interface | Additional information |
| --- | --- | --- | --- | --- | --- | --- |
| Physical and chemical properties of the substance | Molecular weight (value) | Continuous | NA | Proportions | Input |  |
|  | Molecular weight (interval) | Interval | 2 | Proportions |  | Root node |
|  | Hydrophobicity (value) | Continuous | NA | Proportions | Input |  |
|  | Hydrophobicity (interval) | Interval | 2 | Proportions |  | Root node |
|  | Toxicity based on QSAR (value) | Continuous | NA | Statistical rule | Input |  |
|  | Toxicity based on QSAR (interval) | Interval | 7 | Proportions |  | Root node |
|  | Toxicity based on QSAR (level) | Labelled | 5 | Deterministic |  |  |
|  | Membrane crossing | Labelled | 3 | Expert judgement |  |  |
|  | Toxicity to fish predicted from chemical properties | Labelled | 5 | Expert judgement | Output | See Fig. 3a |
| Chemical category of the substance | Chemical category | Labelled | 10 | Uniform distribution | Input | Root node |
|  | Toxicity to fish predicted from chemical category | Labelled | 6 | Proportions | Output | See Fig. 3b |
| Toxicity of the substance to other taxa (algae and Daphnia) | Toxicity to algae (value $n$) | Continuous | NA | Statistical rule | Input | See Fig. 3c |
|  | Toxicity to algae (interval) | Interval | 7 | Proportions |  | Root node |
|  | Toxicity to algae (level) | Labelled | 5 | Deterministic | Output |  |
| | Toxicity to Daphnia (value $n$) | Continuous | NA | Statistical rule | Input | |
|  | Toxicity to Daphnia (interval) | Interval | 7 | Proportions |  | Root node |
|  | Toxicity to Daphnia (level) | Labelled | 5 | Deterministic | Output |  |
|  | Ratio toxicity Daphnia / algae | Continuous |  | Proportions | Calculated from input<sup>a)</sup> | Root node |
|  | Species-specific mode of action | Interval | 3 | Proportions |  |  |
|  | Toxicity to fish predicted from other taxa | Labelled | 5 | Expert judgement | Output |  |
| Toxicity of the substance to fish embryo | Toxicity to embryo (value $n$) | Continuous | NA | Statistical rule | Input | |
|  | Toxicity to embryo (interval) | Interval | 7 | Proportions |  | Root node |
|  | Toxicity to embryo (level) | Labelled | 5 | Deterministic | Output |  |
| Predicted toxicity to juvenile fish | Toxicity to fish predicted from all evidence | Labelled | 5 | Combination rule (equal weights) | Output | See Fig. 3d |

<sup>a)</sup>The web interface calculates the variable "Ratio toxicity Daphnia / algae" from the provided input values to "Toxicity to algae (value n)" and "Toxicity to Daphnia (value n)"

For the physical and chemical properties (Line 1), we chose only two intervals (Table 2) with cut-off values commonly used in ecotoxicology: molecular weight with the threshold 600 g/mol (Brooke et al., 1986) and hydrophobicity with the threshold log K_OW_5.5 (OECD, 2012). Because the use of strict cut-off criteria to define bioaccumulation potential has been criticized (Arnot et al., 2010), the child node “Membrane crossing” was defined by three intervals (low, medium and high) to allow for intermediate levels of membrane crossing potential. The mode-of-action node (Line 3) also had three intervals based on the pre-defined cut-off values of ratio <0.5 or >2 (ECETOC, 2005).

**Table 2.** Prior probabilities for interval nodes with two states: molecular weight and hydrophobicity. The probability of each state was set equal to the proportion of count of values in our dataset.

| Label | Molecular weight (interval) |  |  | Hydrophobicity (interval) |  |  |
| --- | --- | --- | --- | --- | --- | --- |
| | Interval (unit: g/mol) | Count | Probability | Interval (unit: $k_{ow}$) | Count | Probability |
| Low | 0 - 600 | 231 | 0.9914 | -inf - 5.5 | 213 | 0.9682 |
| High | 600 - inf | 2 | 0.0086 | 5.5 - inf | 7 | 0.0318 |
| (sum) |  | 233 | 1 |  | 220 | 1 |

The predicted toxicity nodes (e.g. “Toxicity to algae (level)”) were initially modelled by categorical nodes with 5 states, ranging from very low toxicity (>100 mg/L) to very high toxicity (< 0.01 mg/L). Note that increasing toxicity level corresponds to decreasing concentration of the substance (Table 2). The discretization of continuous toxicity was based on the classification and labelling of toxicity levels in the Globally Harmonized System (OECD, 2001), which has four toxicity levels with interval boundaries 1, 10, and 100 mg/L. To increase the resolution of the highest toxicity levels, we selected 5 intervals with the boundaries 0.01, 0.5, 5 and 100 mg/L.

The integration of the four lines of evidence was based on these 5-state categorical nodes. The toxicity interval nodes (e.g. “Toxicity to algae (interval)”) were added to provide a link to the continuous nodes for input values (e.g. “Toxicity to algae (value 1)”). Since interval nodes must have intervals in increasing order, the order of the toxicity intervals was reversed compared to the order of toxicity states. It was not strictly necessary to keep both the categorical and the interval version of the same toxicity node (e.g. toxicity to algae), but we found that it facilitated the interpretation and helped avoid confusion regarding the order of toxicity intervals vs. toxicity levels. An additional benefit of the interval nodes was to enable the extraction of a single expected value ("mean value") from a toxicity node, which could for example facilitate the comparison of posterior probability distributions of toxicity nodes for different substances. For this purpose, the toxicity interval nodes were given two additional extreme states (Table 3) with very low probability. The extreme states served as buffers for obtaining reasonable calculations of expected values. In this paper, for simplicity, we will focus on results for the 5-state categorical toxicity nodes.

**Table 3.**
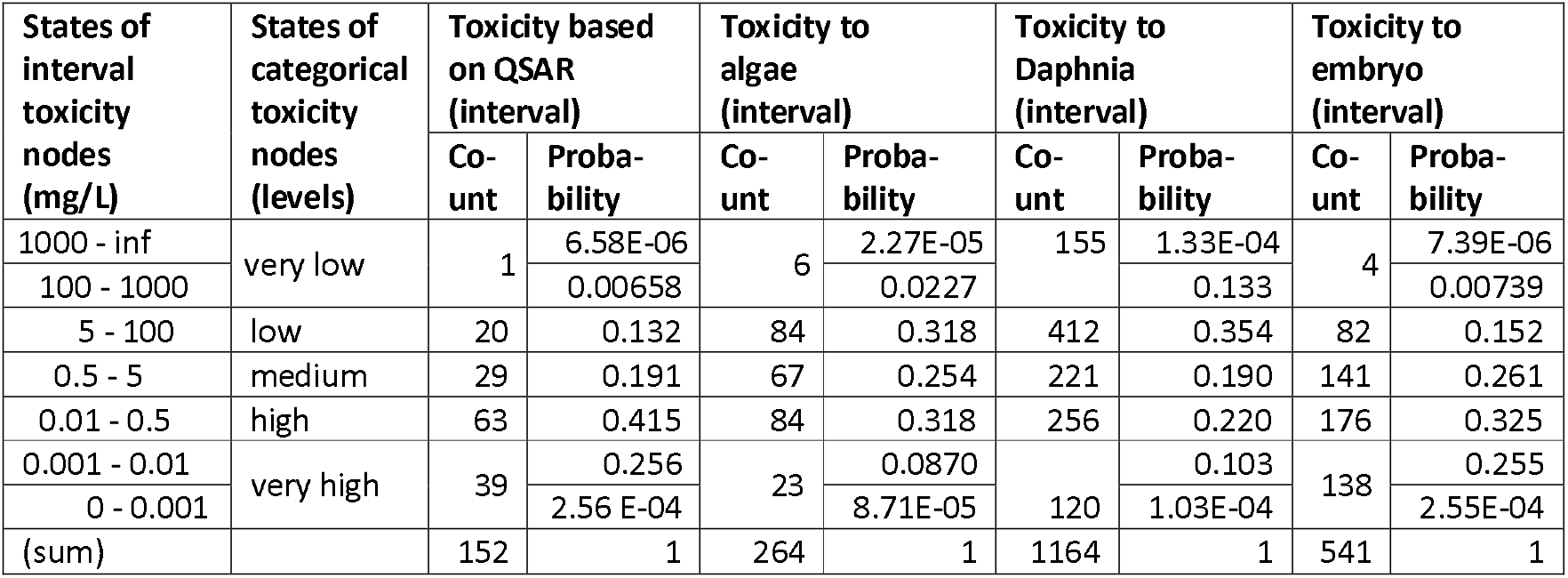
Prior probabilities for toxicity interval nodes (QSAR, algae, Daphnia and embryo), based on counts of values in our dataset. For fish, the toxicity values represent LC50 (the concentration resulting in a lethal effect to 50% of the tested population). For algae and Daphnia, the toxicity values represent EC50 (the concentration resulting in a given sublethal effect to 50% of the tested population). Note that the two most extreme intervals of the interval toxicity nodes (e.g. “100 - 1000” and “1000 - inf”) are merged into one categorical state (“very low”). The probabilities of the two intervals “100 - 1000” and “1000 - Inf” were set to respectively 99.9% and 0.01% of the proportion of counts in toxicity level “very low”. The probabilities of the intervals “0.001 - 0.01” and “0 - 0.001” were calculated correspondingly from the of count of values in toxicity level “very high”.

The continuous input value nodes (Figure 1) allowed users to provide their toxicity data as original values (EC50 or LC50) instead of discretized states. The function of the continuous nodes is to create a likelihood on the interval nodes using multiple findings as well as support by exact values. Although the continuous value nodes serve as input nodes for this BN, they are defined as child nodes of toxicity interval nodes (Figure 1). The reasoning is that the observed toxicity values (e.g. “Toxicity to algae (value 1)”) for a given substance are realisations of an inherent true toxicity value for this substance, which is unknown but can be modelled by the probability distribution of the toxicity interval node (“Toxicity to algae (interval)”.

The BN has multiple continuous input nodes for each endpoint (algae, daphnids and fish embryo) (Figure 1). The purpose was to let the BN model (1) retain as much details as possible from the input values, (2) account for variability in the provided data (i.e. variation among input values), and (3) account for inherent uncertainty in the toxicity values (e.g. a toxicity value closer to an interval boundary would have higher probability of being assigned to the next interval).

In the current version of the web interface, there are ten continuous input nodes for each of these endpoints, assuming that this is a reasonable maximum number of toxicity values available for a chemical substance. (In this dataset, the number of observations per substance exceeds ten for only 3%, 19% and 7% of the substances for algae, *Daphnia* and fish embryos, respectively). If needed, the number of continuous input nodes can easily be increased without affecting the model prediction. The unused input nodes will be so-called barren variables (Kjærulff and Madsen, 2008), which will be updated by evidence in their sibling nodes but will not themselves influence other nodes.

#### 2.3.2 Prior probabilities

Prior probability distributions were defined for all root nodes, i.e. all nodes with no parent nodes in the graph (Table 1). For all root nodes of interval type, the prior probability distribution was set equal to the frequency distribution of our dataset. This approach was used for prior probabilities of the following interval-type nodes: Molecular weight (Table S.2), Hydrophobicity (Table S.4), Toxicity based on QSAR (Table S.7), Toxicity to algae (Table S.13), Toxicity to *Daphnia* (Table S.14), Ratio toxicity *Daphnia* / algae (Table S.15) and Toxicity to embryo (Table S.18). For example, for the node “Molecular weight (interval)”, 231 substances had low molecular weight (<600 g/mol) while only 2 substances had high molecular weight (> 600 g/mol); the resulting prior probability distribution across these two states was 99.14% and 0.86% (Table 2).

For the toxicity interval nodes (QSAR, algae, *Daphnia* and embryo), the probabilities of states were also based on the counts of values. However, because of low numbers of values in the most extreme intervals, the counts were merged for the two intervals “100 - 1000” and “1000 - inf”, and the probabilities of these intervals were set to respectively 99.9% and 0.01% of the proportion of values in toxicity level “very low” (Table 3). The probability of the intervals “0.001 - 0.01” and “0 - 0.001”, likewise, were set to respectively 99.9% and 0.01% of the proportion of values in toxicity level “very high”.

One root node was of categorical type: “Chemical category” (Table 1). Since this was also an input node to be instantiated by the user, the prior distribution was not of importance, and was set to uniform distribution (Table S.11)

### 2.3.3 Conditional probability tables and density functions

The conditional probability tables (CPTs) of the BN were parametrized by four main approaches, illustrated by the four examples in Figure 2: (1) expert knowledge (Figure 2a), (2) frequency distributions derived from our dataset (Figure 2b), (3) statistical considerations (Figure 2c), and a rule for combining the lines of evidence (Figure 2d).

**Figure 2.**
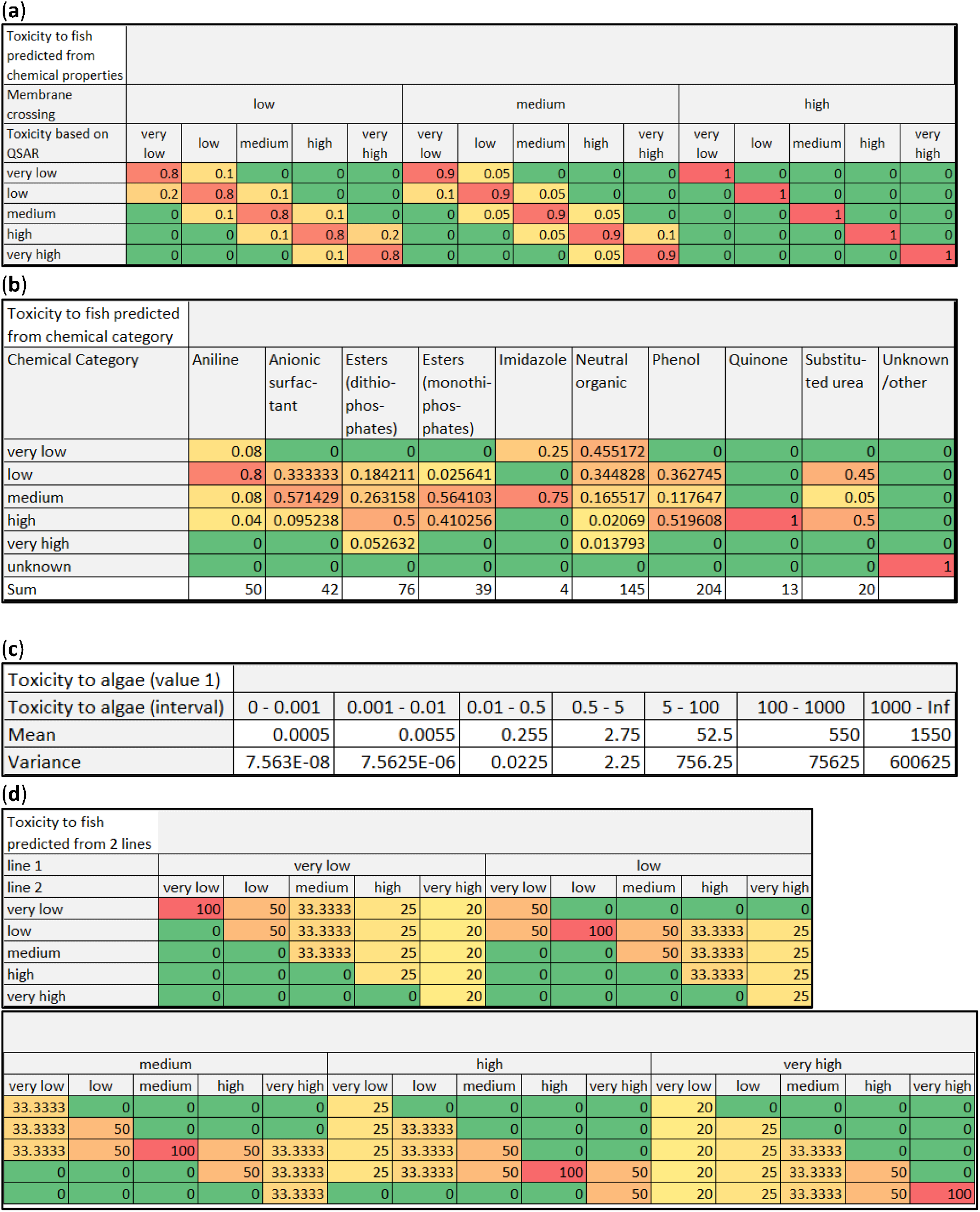
Examples of conditional probability tables (CPTs) and density functions (CDFs) for selected nodes, illustrating the different approaches used for parameterisation of the conditional dependencies. The colour code represents the scale from green (zero) to red (1 or 100%). **(a)** CPTs for the node “Membrane crossing”: the probability of a substance crossing a biological membrane based on its physical properties (molecular weight and hydrophobicity). Probabilities are based on expert judgement. **(b)** Extract of the CPT for the node “Toxicity to fish predicted from chemical category”. The probabilities are derived from frequency distributions in our dataset. The supplementary data contains the full CPT for all 42 chemical categories (Table S1.11). **(c)** CDF for the node “Toxicity to algae (value 1)”: mean and variance of observations for each toxicity interval (Table 1). The values are set based on a statistical rule (see Appendix A). **(d)** CPT for the node “Toxicity to fish predicted from all evidence”: combination rule representing equal weighting of the four lines of evidence. This table shows combination of two of the four lines, while the combination of all four lines (Table S1.21) is described in the Supplementary material.

Expert knowledge (A. Lillicrap) was used for CPT of the nodes “Membrane crossing” (Table S.6), “Toxicity to fish predicted from chemical properties” (Table S.10) and “Toxicity to fish predicted from other taxa” (Table S.17). For “Membrane crossing”, the combination of low hydrophobicity and low molecular weight was assumed to result in respectively 0%, 25% and 75% probability of low, medium and high ability of membrane crossing, while the opposite combination of hydrophobicity and molecular weight was assigned the opposite probability distribution. The two combinations of high/low and low/high hydrophobicity and molecular weight were assigned a probability distribution of 25%, 50% and 25% for the three states of membrane crossing.

An example is shown for “Toxicity to fish predicted from chemical properties” (Figure 2a). When a substance’s ability to cross a biological membrane is high, the predicted toxicity to fish based on chemical properties is equal to the acute fish toxicity modelled by QSAR. When the membrane crossing ability is low, we assigned only 80% chance that the predicted acute toxicity corresponds to the modelled QSAR result.

The reasoning behind the CPT of “Toxicity to fish predicted from other taxa” is explained for all 50 combinations of the parent states by Lillicrap et al. (2019). In brief, if the ratio of the average toxicity values to *Daphnia* vs. algae is between 0.5 and 2, it can be assumed that the chemical has a similar mode of action for the two taxa, and that the chemical will affect fish in a similar way. Therefore, the CPT converts the EC50 values from algae and *Daphnia* to LC50 values for fish with high precision (a narrow distribution), corresponding to assigning a high weight to these pieces of evidence in a WOE approach. Moreover, narrower probability distributions (more weight) have been assigned to the evidence from *Daphnia* than from algae, assuming that the closer phylogenetic relationship of fish with invertebrates than with plants makes their responses to a chemical more similar. Conversely, a ratio of >2 (or <0.5) indicates that the chemical has a species-specific mode of action, which affects *Daphnia* more strongly than than algae (or vice versa). In these cases, we have less confidence in extrapolating the toxicity data from *Daphnia* or algae to fish, and have therefore set wider probability distributions in this part of the CPT (corresponding to lower weighting in a WOE approach).

CPTs for the node “Toxicity to fish predicted from chemical category”, as well as for all continuous nodes (identified in Table 1), were created using frequency distributions derived from our dataset. The CPT for “Toxicity to fish predicted from chemical category” is illustrated in Figure 2b. For each chemical category, the values represent the proportion of juvenile fish LC50 values in each of the given toxicity levels. For example, the 50 observations in the chemical category “Aniline” comprised 4 observations of the toxicity level “very low”, 44 observations of “low”, 4 observations of “medium and two observations of “high”, from altogether 13 chemical substances. The number of observations per chemical category was sometimes quite low (e.g. Imidazole, Figure 2b). To optimise the use of the available data, multiple toxicity observations for the same substance were counted independently. For chemical categories with a low total number of observations (Table S.11), the proportions would not properly represent a probability distribution (Marcot, 2017), therefore the model predictions will be less reliable for these chemical categories. In addition, a chemical category “unknown / other” can be selected. When this state is chosen, this line of evidence is excluded from the final predicted toxicity node.

For the continuous nodes, the conditional probability distributions were quantified by a conditional density function (CDF), defined by a Gaussian distribution with mean and variance specified for each interval of the parent node (i.e. the corresponding interval node, see Table 1). For the physico-chemical nodes of Line 1, mean values were calculated to reflect the chemicals in our dataset. Variances were calculated according to a pre-defined coefficient of variation (CV; standard deviation divided by the mean). We chose not to let the variance be calculated directly from our dataset, as the variance would have been sensitive to the number of observations in each interval. Instead, we assumed low CV of physico-chemical properties of substances such as molecular weight (10% CV) and hydrophobicity (5% CV). For example, for the node “Molecular weight (value)”, the CV was set to 5% of the mean value. In the CDF for this node (Table S.3), the two intervals (<600 g/mol and >600 g/mol) had mean values of 179 and 200,380, respectively, calculated from the respective counts of 231 and 2 observations (see Table 2).

For all continuous nodes representing toxicity values (based on QSAR, algae, *Daphnia* or embryo), the density functions were defined in the same way. The mean for each interval of the parent node was calculated as the midpoint of the interval, except for the extreme interval “1000- inf”; here the mean was set to 1550, which was equal to the lower boundary plus the midpoint of the previous interval. For each interval, the variance was set so that a 90% of a Gaussian distribution was within the interval while 5% of the distribution was in either of the neighbour intervals (see Appendix A). The resulting variances (Figure 2c, Table S.8) correspond to ca. 30% CV of the upper interval boundary, for all intervals except the highest (inf). For toxicity data from standard assays performed in various laboratories, CV up to 30% is a reasonable assumption (Rawlings et al., 2019).

The CPT of the final child node, “Toxicity to fish predicted from all evidence”, was defined by a combination rule to ensure that all four lines of evidence were given equal weight. Following the recommendation of Marcot (2017) to obtain more tractable CPT dimensionality, we first set the probabilities for combination of two lines (Figure 2d). The combination of lines 1+2 and of lines 3+4 each resulted in two intermediate child nodes, which were subsequently combined with the same rule (Figure 2d) to produce a new “grandchild” node. The two intermediate nodes were then absorbed, which resulted in a CPT combining the four “grandparents” nodes with the one “grandchild” node, as described in the Supplementary material (Table S.21).

### 2.4 Model assessment

#### 2.4.1 Sensitivity analysis

The sensitivity of the target node ("Toxicity to fish predicted from all evidence") to the different lines of evidence was analysed by two methods: Parameter sensitivity analysis and value-of-information analysis.

The parameter sensitivity analysis measures the functional relationships between a parameter (i.e., a probability value in the CPT of a node) and the posterior probability of a given state of the target node (Chan and Darwiche, 2002). Due to the structure of the evidence (no downstream evidence relative to the target variable), the functional relationship is linear (Coupé and van der Gaag, 2002). The sensitivity analysis was run relative to all states (toxicity levels) of the target node, under the default scenario of no evidence (no data used to run the model).

Value-of-information (VoI) analysis can quantify the potential benefit of additional information in the face of uncertainty (Keisler et al., 2014), and can aid the decision on allocation of resources between obtaining new information and improving management actions (Mäntyniemi et al., 2009). The method quantifies the VoI as reduction of entropy (a measure of uncertainty) for a given node in isolation, averaging the values of other nodes. The VoI analysis was run under different scenarios of evidence (information applied to the model): the default scenario of no evidence, as well as data for three example substances, using either all available data either for the first three lines of evidence (excluding embryo data) or for all four lines of evidence. The purpose was to assess the potential benefit of including embryo data as a line of evidence.

#### 2.4.2 Model evaluation

The BN model performance and uncertainty was evaluated by several of the metrics recommended (Marcot, 2012). Model performance was assessed by running with input data from our dataset and comparing the outcome (i.e. the predicted acute toxicity of selected chemical substances to juvenile fish), to the observed toxicity of the same substances to this endpoint. The model validation should ideally be performed with an independent dataset that had not already been used for parametrization of the model. However, additional fish embryo toxicity data are notwidely available, and our dataset represents the largest collection of OECD 236 FET data in the public domain to our knowledge. Instead, we applied four different criteria for selecting subsets of our dataset for valiation in order to consider the robustness of the model from different perspectives.

- Subset 1: all chemical substances containing at least one observation of juvenile fish toxicity (AFT) data (number of substances: 159), which was the minimum requirement for comparison with the model prediction. For substances missing one or more input values (e.g. toxicity to algae), the prediction would be based on the prior probabilities of this node. The exception was Line 2 (toxicity of the chemical category), which allowed for the state “unknown”.
- Subset 2: only the chemical categories that were represented with minimum 10 different chemical substances in our dataset (number of substances: 106). Using Phenol as an example, the conditional probabilities of a chemical category in the node “Toxicity to fish predicted from chemical category” (Figure 2b), was calculated as the frequency distribution of observed toxicity to juvenile fish for all substances in the category Phenol. When the predicted toxicity of a substance belonging to the category Phenol was assessed (e.g. Triclosan), the observed toxicity values of Triclosan used for comparison have also been used in the CPT. For chemical categories containing few substances in our dataset, there will be a high overlap between the data used for parametrization and for validation. For chemical categories with 10 or more substances, however, the data used for validation for one of these substances will constitute only a small proportion of the frequency distribution of the CPT. Therefore, a validation based on such subsets will be more independent of the parametrization data.
- Subset 3: all substances with complete cases for input nodes (number of substances: 77). This means that our dataset contained minimum one value for each node defined as Input node (Table 1): Molecular weight, Hydrophobicity, QSAR, Toxicity to algae, Toxicity to *Daphnia* and Toxicity to embryo, as well as Toxicity to juvenile. For this subset, the validation would be less influenced by the prior probabilities (Tables 2-3), since evidence would be available to update the probabilities of all input nodes.
- Subset 4: cases with a minimum of 3 observations for both embryo and juvenile toxicity data (number of substances: 20). The reasoning was that these two nodes are the most crucial for the purpose of the model. A higher number of observations would make the model predictions more precise and reduce bias due to variability in these types of data.

For the purpose of comparing predicted and observed probability distributions, it is common to define the state with the highest probability as the “correct” state (Marcot, 2012). We followed this practice and defined the toxicity interval with the highest probability as the “correct” predicted or observed state. This way, we could calculate the number of correct predictions for each toxicity interval and for each data subset. This practice does not, however, account for the whole probability distribution. More advanced methods for model validation consider the probability distributions of predicted and observed variables will be investigated in further work with this model.

## 3 Web interface to the BN model

A web-based user interface to the BN model for demonstration purposes has been published on HUGIN‘s web portal for BN examples, http://demo.hugin.com/example/FET. Our model is the first example of a BN within the field of ecotoxicology and ecological risk assessent on this platform. The web interface to this preliminary BN version will give researchers and other potential users the opportunity to provide feedback for improving the model. In the longer term, our intention is to further develop this model into a more comprehensive online tool which can be useful for regulatory authorities and chemical industries wanting to submit fish embryo toxicity data in place of acute fish toxicity data.

Here we briefly describe the two interactive pages of the web interface: “Enter values” (Supplementary information, Figure S.1) and “Results” (Figude S.2). In the tab “Enter values”, a user can insert the requested information for any chemical substance for which they want to predict the acute toxicity to juvenile fish. The user should provide the requested information as follows, for the four lines of evidence:

- Line 1: Hydrophobicity (log K_OW_), Molecular weight (g/mol), Toxicity based on QSAR (mg/L). The user must enter one continuous value for each node.
- Line 2: Chemical category. The user must select the category from a drop-down list, which includes the state “Unknown / other”. More chemical categories can be added upon request from users.
- Line 3: Toxicity of the substance to other taxa - algae (OECD 201) and *Daphnia* (OECD 202). The user should first select the number of values (up to ten) for each taxon, then enter the EC_50_ values (mg/L).
- Line 4: Toxicity of the substance to fish embryos (OECD 236). The user should first select the number of values (up to ten), then enter the LC_50_ values (mg/L).

The tab “Enter values” contains the buttons “Compute”, which will carry out the calculation based on these input values, and “Load Case”, which will load a built-in example: the substance carbamazepine.

The tab “Results” displays the posterior probability distributions across the five toxicity levels for all four lines of evidence (“Toxicity to fish predicted from chemical properties”, “Toxicity to fish predicted from chemical category”, “Toxicity to fish predicted from other taxa", and Toxicity to embryo (level)”), as well as for the final child node (“Toxicity to fish predicted from all evidence”) (See Supplementary information). Two buttons generate tables as pop-up windows. The button “View input values” provides a table of the entered values, as well as the calculated ratio of toxicity to *Daphnia* vs. algae (Table 4a). This table also identifies the most sensitive endpoint (algae, *Daphnia* or fish embryo), which is relevant for the application of these results for regulatory risk assessment (Lillicrap et al., 2019). The button “View output values” provides a table with the posterior probability distributions for the five selected nodes mentioned above (Table 4b). The “Results” tab also provides conclusive statements based on the calculated values, such as “The toxicity level of carbamazepine to juvenile fish is most likely low (52.28% probability)”.

**Table 4.** Example of tables of **(a)** input values and **(b)** output values generated by the web interface, for the substance carbamazepine (see Figure 3a). **(a)** The input table includes values that are derived directly from the input values (without calculation of probabilities): mean toxicity values for algae, Daphnia and embryo, and the most sensitive endpoint of those three. **(b)** The output table contains the predicted toxicity for the four lines of evidence, as well the combined predicted toxicity for juvenile fish (as probabilitiy distributions across the five toxicity levels).

| Substance | Node | Value | Unit |
| --- | --- | --- | --- |
| carbamazepine | Molecular weight (value) | 236.28 | g/mol |
| carbamazepine | Hydrophobicity (value) | 2.45 | log $k_{ow}$ |
| carbamazepine | Toxicity based on QSAR (value) | 41.33 | mg/L |
| carbamazepine | Chemical category | Substituted urea |  |
| carbamazepine | Daphnia / algae | 0.98 |  |
| carbamazepine | Toxicity to embryo: value 1 | 148 | mg/L |
| carbamazepine | Toxicity to embryo: value 2 | 158 | mg/L |
| carbamazepine | Toxicity to embryo: value 3 | 150 | mg/L |
| carbamazepine | Toxicity to embryo: value 4 | 156 | mg/L |
| carbamazepine | Toxicity to embryo: value 5 | 157 | mg/L |
| carbamazepine | Toxicity to algae: value 1 | 163.77 | mg/L |
| carbamazepine | Toxicity to algae: value 2 | 49.4 | mg/L |
| carbamazepine | Toxicity to Daphnia: value 1 | 97.815 | mg/L |
| carbamazepine | Toxicity to Daphnia: value 2 | 111 | mg/L |
| carbamazepine | Toxicity to algae: mean | 108.4 | mg/L |
| carbamazepine | Toxicity to Daphnia: mean | 104.4 | mg/L |
| carbamazepine | Toxicity to embryo: mean | 153.3 | mg/L |
| carbamazepine | Most sensitive endpoint | Daphnia |  |

(b)
| Substance | Node | Toxicity level | Probability |
| --- | --- | --- | --- |
| carbamazepine | Toxicity to fish predicted from chemical properties | very low | 0.02 |
| carbamazepine | Toxicity to fish predicted from chemical properties | low | 0.96 |
| carbamazepine | Toxicity to fish predicted from chemical properties | medium | 0.01 |
| carbamazepine | Toxicity to fish predicted from chemical properties | high | 0 |
| carbamazepine | Toxicity to fish predicted from chemical properties | very high | 0 |
| carbamazepine | Toxicity to fish predicted from chemical category | very low | 0 |
| carbamazepine | Toxicity to fish predicted from chemical category | low | 0.45 |
| carbamazepine | Toxicity to fish predicted from chemical category | medium | 0.05 |
| carbamazepine | Toxicity to fish predicted from chemical category | high | 0.5 |
| carbamazepine | Toxicity to fish predicted from chemical category | very high | 0 |
| carbamazepine | Toxicity to fish predicted from other taxa | very low | 0 |
| carbamazepine | Toxicity to fish predicted from other taxa | low | 0.39 |
| carbamazepine | Toxicity to fish predicted from other taxa | medium | 0.5 |
| carbamazepine | Toxicity to fish predicted from other taxa | high | 0.11 |
| carbamazepine | Toxicity to fish predicted from other taxa | very high | 0 |
| carbamazepine | Toxicity to embryo (level) | very low | 0 |
| carbamazepine | Toxicity to embryo (level) | low | 1 |
| carbamazepine | Toxicity to embryo (level) | medium | 0 |
| carbamazepine | Toxicity to embryo (level) | high | 0 |
| carbamazepine | Toxicity to embryo (level) | very high | 0 |
| carbamazepine | Toxicity to fish predicted from all evidence | very low | 0 |
| carbamazepine | Toxicity to fish predicted from all evidence | low | 0.29 |
| carbamazepine | Toxicity to fish predicted from all evidence | medium | 0.52 |
| carbamazepine | Toxicity to fish predicted from all evidence | high | 0.14 |
| carbamazepine | Toxicity to fish predicted from all evidence | very high | 0.05 |

## 4 Results and Discussion

### 4.1 Examples of BN model predictions

Examples of model predictions for three selected substances are presented in Figure 3. All examples are from subset 4 (see section 2.4.2), which has minimum three toxicity data for both embryo and juvenile fish. The three examples represent three different levels of observed toxicity to juvenile fish: low, medium and high.

**Figure 3.**
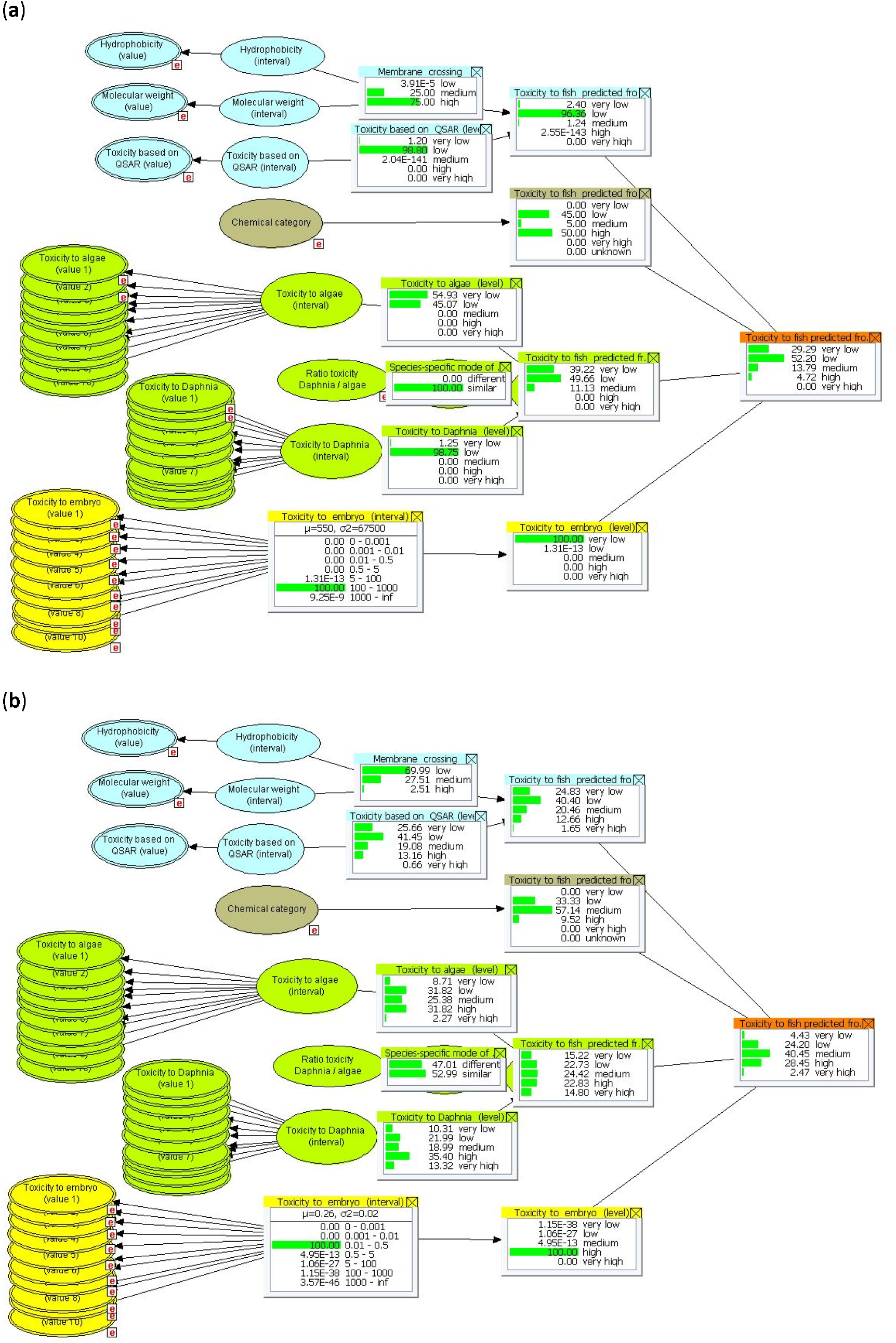
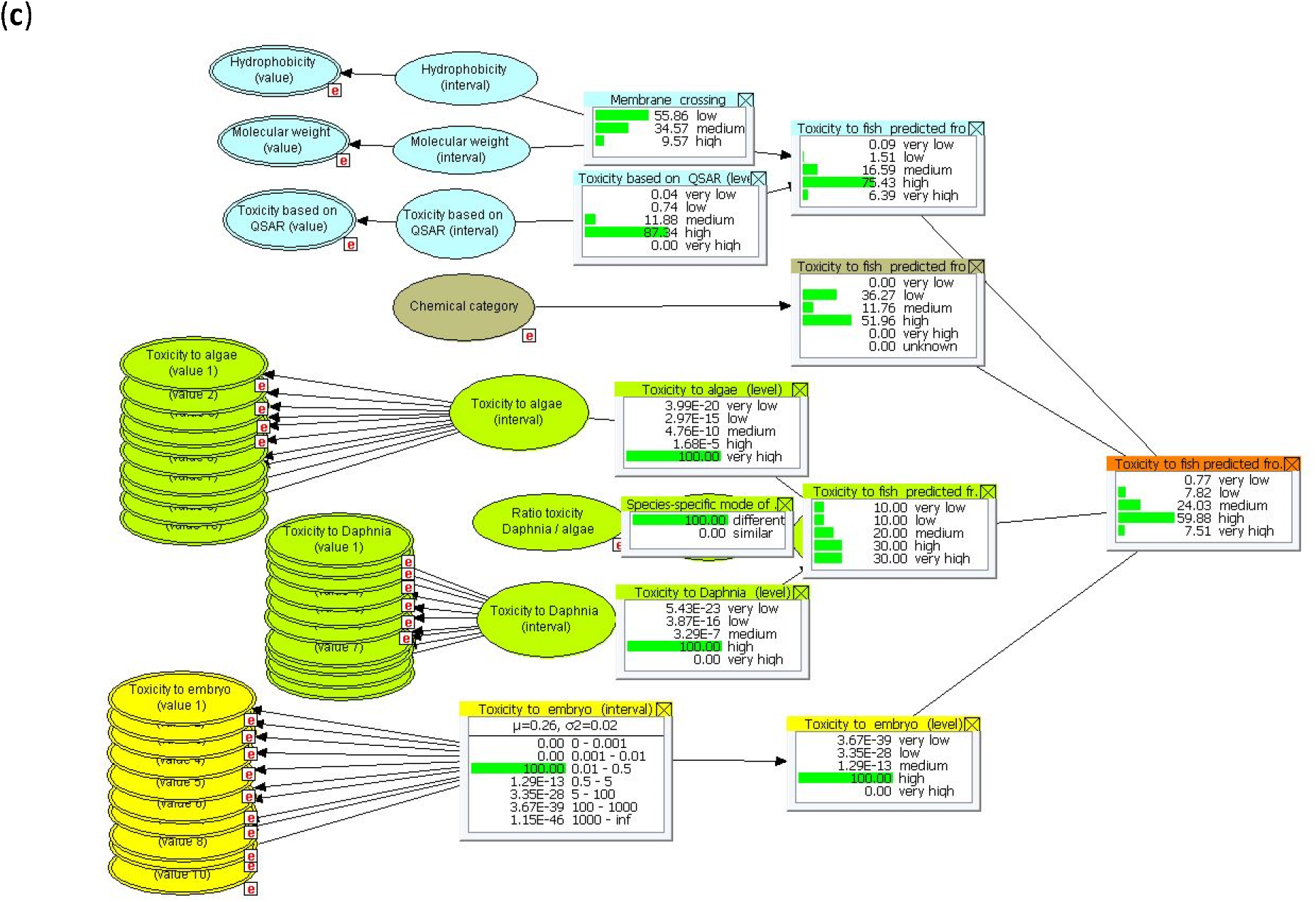
Examples of BN model predictions for three selected substances: **(a)** Carbamazepine, **(b)** Tetradecyl sulfate, **(c)** Triclosan. Monitor windows with posterior probability distributions are shown for selected nodes in each line of evidence (please see Figure 1 for complete node lables and arrows). The red “e” boxes indicate input nodes that have been instantiated with evidence. The full set of input values and selected output values for example (a) are given in Table 4.

The first example (Figure 3a), Carbamazepine (an antiepileptic drug in the chemical category Substituted urea), is also used as a built-in example in the online demonstration model (Figures S.1-S.2). The observed toxicity to juvenile fish, to which the predicted toxicity will be compared, is 100% in the low toxicity interval (four observations; not shown). The node “Toxicity to embryo (level)” was almost 100% “very low”, while the toxicity based on QSAR, algae and *Daphnia* was “low” to “very low”. The information from the chemical category, however, indicated a 50% probability of high toxicity to juvenile fish. This line of evidence contributed to higher predicted toxicity to juvenile fish (26% probability of medium or higher toxicity) than the other three lines. After combination of the four lines, the most likely state was low toxicity to juvenile fish (52% probability), which was consistent with the measured toxicity interval, although with higher uncertainty. In this case, information on toxicity to embryo (very low) alone would have underestimated the risk to juvenile fish (low), while the embryo data in combination with the other three lines of evidence resulted in a more accurate prediction on toxicity to juvenile fish.

The second example (Figure 3b) was Tetradecyl sulfate, a drug in the chemical category Anionic surfactant. Our dataset contained three LC50 values from AFT assays (toxicity to juvenile fish), all of which fell into the medium toxicity interval. The node “Toxicity to embryo (level)” predicted a 100% probability the high toxicity interval, while the predictions from the other lines of evidence were centered around low-to-medium toxicity interval. The resulting predicted toxicity to juvenile fish had highest probability (41%) of the medium state, which was consistent with the measured toxicity. In this case, information on toxicity to embryo alone would have overestimated the risk to juvenile fish, while the embryo data in combination with the other lines of evidence again resulted in a more accurate prediction, although with lower precision than the original AFT data.

The third example (Figure 3c), Triclosan (an antimicrobial agent in the chemical category Phenol), had six LC50 values for juvenile fish in the high toxicity interval. The data based on embryo, *Daphnia* and QSAR showed the same toxicity level, while toxicity to algae was very high. Toxicity based on the chemical category, on the other hand, was almost 50% likely to be medium or lower. When the four lines of evidence were combined, the most probable toxicity level was high (60%), which was again consistent with the observations.

### 4.2 Sensitivity analysis

The parameter sensitivity analysis (Table 5) showed that under the default evidence scenario of no evidence, the stgraphate “very high” of the target node (“Toxicity to fish predicted from all evidence”) as expected was the most sensitive to changes in single parameters in the probability tables of the parent nodes. The target node was most sensitive to parameters of the nodes “Toxicity to embryo (interval)” and “Toxicity based on QSAR (interval)”. The sensitivity value (i.e., the slope of the posterior probabilities of the target node to these parameters) was 0.16 when averaged across all states of the target node. The highest sensitivity was found for the state “high toxicity” of the target node, with slopes 0.21 (QSAR) and 0.19 (embryo) and. These nodes are root nodes (Table 1), with probability tables containing prior probabilities (Tables S.6 and S.18). The analysis shows that the prior probabilities of these nodes may have a strong influence on the prediction, which implies that the approach used for setting priors (Appendix A) may need to be refined.

**Table 5.** Parameter sensitivity analysis for the scenario default scenario of no evidence. The target node is “Toxicity to fish predicted from all evidence”. The sensitivity values represent the slope of the linear relationship between a change in one parameter of the CPT for a given node and the resulting change in the probability of each state of the target node. The maximum and minimum values are extracted across all states of the parent nodes. For example, consider the parent node “Toxicity to embryo (interval)”. If the probability of each state separately is changed by +0.1, then the resulting change in the posterior probability of predicted toxicity being “very low” ranges from 0.017 to −0.0072. In the column “all states”, max an min valus are averaged across the five states of the parent node. The rows are ordered by decreasing sensitivity in the columns “all states”, where sensitivity values are averaged across the five states of the target node.

| Target node | all states |  | very low |  | low |  | medium |  | high |  | very high |  |
| --- | --- | --- | --- | --- | --- | --- | --- | --- | --- | --- | --- | --- |
| Parent node | max | min | max | min | max | min | max | min | max | min | max | min |
| Toxicity to embryo (interval) | 0.162 | -0.085 | 0.170 | -0.072 | 0.150 | -0.156 | 0.162 | -0.114 | 0.194 | -0.075 | 0.134 | -0.008 |
| Toxicity based on QSAR (interval) | 0.160 | -0.083 | 0.159 | -0.069 | 0.140 | -0.172 | 0.163 | -0.099 | 0.207 | -0.070 | 0.130 | -0.005 |
| Chemical category | 0.082 | -0.058 | 0.081 | -0.047 | 0.119 | -0.099 | 0.070 | -0.070 | 0.118 | -0.070 | 0.021 | -0.006 |
| Toxicity to Daphnia (interval) | 0.039 | -0.037 | 0.044 | -0.028 | 0.063 | -0.048 | 0.025 | -0.040 | 0.043 | -0.057 | 0.018 | -0.012 |
| Toxicity to algae (interval) | 0.023 | -0.019 | 0.025 | -0.016 | 0.026 | -0.029 | 0.020 | -0.022 | 0.027 | -0.023 | 0.017 | -0.005 |
| Tox. pred. from chemical category | 0.018 | -0.016 | 0.020 | -0.020 | 0.022 | -0.021 | 0.021 | -0.021 | 0.017 | -0.016 | 0.010 | -0.001 |
| Tox. pred. from chemical properties | 0.012 | -0.009 | 0.017 | -0.010 | 0.014 | -0.019 | 0.017 | -0.007 | 0.009 | -0.006 | 0.004 | -0.002 |
| Tox. pred. from other taxa | 0.010 | -0.007 | 0.013 | -0.005 | 0.012 | -0.009 | 0.010 | -0.008 | 0.009 | -0.010 | 0.006 | -0.003 |
| Tox. pred. from all evidence | 0.009 | -0.006 | 0.008 | -0.006 | 0.010 | -0.008 | 0.010 | -0.008 | 0.010 | -0.006 | 0.006 | -0.001 |
| Ratio toxicity Daphnia / algae | 0.003 | -0.003 | 0.001 | -0.001 | 0.005 | -0.006 | 0.002 | -0.002 | 0.005 | -0.005 | 0.001 | -0.001 |
| Membrane crossing | 0.001 | -0.001 | 0.001 | -0.001 | 0.002 | -0.002 | 0.001 | -0.001 | 0.001 | -0.001 | 0.001 | -0.001 |
| Hydrophobicity (interval) | 0.001 | -0.001 | 0.001 | -0.001 | 0.001 | -0.001 | 0.001 | -0.001 | 0.001 | -0.001 | 0.001 | -0.001 |
| Molecular weight (interval) | 0.001 | -0.001 | 0.001 | -0.001 | 0.001 | -0.001 | 0.001 | -0.001 | 0.001 | -0.001 | 0.001 | -0.001 |

The third most influential parameters were for “Chemical category”, to which the sensitivity of the target node was 0.08 on average and 0.12 for the state “high toxicity”. The prior probability table of this root node has a uniform distribution, assuming that for a new chemical substance, all chemical categories are equally likely. But considering the high influence of this probability table on the target node, we could consider a more informative prior probability distribution, e.g. which better reflects the frequency of the different chemical categories in our dataset or other larger datasets.

The fourth and fifth most influential parameters were for “Toxicity to *Daphnia* (interval)” and “Toxicity to algae (interval)”. The influence of these nodes on the target node were weakened by the additional node “species-specific mode of action”, which is meant to introduce uncertainty related to the extrapolation of results from plants and invertebrates to fish. The target node is slightly more sensitive to changes in parameters for *Daphnia* than for algae. This is consistent with the closer phylogenetic relationship of fish with invertebrates than with plants, which we tried to account for in the CPT (Table S.17) (Lillicrap et al., 2019).

In general, this sensitivity analysis reflects the strongest influence of parameters in the parts of the model that are most directly based on data (such as Toxicity based on QSAR and Toxicity to embryo), and weaker influence in lines of evidence that rely strongly on expert knowledge, such as Membrane crossing and Toxicity predicted from other taxa. Although these lines of evidence are also based on data (e.g. measured toxicity to algae and *Daphnia*), expert knowledge is applied in weighting and combining these pieces of evidence. Hence, there is a potential for making the model more sensitive by refining the CPTs that are currently based on expert knowledge (e.g. Figure 2a). The sensitivity analysis also shows that the posterior distribution of the target node is robust to changes in any single parameter of the BN as all sensitivity values are less than one.

The value-of-information (VOI) analysis (Table 6) showed that under the default scenario of no evidence, all four lines of evidence contribute almost equally and relatively little to entropy reduction (4-7% of the entropy of the target node). Line of evidence 3 (other taxa) contribute slightly more than the other lines. This can reflect the fact that the conditional probabilities of “Toxicity predicted from other taxa” (Table S.17) can be either highly correlated with target node or be non-informative (flat distribution), depending on the input values (node “Ratio toxicity *Daphnia* / algae” (Lillicrap et al., 2019).

**Table 6.** Value-of-information analysis run for five evidence scenarios: no evidence (default) and three example substances with or without data on toxicity to embryo (Line of evidence 4). The row “Entropy of target node” contains the entropy (uncertainty) of the node “Toxicity to fish predicted from all evidence” under the given evidence scenario. The four subsequent rows contain the maximal entropy reduction that can provided by additional information in each of the four parent nodes, considered separately while averaging the values of the other nodes.

|  | No evidence | Carbamazepine |  | Tetradecyl sulfate |  | Triclosan |  |
| --- | --- | --- | --- | --- | --- | --- | --- |
|  |  | Lines 1-3 | Lines 1-4 | Lines 1-3 | Lines 1-4 | Lines 1-3 | Lines 1-4 |
| Entropy of target node | 1.36 | 1.13 | 0.93 | 1.29 | 1.30 | 1.31 | 1.08 |
| 1) Chemical properties | 0.05 | 0.01 | 0.01 | 0.06 | 0.07 | 0.02 | 0.03 |
| 2) Chemical category | 0.05 | 0.07 | 0.06 | 0.03 | 0.02 | 0.04 | 0.07 |
| 3) Other taxa | 0.09 | 0.05 | 0.09 | 0.09 | 0.10 | 0.09 | 0.14 |
| 4) Embryo | 0.06 | 0.08 | 0 | 0.03 | 0 | 0.06 | 0 |

For scenarios where information is provided only for the lines of evidence 1-3 (i.e. excluding information on toxicity to embryos), additional information on embryo toxicity can reduce the uncertainty of the target node by 5-7% (Table 6). The relative importance of information on embryo toxicity varies among the three example substances, which suggests that the importance of this type of information cannot be generalised across substances. Under the scenario of full information (Lines 1-4), further information on embryo toxicity will not contribute to reduction in entropy. This result suggests that the information on embryo toxicity used in each example has already exerted a strong influence on the prediction, which is consistent with the outcome of the parameter sensitivity analysis (Table 5).

### 4.3 Model performance

The performance of the BN for the four different subsets of our dataset is reported in Table 7 and summarised in Figure 4. For each substance, the predicted juvenile toxicity state with the highest probability was compared with the most frequently observed juvenile toxicity state. The first three subsets, which have a number of substances ranging from 77 to 159, showed very similar results: The percentage of correctly predicted toxicity level was 69-71%, while the percentage of overestimated toxicity was 14-18% and the percentage of underestimated toxicity was 12-16%. The model typically overestimated toxicity when the observed toxicity was very low or low. Conversely, the few observed cases of high toxicity were sometimes underestimated as medium. The selected subsets of dataset were dominated by substances with very low to medium toxicity, which can explain the slightly higher proportion of cases with overestimated toxicity. If the model is applied to a new substance with high toxicity, the model is more likely to underestimate its toxicity as medium than to overestimate it as very high.

**Table 7.** Summary of BN model predictions for different subsets of our dataset. **(a)** Subset 1: All chemical substances with juvenile toxicity data available (minimum one value); no. of substances (n) = 159. **(b)** Subset 2: All substances from chemical categories with data from minimum 10 different substances; n = 106. **(c)** Subset 3: All complete cases, i.e. substances with minimum one value for each input node (see Table 1); n = 77. **(d)** Subset 4: Selected cases, i.e. substances with minimum three values for both embryo and juvenile toxicity data; n = 20.

| Predicted \ Observed | very low | low | medium | high | very high | sum |
| --- | --- | --- | --- | --- | --- | --- |
| very low | 26 | 16 | 0 | 0 | 0 | 42 |
| low | 2 | 42 | 10 | 0 | 0 | 54 |
| medium | 2 | 8 | 25 | 3 | 0 | 38 |
| high | 0 | 2 | 4 | 15 | 0 | 21 |
| very high | 0 | 0 | 1 | 2 | 1 | 4 |

**(b)**
| Predicted \ Observed | very low | low | medium | high | very high | sum |
| --- | --- | --- | --- | --- | --- | --- |
| very low | 23 | 12 | 0 | 0 | 0 | 35 |
| low | 1 | 35 | 5 | 0 | 0 | 41 |
| medium | 1 | 6 | 13 | 1 | 0 | 21 |
| high | 0 | 2 | 2 | 4 | 0 | 8 |
| very high | 0 | 0 | 1 | 0 | 0 | 1 |

**(c)**
| Predicted \ Observed | very low | low | medium | high | very high | sum |
| --- | --- | --- | --- | --- | --- | --- |
| very low | 10 | 6 | 0 | 0 | 0 | 16 |
| low | 1 | 28 | 4 | 0 | 0 | 33 |
| medium | 1 | 7 | 10 | 1 | 0 | 19 |
| high | 0 | 0 | 2 | 6 | 0 | 8 |
| very high | 0 | 0 | 0 | 1 | 0 | 1 |

**(d)**
| Predicted \ Observed | very low | low | medium | high | very high | sum |
| --- | --- | --- | --- | --- | --- | --- |
| very low | 3 | 0 | 0 | 0 | 0 | 3 |
| low | 0 | 7 | 0 | 0 | 0 | 7 |
| medium | 0 | 3 | 4 | 0 | 0 | 7 |
| high | 0 | 0 | 1 | 2 | 0 | 3 |
| very high | 0 | 0 | 0 | 0 | 0 | 0 |

**Figure 4.**
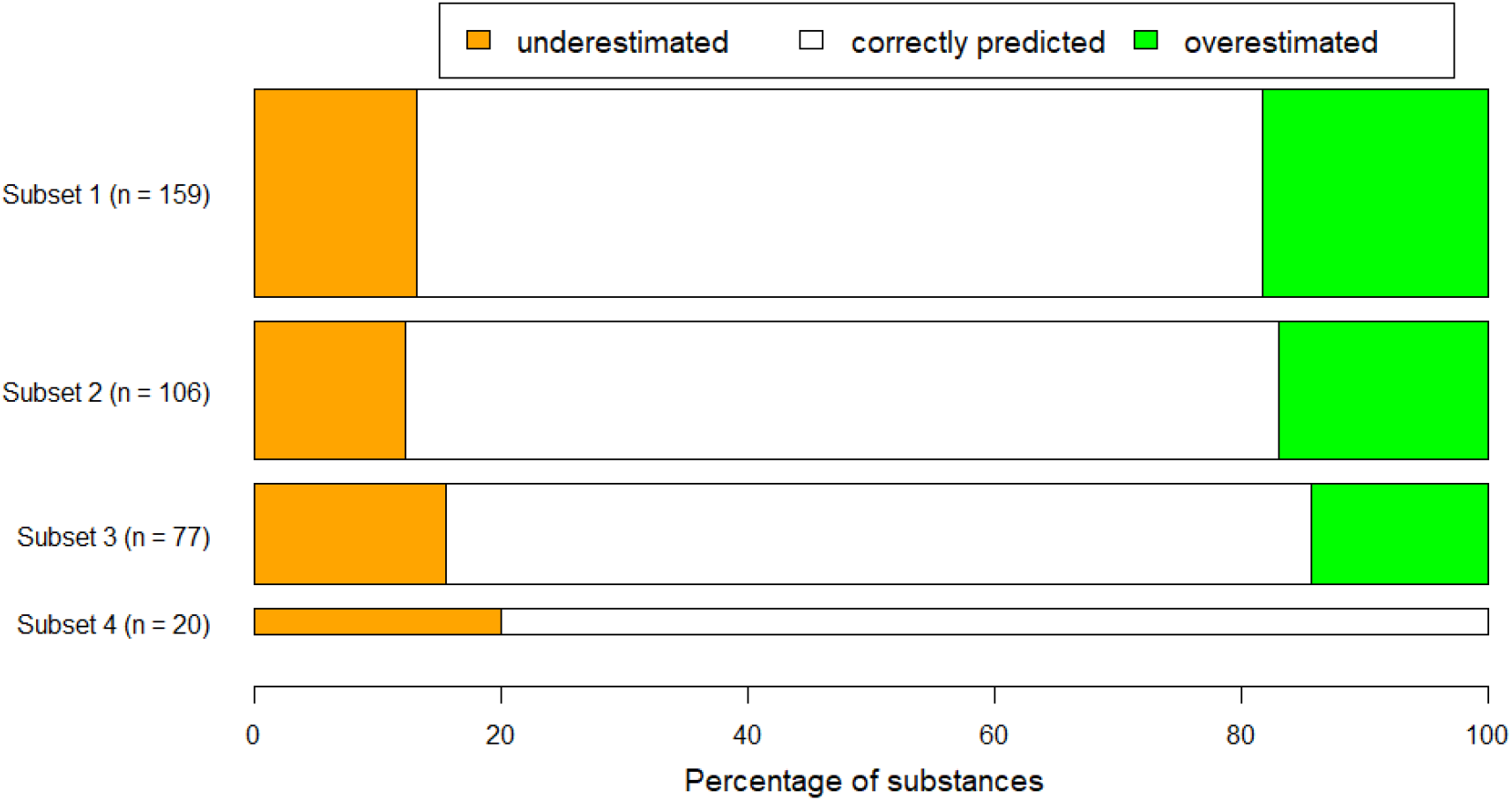
Summary of the BN model’s perfomance: the ability to predict the correct interval of toxicity to juvenile fish for the four subsets of chemical substances (see Table 7). “Correctly predicted” means that the predicted toxicity interval (i.e. the interval with the highest posterior probability) for a given substance is identical to the observed toxicity interval (i.e. the interval with the highest frequency of observations for juvenile fish) for that substance. “Underestimated” means that predicted toxicity interval is lower than the observed interval. The height of each bar corresponds to the number of substances (n) in the subset. For more details, see Table 7.

The fourth subset, which had the strictest criteria for selection of substances (minimum 3 observations of both juvenile and embryo), had the highest correct prediction rate (80%). Although this increase in prediction rate may simply be a random change due to the lower sample size (n = 20), this improvement could also indicate that the higher number of observations leads to more accurate predictions and therefore better model performance. The better performance of this final subset therefore lends support to our decision of designing the BN with multiple continuous input nodes, which allows for the use of more observations.

In practice, the risk assessment for a chemical substance is determined by the most sensitive endpoint - algae, *Daphnia* or fish. For example, for the selected substances in Subset 4 (Table 7d), the four substances with underestimated toxicity to juvenile fish were all more toxic to algae or *Daphnia* than to fish embryos. In such cases, the predicted toxicity to juvenile fish is less relevant, since the risk assessment will be driven by another endpoint. For this reason, in the web interface to the model, information on the most sensitive endpoint is extracted from the input data and provided in the input table (Table 4a) and in the conclusive statements (Figure S.2). These implications are further discussed by Lillicrap et al. (2019).

### 4.4 Further development of the BN model

Although this preliminary version of the BN model shows a high rate of correct predictions (up to 80%; Figure 4), the model performance should be further improved. Moreover, the model showed relatively low value of information for reducing uncertainty of model predictions, as indicated in Table 6, which suggests that the model needs refinement for making better use of provided input data (cf. Table 4a). To increase the model sensitivity to the lines with weakest influence and to improve the accuracy of model predictions, the following issues should be addressed.

For most nodes in Lines 1-3 (Figure 1), which do not depend on fish embryo toxicity data, the prior probability tables and conditional probability tables could be parametrized based on independent datasets. This way, our dataset could be reserved for model validation. A candidate dataset for model parametrisation is the aquatic toxicology database EnviroTox (Connors et al., 2019). However, the EnviroTox database has not yet received the same level of curation as our dataset and may introduce additional noise into our model. For components of the model where data are not available, the CPTs based on expert knowledge by the authors can be strengthened by a more formal approach for external expert elicitation (e.g. (Castelletti and Soncini-Sessa, 2007; Marcot, 2017)).

The four lines of evidence has so far been integrated by equal weighting as a starting point. The CPT for combining the four lines (Table S.21) could instead be trained by data to optimize the weighting of the four lines. However, training the model by advanced methods such as machine-learning algorithms (Marcot and Penman, 2019) will require a higher number of substances with complete set of input data, including fish embryo toxicity data, than what is currently available.

The toxicity intervals are relatively wide, some spanning more than an order of magnitude (Table 3). Toxicity nodes with higher resolution (more toxicity intervals) may be perceived as more informative. However, a model with more than 5 toxicity levels would require different approaches to parametrization of the CPTs, whether these are based on expert knowledge (e.g. Figure 2a) or on frequency distributions (e.g. Figure 2b).

The width of the toxicity intervals is decreasing exponentially: e.g. the state “low” has width 95 mg/L, “medium” has width 4.5 mg/L, and “high” has width 0.49 mg/L. While this non-linear scale reflects the intervals and threshold values typically used in regulatory ecotoxicology, it introduces a bias in the probability calculations (explained in Appendix A). Lower toxicity levels (corresponding to higher concentrations) with wider toxicity intervals will inherently have lower probability than higher toxicity levels. The BN model may therefore show a tendency of overestimating toxicity (Figure 4). From regulatory point of view, such a bias may not be a problem since the biased predictions will be more protective for the environment. However, while risk assessment calculations often have built-in safety factors to obtain more protective assessments, we aim for accuracy of the model predictions for this BN model. A possible solution for reducing the bias is to model the toxicity values on logarithmic scale, so that the toxicity intervals will get more equal widths.

In principle, this hybrid BN could be further developed into a continuous-variable BN. An example of a continuous-variable BN modelling framework, developed for setting nitrogen criteria in streams and rivers, is presented by (Qian and Miltner, 2015). Their model retained the BN’s graphical representation of hypothesized causal connections among variables, while employing statistical modelling approaches for establishing functional relationships among these variables. A continuous BN model can avoid the problems related to discretisation of continuous toxicity value into intervals (Nojavan A et al., 2017), and provide more precise predictions. However, constructing a continuous-variable BN poses other challenges. In our case, converting the BN into a continuous model would require more advanced approaches to model parameterisation, especially for CPTs that are currently based on expert knowledge.

## 5 Conclusions

We have developed a Bayesian network model for predicting the acute toxicity of chemical substances to juvenile fish, based on information on fish embryo toxicity in combination with physical and chemical properties, chemical category and toxicity of the substance to other taxa. This model can support a Weight-of-Evidence approach for replacing the OECD 203 acute fish toxicity assay (based on juvenile fish) with animal alternative approaches, such as the OECD 236 fish embryo toxicity assay. As a measure of model performance, the BN predicted the correct toxicity interval for 69%-80% of substances in our dataset, given different quality criteria. For the subset of 20 substances with the highest quality criteria, the prediction rate was 80% correct. The model predictions were most sensitive to the model components that were quantified by toxicity data (fish embryo and QSAR), and least sensitive to the components that were quantified by expert knowledge (e.g. involving the chemical’s mode of action inferred from testing of other taxa). The model is publicly available through a web interface for testing and feedback. Future development this model should include more lines of evidence, refinement of the discretisation of toxicity intervals, training of the model with a larger toxicity dataset to weight the lines of evidence differently. A more mature version of this model can be a useful WOE tool for predicting fish acute toxicity from fish embryo toxicity data. The model will also be a contribution to the trend of developing more quantitative weight-of-evidence approaches, which are needed both in the context of animal alternatives in ecotoxicology and in environmental assessment more generally.

## Appendices Appendix A

Conditional probability distributions for continuous toxicity value nodes. The continuous toxicity values nodes for QSAR, algae, *Daphnia* and embryo are input nodes for the BN model, but also child nodes of the respective interval-type nodes. Therefore, their conditional probability distributions must be specified by mean and variance for the seven toxicity intervals, as described in Section 2.4.3 Conditional Probability Tables.

Figure A.1 illustrates the probability distributions for each of the states ranging from very high toxicity (a) to very low toxicity (e), defined as Gaussian probability density functions. For each interval, the variance of the probability density function was set so that 90% of the distribution was within the interval while 5% of the distribution was in either of the neighbour intervals. For example, for the interval “medium toxicity” (Figure A.1c), 90% of the area below the curve is contained within the interval 0.5-5 mg/L, 5% of the area is in the range <0.5 mg/L (“high toxicity” or higher), and 5% of the area is in the range >5 mg/L (“low toxicity” or lower). This can be interpreted as follows. Let us assume that the true but unknown toxicity of a substance to algae is in the interval “medium toxicity”. Then there is a 5% probability that a toxicity test of this substance will wrongly result in “low toxicity” to algae, and 5% probability that the test will wrongly result in “high toxicity”.

For all intervals displayed in Figure A.1., the resulting variances (Figure 2c, Table S1.13) correspond to ca. 30% CV of the upper interval boundary (i.e., the lowest toxicity of the interval).

When the BN model is run, the CPT of the continuous toxicity nodes (e.g., “Toxicity to algae (value 1)”are used together with evidence (inserted continuous toxicity values) to update the probabilities of the interval nodes (e.g., “Toxicity to algae (interval)” by backward calculation. Since the conditional probility distributions of the higher toxicity states have more narrow intervals than the lower toxicity states, the higher toxicity states will generally have higher probability density (Figure A.1). As a consequence, if one enters a continuous toxicity value that is close to the lower boundary (i.e. close to the higher toxicity interval), then the posterior distribution of the interval toxicity node is more likely to be in the higher toxicity interval. For example (Figure A.2a), consider an observed toxicity value of 6 mg/L, which belongs to the interval “low toxicity” (5 - 100 mg/L). However, for this interval, the conditional probability density of 6 mg/L is only 0.0038, while for the neighbour interval “medium toxicity”, the conditional probability density is five times as high (0.019). Therefore, this observation will will result in a relatively high posterior probability of the “medium toxicity” interval. In contrast, consider an observed value is 4 mg/L, which belongs to the interval “medium toxicty”, but is close “low toxicity”. The probability density of this value of the correct interval “medium toxicity” is 0.19 (Figure A.2b), which is 56 times as high as as the probability density of the neigbhour interval “low toxicity” (0.0034). Hence, this observation result in virtually zero probability of “low probability”. In summary, the lower toxicity levels (corresponding to higher concentrations) with wider toxicity intervals will inherently have lower probability than the higher toxicity levels.

**Figure A.1.**
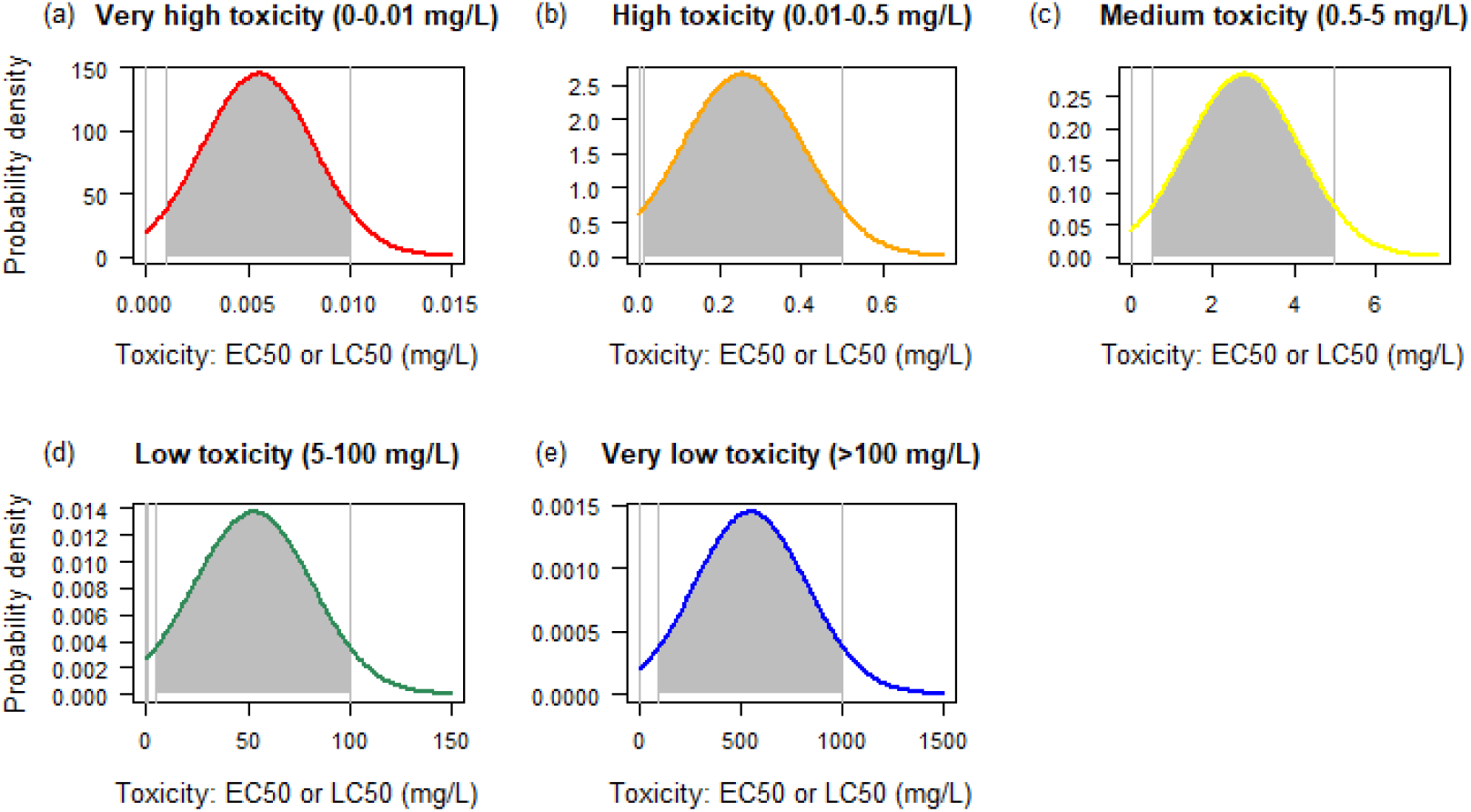
Probability density function used for setting conditional probabilities of the continuous interval nodes, for the five toxicity intervals ranging from very high (a) to very low (e). Coloured curves are the probability density function generated by gaussan distribution with mean and variance as specified in Figure 2c. Vertical lines indicate the toxicity interval boundaries, and the grey areas represent the probability of values occuring in this interval.

**Figure A.2.**
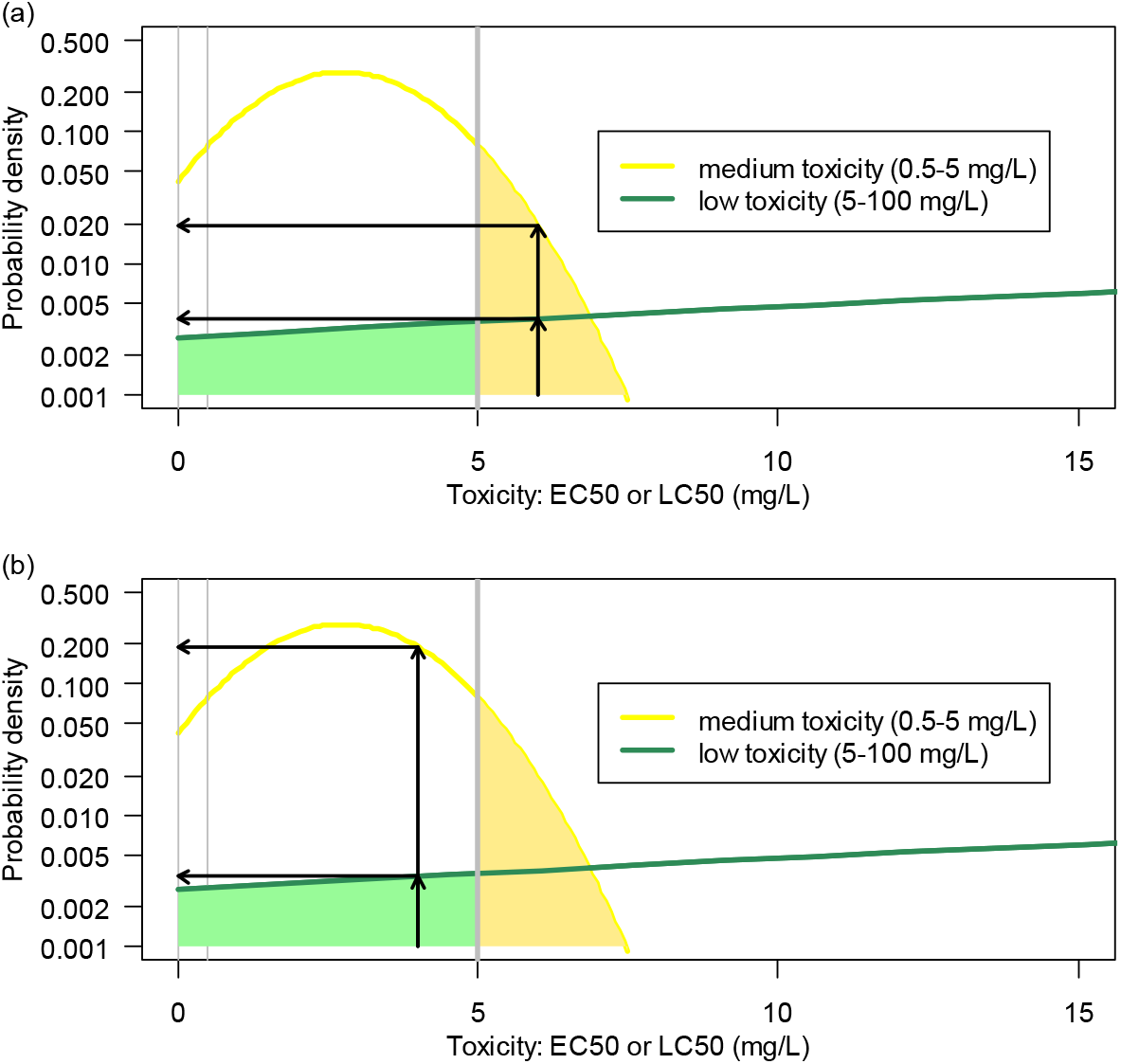
Probability density functions of conditional probabilities for two neighbour toxicity intervals: medium toxiciy (0.5-5 mg/L) and low toxicity (5-100 mg/L). The shaded yellow area represents the probability that a toxicity test of a substance with (true) medium toxicity results in an observation of low toxicity. Conversely, the shaded green area represents the probability that a toxicity test of a substance with (true) low toxicity results in an observation of medium toxicity. Note the logarithmic scale of the y-axis.

## Supplementary material

1. The file “FET_BN_Supplementary_tables_20190828.docx” contains the tables of conditional dependencies of all nodes in the BN model (Tables S.1 to S.21).
2. The file “FET_BN_Supplementary_figures_20190828.docx” contains pictures (Figures S.1 and S.2) showing an example of running the BN model from the web interface (http://demo.hugin.com/demo/FET).

## Author contributions

Conceptualization, A.D.L., S.E.B. and S.J.M; Data curation, K.A.C, J.M.R and S.E.B.; Data analysis, S.J.M.; A.L.M. and R.W., Methodology, S.J.M., A.D.L., W.G.L., A.L.M. and R.W.; Funding acquisition, S.J.M. and A.D.L.; Project administration, S.J.M.; Web interface, A.L.M; Visualization, S.J.M.; Writing – original draft, S.J.M. and A.L.M.; Writing – review & editing; S.J.M, R.W., A.L.M., A.D.L, K.A.C., J.M.R., W.G.L.

