## Supplementary tables for "Development of a hybrid Bayesian network model for predicting acute fish toxicity using multiple lines of evidence"

This supplementary file provides the parameters of all nodes of the Bayesian network model, as prior probabilities (for root nodes), conditional density functions (for continuous child nodes) and conditional probability tables (for interval and categorical child nodes). Table S.1 gives the list of nodes with node names (used in Tables S.2-S.21, which are generated by the BN software) and node labels (used in Figures 1-3 and elsewhere in the main text)

Table S.1. List of probability tables for all nodes in the BN model. Type of probability distribution: prior = prior probability distribution, CDF = conditional density function, CPT = conditional probability table.

| **Line no.** | **Node name** | **Node label** | **Table no.** | **Type of probability distribution** |
| --- | --- | --- | --- | --- |
| 1 | mol_weight | Molecular weight (interval) | S.2 | prior |
| 1 | Molweight_val | Molecular weight (value) | S.3 | CDF |
| 1 | Hydrophobicity | Hydrophobicity (interval) | S.4 | prior |
| 1 | Hydrophobicity_val | Hydrophobicity (value) | S.5 | CDF |
| 1 | cross_membrane | Membrane crossing | S.6 | CPT |
| 1 | Tox_QSAR_int | Toxicity based on QSAR (interval) | S.7 | prior |
| 1 | QSAR_val | Toxicity based on QSAR (value) | S.8 | CDF |
| 1 | QSAR | Toxicity based on QSAR (level) | S.9 | CPT |
| 1 | Tox_fish_chem_prop | Toxicity to fish predicted from chemical properties | S.10 | CPT |
| 2 | Chemical_category1 | Chemcial category | S.11 | prior |
| 2 | Chemical_Category_juvenile1 | Toxicity to fish predicted from chemical category | S.12 | CPT |
| 3 | Tox_algae_int | Toxicity to algae (interval) | S.13 | prior |
| 3 | Tox_algae_val_1^1)^ | Toxicity to algae (value 1) | S.8 | CDF |
| 3 | Tox_algae_lev | Toxicity to algae (level) | S.9 | CPT |
| 3 | Tox_Daphnia_int | Toxicity to Daphnia (interval) | S.14 | prior |
| 3 | Tox_Daphnia_val_1^1)^ | Toxicity to Daphnia (value 1) | S.8 | CDF |
| 3 | Tox_Daphnia_lev | Toxicity to Daphnia (level) | S.9 | CPT |
| 3 | Daphnia_algae_ratio | Ratio toxicity Daphnia / algae | S.15 | prior |
| 3 | Species_specific_MoA | Species-specific mode of action | S.16 | CPT |
| 3 | Tox_fish_oth_species | Toxicity to fish predicted from other taxa | S.17 | CPT |
| 4 | Tox_emb_int | Toxicity to embryo (interval) | S.18 | prior |
| 4 | Tox_embryo_val_1^1)^ | Toxicity to embryo (value 1) | S.8 | CDF |
| 4 | Tox_emb_lev | Toxicity to embryo (level) | S.9 | CPT |
| all | ^2)^ | Toxicity to fish predicted from all evidence (1) | S.19 | CPT |
| all | ^2)^ | Toxicity to fish predicted from all evidence (2) | S.20 | CPT |
| all | Tox_fish_pred | Toxicity to fish predicted from all evidence | S.21 | CPT |

^1)^ The CDF for a continuous value node is identical for all of its sibling nodes

^2)^ The nodes "Toxicity to fish predicted from all evidence (1)" and "Toxicity to fish predicted from all evidence (2)" were temporary nodes, which were used to facilitate the generation of the CPT for "Toxicity to fish predicted from all evidence". An algorithm was developed for combining two parent nodes with 5 toxicity states (Table S.19 and Table S.20). One node also contains the sixth state "unknown" (Table S.19). The two intermediate nodes (Toxicity to fish predicted from all evidence (1) and (2)) were subsequently combined using the same algorithm. The two intermediate nodes were absorbed to generate a CPT combining all four parent nodes. For cases with state = unknown for the parent node"Chemical category", some probability distributions were adjusted to let the other three parent nodes have equal weight (e.g. from 0:0.5:0.25:0.25 to 0:0.33:0.33:0.33).

Line 1: Toxicity based on physical and chemical properties

Table S.2. Molecular weight (interval)

| 0 - 600 | 0.991416 |
| --- | --- |
| 600 - inf | 0.008584 |

Table S.3. Molecular weight (value)

| mol_weight | <600 | >600 |
| --- | --- | --- |
| Mean | 179.337 | 200380 |
| Variance | 321.616 | 3.614E9 |

Table S.4. Hydrophobicity (interval)

| -inf - 5.5 | 0.968182 |
| --- | --- |
| 5.5 - inf | 0.031818 |

Table S.5. Hydrophobicity (value)

| Hydrophobicity | <5.5 | >5.5 |
| --- | --- | --- |
| Mean | 1.733 | 6.50343 |
| Variance | 0.0075081 | 0.105736 |

Table S.6. Membrane crossing

| Hydrophobicity | <5.5 | | >5.5 | |
| --- | --- | --- | --- | --- |
| mol_weight | <600 | >600 | <600 | >600 |
| low | 0 | 0.25 | 0.25 | 0.75 |
| medium | 0.25 | 0.5 | 0.5 | 0.25 |
| high | 0.75 | 0.25 | 0.25 | 0 |

Table S.7. Toxicity based on QSAR (interval)

| 0 - 0.001 | 6.57895E-6 |
| --- | --- |
| 0.001 - 0.01 | 0.006572 |
| 0.01 - 0.5 | 0.131579 |
| 0.5 - 5 | 0.190789 |
| 5 - 100 | 0.414474 |
| 100 - 1000 | 0.256322 |
| 1000 - inf | 0.000257 |

Table S.8. Toxicity based on QSAR (value)

| Tox_QSAR_int | 0 - 0.001 | 0.001 - 0.01 | 0.01 - 0.5 | 0.5 - 5 | 5 - 100 | 100 - 1000 | 1000 - inf |
| --- | --- | --- | --- | --- | --- | --- | --- |
| Mean | 0.0005 | 0.0055 | 0.255 | 2.75 | 52.5 | 550 | 1550 |
| Variance | 7.5625E-8 | 7.5625E-6 | 0.0225 | 2.25 | 756.25 | 75625 | 600625 |

Table S.9. Toxicity based on QSAR (level)

| Tox_QSAR_int | 0 - 0.001 | 0.001 - 0.01 | 0.01 - 0.5 | 0.5 - 5 | 5 - 100 | 100 - 1000 | 1000 - inf |
| --- | --- | --- | --- | --- | --- | --- | --- |
| very low | 0 | 0 | 0 | 0 | 0 | 1 | 1 |
| low | 0 | 0 | 0 | 0 | 1 | 0 | 0 |
| medium | 0 | 0 | 0 | 1 | 0 | 0 | 0 |
| high | 0 | 0 | 1 | 0 | 0 | 0 | 0 |
| very high | 1 | 1 | 0 | 0 | 0 | 0 | 0 |

Table S.10. Toxicity to fish predicted from chemical properties

| cross_membrane | low | | | | | medium | | | | |
| --- | --- | --- | --- | --- | --- | --- | --- | --- | --- | --- |
| QSAR | very low | low | medium | high | very high | very low | low | medium | high | very high |
| very low | 0.8 | 0.1 | 0 | 0 | 0 | 0.9 | 0.05 | 0 | 0 | 0 |
| low | 0.2 | 0.8 | 0.1 | 0 | 0 | 0.1 | 0.9 | 0.05 | 0 | 0 |
| medium | 0 | 0.1 | 0.8 | 0.1 | 0 | 0 | 0.05 | 0.9 | 0.05 | 0 |
| high | 0 | 0 | 0.1 | 0.8 | 0.2 | 0 | 0 | 0.05 | 0.9 | 0.1 |
| very high | 0 | 0 | 0 | 0.1 | 0.8 | 0 | 0 | 0 | 0.05 | 0.9 |

| cross_membrane | high | | | | |
| --- | --- | --- | --- | --- | --- |
| QSAR | very low | low | medium | high | very high |
| very low | 1 | 0 | 0 | 0 | 0 |
| low | 0 | 1 | 0 | 0 | 0 |
| medium | 0 | 0 | 1 | 0 | 0 |
| high | 0 | 0 | 0 | 1 | 0 |
| very high | 0 | 0 | 0 | 0 | 1 |

Line 2: Toxicity based on chemical category

Table S.11. Chemcial category. Note: this table also displays the count of substances in our dataset per chemical category (with any type of data) and the total count of observations of toxicity to juvenile fish within the category.

| Chemical category | Prior probability | Count of substances | Count of observations |
| --- | --- | --- | --- |
| Acrylates | 0.0233 | 1 | 1 |
| Aldehydes (mono) | 0.0233 | 1 | 8 |
| Aliphatic amine | 0.0233 | 56 | 60 |
| Amide | 0.0233 | 5 | 2 |
| Aniline | 0.0233 | 19 | 50 |
| Anionic surfactant | 0.0233 | 10 | 42 |
| Carbamate ester | 0.0233 | 1 | 82 |
| Carbamate esters, phenyl | 0.0233 | 1 | 14 |
| Cationic surfactant | 0.0233 | 5 | 4 |
| Dintrophenol | 0.0233 | 1 | 19 |
| Epoxides, mono | 0.0233 | 1 | 58 |
| Ester | 0.0233 | 4 | 37 |
| Esters (dithiophosphates) | 0.0233 | 2 | 76 |
| Esters (monothiphosphates) | 0.0233 | 1 | 39 |
| Halo alcohols | 0.0233 | 1 | 4 |
| Hydrazine | 0.0233 | 4 | 1 |
| Imidazole | 0.0233 | 2 | 4 |
| Imide | 0.0233 | 1 | 1 |
| Inorganic | 0.0233 | 2 | 58 |
| Metal | 0.0233 | 7 | 202 |
| Monoaldehyde | 0.0233 | 2 | 1 |
| Neutral organic | 0.0233 | 39 | 145 |
| Neutral organic (acid) | 0.0233 | 12 | 10 |
| Nonionic surfactant | 0.0233 | 4 | 2 |
| Organometal | 0.0233 | 1 | 7 |
| Pesticide | 0.0233 | 2 | 89 |
| Phenol | 0.0233 | 19 | 204 |
| Phenols-acid | 0.0233 | 1 | 3 |
| Polyme0072 | 0.0233 | 3 | 2 |
| Polynitrophenol | 0.0233 | 1 | 8 |
| Polyphenol | 0.0233 | 2 | 10 |
| Pyrethroid | 0.0233 | 1 | 42 |
| Quinone | 0.0233 | 2 | 13 |
| Quinone/Hydroquinone | 0.0233 | 2 | 7 |
| Substituted urea | 0.0233 | 3 | 20 |
| Thiocarbamate, di | 0.0233 | 1 | 9 |
| Triazine | 0.0233 | 1 | 21 |
| Triazoles | 0.0233 | 1 | 6 |
| Vinyl allyl aldehyde | 0.0233 | 1 | 15 |
| Vinyl/Allyl ether | 0.0233 | 1 | 67 |
| Vinyl/Allyl halide | 0.0233 | 1 | 15 |
| Zwitterionic surfactant | 0.0233 | 1 | 1 |

Table S.12. Toxicity to fish predicted from chemical category

| Chemical category | Acrylates | Aldehydes (mono) | Aliphatic amine | Amide | Aniline | Anionic surfactant | Carbamate ester |
| --- | --- | --- | --- | --- | --- | --- | --- |
| very low | 0 | 0 | 0.5 | 1 | 0.08 | 0 | 0 |
| low | 0 | 0.75 | 0.366666667 | 0 | 0.8 | 0.333333333 | 0.487804878 |
| medium | 1 | 0.25 | 0.116666667 | 0 | 0.08 | 0.571428571 | 0.512195122 |
| high | 0 | 0 | 0.016666667 | 0 | 0.04 | 0.095238095 | 0 |
| very high | 0 | 0 | 0 | 0 | 0 | 0 | 0 |
| unknown | 0 | 0 | 0 | 0 | 0 | 0 | 0 |

| Chemical category | Carbamate esters, phenyl | Cationic surfactant | Dintrophenol | Epoxides, mono | Ester | Esters (dithiophosphates) | Esters (monothiphosphates) |
| --- | --- | --- | --- | --- | --- | --- | --- |
| very low | 0.071428571 | 0.25 | 0 | 0 | 0.189189189 | 0 | 0 |
| low | 0.714285714 | 0.25 | 0.736842105 | 0 | 0.216216216 | 0.184210526 | 0.025641026 |
| medium | 0.214285714 | 0.5 | 0.210526316 | 0 | 0.567567568 | 0.263157895 | 0.564102564 |
| high | 0 | 0 | 0.052631579 | 0 | 0.027027027 | 0.5 | 0.41025641 |
| very high | 0 | 0 | 0 | 1 | 0 | 0.052631579 | 0 |
| unknown | 0 | 0 | 0 | 0 | 0 | 0 | 0 |

| Chemical category | Halo alcohols | Hydrazine | Imidazole | Imide | Inorganic | Metal | Monoaldehyde |
| --- | --- | --- | --- | --- | --- | --- | --- |
| very low | 1 | 0 | 0.25 | 0 | 0.431034483 | 0.148514851 | 0 |
| low | 0 | 1 | 0 | 0 | 0.137931034 | 0.400990099 | 1 |
| medium | 0 | 0 | 0.75 | 1 | 0.25862069 | 0.242574257 | 0 |
| high | 0 | 0 | 0 | 0 | 0.172413793 | 0.118811881 | 0 |
| very high | 0 | 0 | 0 | 0 | 0 | 0.089108911 | 0 |
| unknown | 0 | 0 | 0 | 0 | 0 | 0 | 0 |

| Chemical category | Neutral organic | Neutral organic (acid) | Nonionic surfactant | Organometal | Pesticide | Phenol | Phenols-acid |
| --- | --- | --- | --- | --- | --- | --- | --- |
| very low | 0.455172414 | 0.9 | 0 | 0 | 0 | 0 | 0.666666667 |
| low | 0.344827586 | 0.1 | 0 | 0 | 0.157303371 | 0.362745098 | 0.333333333 |
| medium | 0.165517241 | 0 | 0.5 | 0 | 0.280898876 | 0.117647059 | 0 |
| high | 0.020689655 | 0 | 0.5 | 1 | 0.370786517 | 0.519607843 | 0 |
| very high | 0.013793103 | 0 | 0 | 0 | 0.191011236 | 0 | 0 |
| unknown | 0 | 0 | 0 | 0 | 0 | 0 | 0 |

| Chemical category | Polymer | Polynitrophenol | Polyphenol | Pyrethroid | Quinone | Quinone/Hydroquinone | Substituted urea |
| --- | --- | --- | --- | --- | --- | --- | --- |
| very low | 0 | 0 | 0 | 0 | 0 | 0 | 0 |
| low | 0 | 0 | 0.4 | 0 | 0 | 0 | 0.45 |
| medium | 0.5 | 0.625 | 0.5 | 0 | 0 | 0.142857143 | 0.05 |
| high | 0.5 | 0.375 | 0.1 | 0.119047619 | 1 | 0.857142857 | 0.5 |
| very high | 0 | 0 | 0 | 0.880952381 | 0 | 0 | 0 |
| unknown | 0 | 0 | 0 | 0 | 0 | 0 | 0 |

| Chemical category | Thiocarbamate, di | Triazine | Triazoles | Vinyl allyl aldehyde | Vinyl/Allyl ether | Vinyl/Allyl halide | Zwitterionic surfactant |
| --- | --- | --- | --- | --- | --- | --- | --- |
| very low | 0 | 0 | 0 | 0 | 0 | 0 | 0 |
| low | 0 | 0.857142857 | 0 | 0 | 0 | 0.733333333 | 1 |
| medium | 0 | 0.142857143 | 1 | 0 | 0 | 0.266666667 | 0 |
| high | 1 | 0 | 0 | 1 | 0.671641791 | 0 | 0 |
| very high | 0 | 0 | 0 | 0 | 0.328358209 | 0 | 0 |
| unknown | 0 | 0 | 0 | 0 | 0 | 0 | 0 |

Line 3: Toxicity based on other taxa

Table S.13: Toxicity to algae (interval)

| 0 - 0.001 | 2.27273E-5 |
| --- | --- |
| 0.001 - 0.01 | 0.022704 |
| 0.01 - 0.5 | 0.318182 |
| 0.5 - 5 | 0.253788 |
| 5 - 100 | 0.318182 |
| 100 - 1000 | 0.087034 |
| 1000 - inf | 8.71212E-5 |

Toxicity to algae (value 1); ...; Toxicity to algae (value 10): see Table S.8: Toxicity based on QSAR

Toxicity to algae (level): See Table S.9 Toxicity based on QSAR (level)

Table S.14: Toxicity to Daphnia (interval)

| 0 - 0.001 | 0.000133 |
| --- | --- |
| 0.001 - 0.01 | 0.133028 |
| 0.01 - 0.5 | 0.353952 |
| 0.5 - 5 | 0.189863 |
| 5 - 100 | 0.219931 |
| 100 - 1000 | 0.10299 |
| 1000 - inf | 0.000103 |

Toxicity to Daphnia (value 1); ...; Toxicity to Daphnia (value 10): see Table S.8: Toxicity based on QSAR

Toxicity to Daphnia (level): See Table S.9 Toxicity based on QSAR (level)

Table S.15: Ratio toxicity Daphnia / algae

| 0 - 0.5 | 0.467066 |
| --- | --- |
| 0.5 - 2 | 0.52994 |
| 2 - inf | 0.002994 |

Table S.16: Species-specific mode of action

| Daphnia_algae_ratio | 0 - 0.5 | 0.5 - 2 | 2 - inf |
| --- | --- | --- | --- |
| different | 1 | 0 | 1 |
| similar | 0 | 1 | 0 |

Table S.17: Toxicity to fish predicted from other taxa

| Species_specific_MoA | different | | | | | | | | | |
| --- | --- | --- | --- | --- | --- | --- | --- | --- | --- | --- |
| Tox_Daphnia_lev | very low | | | | | low | | | | |
| Tox_algae_lev | very low | low | medium | high | very | very low | low | medium | high | very |
| very low | 0.3 | 0.3 | 0.2 | 0.25 | 0.5 | 0.25 | 0.3 | 0.15 | 0.25 | 0.25 |
| low | 0.3 | 0.3 | 0.3 | 0.5 | 0.25 | 0.25 | 0.3 | 0.3 | 0.5 | 0.5 |
| medium | 0.2 | 0.2 | 0.2 | 0.25 | 0.25 | 0.25 | 0.2 | 0.3 | 0.25 | 0.25 |
| high | 0.1 | 0.1 | 0.2 | 0 | 0 | 0.15 | 0.1 | 0.15 | 0 | 0 |
| very high | 0.1 | 0.1 | 0.1 | 0 | 0 | 0.1 | 0.1 | 0.1 | 0 | 0 |

| Species_specific_MoA | different | | | | | | | | | |
| --- | --- | --- | --- | --- | --- | --- | --- | --- | --- | --- |
| Tox_Daphnia_lev | medium | | | | | high | | | | |
| Tox_algae_lev | very low | low | medium | high | very | very low | low | medium | high | very |
| very low | 0.1 | 0.1 | 0.1 | 0.1 | 0.1 | 0.1 | 0.1 | 0.1 | 0.1 | 0.1 |
| low | 0.25 | 0.25 | 0.25 | 0.2 | 0.2 | 0.2 | 0.2 | 0.2 | 0.1 | 0.1 |
| medium | 0.3 | 0.3 | 0.3 | 0.4 | 0.4 | 0.25 | 0.25 | 0.25 | 0.25 | 0.2 |
| high | 0.25 | 0.25 | 0.25 | 0.2 | 0.2 | 0.3 | 0.3 | 0.3 | 0.3 | 0.3 |
| very high | 0.1 | 0.1 | 0.1 | 0.1 | 0.1 | 0.15 | 0.15 | 0.15 | 0.25 | 0.3 |

| Species_specific_MoA | different | | | | | similar | | | | |
| --- | --- | --- | --- | --- | --- | --- | --- | --- | --- | --- |
| Tox_Daphnia_lev | very high | | | | | very low | | | | |
| Tox_algae_lev | very low | low | medium | high | very | very low | low | medium | high | very |
| very low | 0.1 | 0.1 | 0.1 | 0.1 | 0.1 | 1 | 0.5 | 0.2 | 0.2 | 0.2 |
| low | 0.15 | 0.15 | 0.15 | 0.1 | 0.1 | 0 | 0.5 | 0.2 | 0.2 | 0.2 |
| medium | 0.2 | 0.2 | 0.2 | 0.2 | 0.2 | 0 | 0 | 0.2 | 0.2 | 0.2 |
| high | 0.3 | 0.3 | 0.3 | 0.3 | 0.3 | 0 | 0 | 0.2 | 0.2 | 0.2 |
| very high | 0.25 | 0.25 | 0.25 | 0.3 | 0.3 | 0 | 0 | 0.2 | 0.2 | 0.2 |

| Species_specific_MoA | similar | | | | | | | | | |
| --- | --- | --- | --- | --- | --- | --- | --- | --- | --- | --- |
| Tox_Daphnia_lev | low | | | | | medium | | | | |
| Tox_algae_lev | very low | low | medium | high | very | very low | low | medium | high | very |
| very low | 0.5 | 0.25 | 0 | 0.2 | 0.2 | 0.2 | 0.25 | 0.1 | 0.1 | 0.2 |
| low | 0.5 | 0.5 | 0.5 | 0.2 | 0.2 | 0.2 | 0.25 | 0.25 | 0.25 | 0.2 |
| medium | 0 | 0.25 | 0.5 | 0.2 | 0.2 | 0.2 | 0.25 | 0.3 | 0.3 | 0.2 |
| high | 0 | 0 | 0 | 0.2 | 0.2 | 0.2 | 0.25 | 0.25 | 0.25 | 0.2 |
| very high | 0 | 0 | 0 | 0.2 | 0.2 | 0.2 | 0 | 0.1 | 0.1 | 0.2 |

| Species_specific_MoA | similar | | | | | | | | | |
| --- | --- | --- | --- | --- | --- | --- | --- | --- | --- | --- |
| Tox_Daphnia_lev | high | | | | | very high | | | | |
| Tox_algae_lev | very low | low | medium | high | very high | very low | low | medium | high | very high |
| very low | 0.2 | 0.2 | 0 | 0 | 0 | 0.2 | 0.2 | 0.2 | 0 | 0 |
| low | 0.2 | 0.2 | 0.1 | 0 | 0 | 0.2 | 0.2 | 0.2 | 0 | 0 |
| medium | 0.2 | 0.2 | 0.4 | 0.25 | 0 | 0.2 | 0.2 | 0.2 | 0 | 0 |
| high | 0.2 | 0.2 | 0.4 | 0.5 | 0.5 | 0.2 | 0.2 | 0.2 | 0.5 | 0 |
| very high | 0.2 | 0.2 | 0.1 | 0.25 | 0.5 | 0.2 | 0.2 | 0.2 | 0.5 | 1 |

Line 4: Toxicity based on embryo

Table S.18: Toxicity to embryo (interval)

| 0 - 0.001 | 7.39373E-6 |
| --- | --- |
| 0.001 - 0.01 | 0.007386 |
| 0.01 - 0.5 | 0.151571 |
| 0.5 - 5 | 0.260628 |
| 5 - 100 | 0.325323 |
| 100 - 1000 | 0.254828 |
| 1000 - inf | 0.000255 |

Toxicity to embryo (value 1); ...; Toxicity to embryo (value 10): see Table S.8: Toxicity based on QSAR

Toxicity to embryo (level): See Table S.9 Toxicity based on QSAR (level)

Combination of the the 4 lines of evidence

Table S.19: Toxicity to fish predicted from all evidence (1). Truncated node names: "Chemical_Cate" = ChemicalCategory1; "Tox_fish_chem" = Tox_fish_chem_prop.

| Chemical_Cate | very low | | | | | low | | | | |
| --- | --- | --- | --- | --- | --- | --- | --- | --- | --- | --- |
| Tox_fish_chem | very low | low | medium | high | very | very low | low | medium | high | very |
| very low | 1 | 0.5 | 0.33333 | 0.25 | 0.2 | 0.5 | 0 | 0 | 0 | 0 |
| low | 0 | 0.5 | 0.33333 | 0.25 | 0.2 | 0.5 | 1 | 0.5 | 0.33333 | 0.25 |
| medium | 0 | 0 | 0.33333 | 0.25 | 0.2 | 0 | 0 | 0.5 | 0.33333 | 0.25 |
| high | 0 | 0 | 0 | 0.25 | 0.2 | 0 | 0 | 0 | 0.33333 | 0.25 |
| very high | 0 | 0 | 0 | 0 | 0.2 | 0 | 0 | 0 | 0 | 0.25 |

| Chemical_Cate | medium | | | | | high | | | | |
| --- | --- | --- | --- | --- | --- | --- | --- | --- | --- | --- |
| Tox_fish_chem | very low | low | medium | high | very | very low | low | medium | high | very |
| very low | 0.33333 | 0 | 0 | 0 | 0 | 0.25 | 0 | 0 | 0 | 0 |
| low | 0.33333 | 0.5 | 0 | 0 | 0 | 0.25 | 0.33333 | 0 | 0 | 0 |
| medium | 0.33333 | 0.5 | 1 | 0.5 | 0.33333 | 0.25 | 0.33333 | 0.5 | 0 | 0 |
| high | 0 | 0 | 0 | 0.5 | 0.33333 | 0.25 | 0.33333 | 0.5 | 1 | 0.5 |
| very high | 0 | 0 | 0 | 0 | 0.33333 | 0 | 0 | 0 | 0 | 0.5 |

| Chemical_Cate | very high | | | | | unknown | | | | |
| --- | --- | --- | --- | --- | --- | --- | --- | --- | --- | --- |
| Tox_fish_chem | very low | low | medium | high | very | very low | low | medium | high | very |
| very low | 0.2 | 0 | 0 | 0 | 0 | 1 | 0 | 0 | 0 | 0 |
| low | 0.2 | 0.25 | 0 | 0 | 0 | 0 | 1 | 0 | 0 | 0 |
| medium | 0.2 | 0.25 | 0.33333 | 0 | 0 | 0 | 0 | 1 | 0 | 0 |
| high | 0.2 | 0.25 | 0.33333 | 0.5 | 0 | 0 | 0 | 0 | 1 | 0 |
| very high | 0.2 | 0.25 | 0.33333 | 0.5 | 1 | 0 | 0 | 0 | 0 | 1 |

Table S.20. Toxicity to fish predicted from all evidence (2). Truncated node names: "Tox_fish_oth_s" = Tox_fish_oth_species.

| Tox_fish_oth_s | very low | | | | | low | | | | |
| --- | --- | --- | --- | --- | --- | --- | --- | --- | --- | --- |
| Tox_emb_lev | very low | low | medium | high | very | very low | low | medium | high | very |
| very low | 1 | 0.5 | 0.33333 | 0.25 | 0.2 | 0.5 | 0 | 0 | 0 | 0 |
| low | 0 | 0.5 | 0.33333 | 0.25 | 0.2 | 0.5 | 1 | 0.5 | 0.33333 | 0.25 |
| medium | 0 | 0 | 0.33333 | 0.25 | 0.2 | 0 | 0 | 0.5 | 0.33333 | 0.25 |
| high | 0 | 0 | 0 | 0.25 | 0.2 | 0 | 0 | 0 | 0.33333 | 0.25 |
| very high | 0 | 0 | 0 | 0 | 0.2 | 0 | 0 | 0 | 0 | 0.25 |

| Tox_fish_oth_s | medium | | | | | high | | | | |
| --- | --- | --- | --- | --- | --- | --- | --- | --- | --- | --- |
| Tox_emb_lev | very low | low | medium | high | very | very low | low | medium | high | very |
| very low | 0.33333 | 0 | 0 | 0 | 0 | 0.25 | 0 | 0 | 0 | 0 |
| low | 0.33333 | 0.5 | 0 | 0 | 0 | 0.25 | 0.33333 | 0 | 0 | 0 |
| medium | 0.33333 | 0.5 | 1 | 0.5 | 0.33333 | 0.25 | 0.33333 | 0.5 | 0 | 0 |
| high | 0 | 0 | 0 | 0.5 | 0.33333 | 0.25 | 0.33333 | 0.5 | 1 | 0.5 |
| very high | 0 | 0 | 0 | 0 | 0.33333 | 0 | 0 | 0 | 0 | 0.5 |

| Tox_fish_oth_s | very high | | | | |
| --- | --- | --- | --- | --- | --- |
| Tox_emb_lev | very low | low | medium | high | very |
| very low | 0.2 | 0 | 0 | 0 | 0 |
| low | 0.2 | 0.25 | 0 | 0 | 0 |
| medium | 0.2 | 0.25 | 0.33333 | 0 | 0 |
| high | 0.2 | 0.25 | 0.33333 | 0.5 | 0 |
| very high | 0.2 | 0.25 | 0.33333 | 0.5 | 1 |

Table S.21. Toxicity to fish predicted from all evidence. Truncated node names: "Chemical_Cat" = ChemicalCategory1; "Tox_fish_che" = Tox_fish_chem_prop; "Tox_fish_oth " = Tox_fish_oth_species.

| Chemical_Cate | very low | | | | | | | | | |
| --- | --- | --- | --- | --- | --- | --- | --- | --- | --- | --- |
| Tox_fish_chem | very low | | | | | | | | | |
| Tox_fish_oth_s | very low | | | | | low | | | | |
| Tox_emb_lev | very low | low | medium | high | very high | very low | low | medium | high | very high |
| very low | 1 | 0.75 | 0.611111 | 0.520834 | 0.456667 | 0.75 | 0.5 | 0.416667 | 0.361111 | 0.320833 |
| low | 0 | 0.25 | 0.277778 | 0.270833 | 0.256667 | 0.25 | 0.5 | 0.416667 | 0.361111 | 0.320833 |
| medium | 0 | 0 | 0.111111 | 0.145833 | 0.156667 | 0 | 0 | 0.166667 | 0.194444 | 0.195833 |
| high | 0 | 0 | 0 | 0.0625 | 0.09 | 0 | 0 | 0 | 0.083334 | 0.1125 |
| very high | 0 | 0 | 0 | 0 | 0.04 | 0 | 0 | 0 | 0 | 0.05 |

| Chemical_Cate | very low | | | | | | | | | |
| --- | --- | --- | --- | --- | --- | --- | --- | --- | --- | --- |
| Tox_fish_chem | very low | | | | | | | | | |
| Tox_fish_oth_s | medium | | | | | high | | | | |
| Tox_emb_lev | very low | low | medium | high | very high | very low | low | medium | high | very high |
| very low | 0.611111 | 0.416667 | 0.333333 | 0.291667 | 0.261111 | 0.520834 | 0.361111 | 0.291667 | 0.25 | 0.225 |
| low | 0.277778 | 0.416667 | 0.333333 | 0.291667 | 0.261111 | 0.270833 | 0.361111 | 0.291667 | 0.25 | 0.225 |
| medium | 0.111111 | 0.166667 | 0.333333 | 0.291667 | 0.261111 | 0.145833 | 0.194444 | 0.291667 | 0.25 | 0.225 |
| high | 0 | 0 | 0 | 0.125 | 0.15 | 0.0625 | 0.083334 | 0.125 | 0.25 | 0.225 |
| very high | 0 | 0 | 0 | 0 | 0.066667 | 0 | 0 | 0 | 0 | 0.1 |

| Chemical_Cate | very low | | | | | | | | | |
| --- | --- | --- | --- | --- | --- | --- | --- | --- | --- | --- |
| Tox_fish_chem | very low | | | | | low | | | | |
| Tox_fish_oth_s | very high | | | | | very low | | | | |
| Tox_emb_lev | very low | low | medium | high | very high | very low | low | medium | high | very high |
| very low | 0.456667 | 0.320833 | 0.261111 | 0.225 | 0.2 | 0.75 | 0.5 | 0.388889 | 0.322917 | 0.278333 |
| low | 0.256667 | 0.320833 | 0.261111 | 0.225 | 0.2 | 0.25 | 0.5 | 0.472222 | 0.427083 | 0.386667 |
| medium | 0.156667 | 0.195833 | 0.261111 | 0.225 | 0.2 | 0 | 0 | 0.138889 | 0.177083 | 0.186667 |
| high | 0.09 | 0.1125 | 0.15 | 0.225 | 0.2 | 0 | 0 | 0 | 0.072917 | 0.103333 |
| very high | 0.04 | 0.05 | 0.066667 | 0.1 | 0.2 | 0 | 0 | 0 | 0 | 0.045 |

| Chemical_Cate | very low | | | | | | | | | |
| --- | --- | --- | --- | --- | --- | --- | --- | --- | --- | --- |
| Tox_fish_chem | low | | | | | | | | | |
| Tox_fish_oth_s | low | | | | | medium | | | | |
| Tox_emb_lev | very low | low | medium | high | very high | very low | low | medium | high | very high |
| very low | 0.5 | 0.25 | 0.208333 | 0.180556 | 0.160417 | 0.388889 | 0.208333 | 0.166667 | 0.145833 | 0.130556 |
| low | 0.5 | 0.75 | 0.583334 | 0.486111 | 0.420833 | 0.472222 | 0.583334 | 0.416667 | 0.354167 | 0.311111 |
| medium | 0 | 0 | 0.208333 | 0.236111 | 0.233333 | 0.138889 | 0.208333 | 0.416667 | 0.354167 | 0.311111 |
| high | 0 | 0 | 0 | 0.097222 | 0.129167 | 0 | 0 | 0 | 0.145833 | 0.172222 |
| very high | 0 | 0 | 0 | 0 | 0.05625 | 0 | 0 | 0 | 0 | 0.075 |

| Chemical_Cate | very low | | | | | | | | | |
| --- | --- | --- | --- | --- | --- | --- | --- | --- | --- | --- |
| Tox_fish_chem | low | | | | | | | | | |
| Tox_fish_oth_s | high | | | | | very high | | | | |
| Tox_emb_lev | very low | low | medium | high | very high | very low | low | medium | high | very high |
| very low | 0.322917 | 0.180556 | 0.145833 | 0.125 | 0.1125 | 0.278333 | 0.160417 | 0.130556 | 0.1125 | 0.1 |
| low | 0.427083 | 0.486111 | 0.354167 | 0.291667 | 0.258333 | 0.386667 | 0.420833 | 0.311111 | 0.258333 | 0.225 |
| medium | 0.177083 | 0.236111 | 0.354167 | 0.291667 | 0.258333 | 0.186667 | 0.233333 | 0.311111 | 0.258333 | 0.225 |
| high | 0.072917 | 0.097222 | 0.145833 | 0.291667 | 0.258333 | 0.103333 | 0.129167 | 0.172222 | 0.258333 | 0.225 |
| very high | 0 | 0 | 0 | 0 | 0.1125 | 0.045 | 0.05625 | 0.075 | 0.1125 | 0.225 |

| Chemical_Cate | very low | | | | | | | | | |
| --- | --- | --- | --- | --- | --- | --- | --- | --- | --- | --- |
| Tox_fish_chem | medium | | | | | | | | | |
| Tox_fish_oth_s | very low | | | | | low | | | | |
| Tox_emb_lev | very low | low | medium | high | very high | very low | low | medium | high | very high |
| very low | 0.611111 | 0.388889 | 0.296296 | 0.243056 | 0.207778 | 0.388889 | 0.166667 | 0.138889 | 0.12037 | 0.106944 |
| low | 0.277778 | 0.472222 | 0.407407 | 0.354167 | 0.313333 | 0.472222 | 0.666666 | 0.472222 | 0.37963 | 0.322222 |
| medium | 0.111111 | 0.138889 | 0.296296 | 0.3125 | 0.302222 | 0.138889 | 0.166667 | 0.388889 | 0.37963 | 0.35 |
| high | 0 | 0 | 0 | 0.090278 | 0.124444 | 0 | 0 | 0 | 0.12037 | 0.155556 |
| very high | 0 | 0 | 0 | 0 | 0.052222 | 0 | 0 | 0 | 0 | 0.065278 |

| Chemical_Cate | very low | | | | | | | | | |
| --- | --- | --- | --- | --- | --- | --- | --- | --- | --- | --- |
| Tox_fish_chem | medium | | | | | | | | | |
| Tox_fish_oth_s | medium | | | | | high | | | | |
| Tox_emb_lev | very low | low | medium | high | very high | very low | low | medium | high | very high |
| very low | 0.296296 | 0.138889 | 0.111111 | 0.097222 | 0.087037 | 0.243056 | 0.12037 | 0.097222 | 0.083334 | 0.075 |
| low | 0.407407 | 0.472222 | 0.277778 | 0.236111 | 0.207407 | 0.354167 | 0.37963 | 0.236111 | 0.194444 | 0.172222 |
| medium | 0.296296 | 0.388889 | 0.611111 | 0.486111 | 0.411111 | 0.3125 | 0.37963 | 0.486111 | 0.361111 | 0.311111 |
| high | 0 | 0 | 0 | 0.180556 | 0.207407 | 0.090278 | 0.12037 | 0.180556 | 0.361111 | 0.311111 |
| very high | 0 | 0 | 0 | 0 | 0.087037 | 0 | 0 | 0 | 0 | 0.130556 |

| Chemical_Cate | very low | | | | | | | | | |
| --- | --- | --- | --- | --- | --- | --- | --- | --- | --- | --- |
| Tox_fish_chem | medium | | | | | high | | | | |
| Tox_fish_oth_s | very high | | | | | very low | | | | |
| Tox_emb_lev | very low | low | medium | high | very high | very low | low | medium | high | very high |
| very low | 0.207778 | 0.106944 | 0.087037 | 0.075 | 0.066667 | 0.520834 | 0.322917 | 0.243056 | 0.197917 | 0.168333 |
| low | 0.313333 | 0.322222 | 0.207407 | 0.172222 | 0.15 | 0.270833 | 0.427083 | 0.354167 | 0.302083 | 0.264167 |
| medium | 0.302222 | 0.35 | 0.411111 | 0.311111 | 0.261111 | 0.145833 | 0.177083 | 0.3125 | 0.302083 | 0.280833 |
| high | 0.124444 | 0.155556 | 0.207407 | 0.311111 | 0.261111 | 0.0625 | 0.072917 | 0.090278 | 0.197917 | 0.2225 |
| very high | 0.052222 | 0.065278 | 0.087037 | 0.130556 | 0.261111 | 0 | 0 | 0 | 0 | 0.064167 |

| Chemical_Cate | very low | | | | | | | | | |
| --- | --- | --- | --- | --- | --- | --- | --- | --- | --- | --- |
| Tox_fish_chem | high | | | | | | | | | |
| Tox_fish_oth_s | low | | | | | medium | | | | |
| Tox_emb_lev | very low | low | medium | high | very high | very low | low | medium | high | very high |
| very low | 0.322917 | 0.125 | 0.104167 | 0.090278 | 0.080208 | 0.243056 | 0.104167 | 0.083334 | 0.072917 | 0.065278 |
| low | 0.427083 | 0.583333 | 0.395833 | 0.3125 | 0.2625 | 0.354167 | 0.395833 | 0.208333 | 0.177083 | 0.155556 |
| medium | 0.177083 | 0.208333 | 0.395833 | 0.354167 | 0.314583 | 0.3125 | 0.395833 | 0.583333 | 0.427083 | 0.35 |
| high | 0.072917 | 0.083334 | 0.104167 | 0.243056 | 0.2625 | 0.090278 | 0.104167 | 0.125 | 0.322917 | 0.322222 |
| very high | 0 | 0 | 0 | 0 | 0.080208 | 0 | 0 | 0 | 0 | 0.106944 |

| Chemical_Cate | very low | | | | | | | | | |
| --- | --- | --- | --- | --- | --- | --- | --- | --- | --- | --- |
| Tox_fish_chem | high | | | | | | | | | |
| Tox_fish_oth_s | high | | | | | very high | | | | |
| Tox_emb_lev | very low | low | medium | high | very high | very low | low | medium | high | very high |
| very low | 0.197917 | 0.090278 | 0.072917 | 0.0625 | 0.05625 | 0.168333 | 0.080208 | 0.065278 | 0.05625 | 0.05 |
| low | 0.302083 | 0.3125 | 0.177083 | 0.145833 | 0.129167 | 0.264167 | 0.2625 | 0.155556 | 0.129167 | 0.1125 |
| medium | 0.302083 | 0.354167 | 0.427083 | 0.270833 | 0.233333 | 0.280833 | 0.314583 | 0.35 | 0.233333 | 0.195833 |
| high | 0.197917 | 0.243056 | 0.322917 | 0.520834 | 0.420833 | 0.2225 | 0.2625 | 0.322222 | 0.420833 | 0.320833 |
| very high | 0 | 0 | 0 | 0 | 0.160417 | 0.064167 | 0.080208 | 0.106944 | 0.160417 | 0.320833 |

| Chemical_Cate | very low | | | | | | | | | |
| --- | --- | --- | --- | --- | --- | --- | --- | --- | --- | --- |
| Tox_fish_chem | very high | | | | | | | | | |
| Tox_fish_oth_s | very low | | | | | low | | | | |
| Tox_emb_lev | very low | low | medium | high | very high | very low | low | medium | high | very high |
| very low | 0.456667 | 0.278333 | 0.207778 | 0.168333 | 0.142667 | 0.278333 | 0.1 | 0.083333 | 0.072222 | 0.064167 |
| low | 0.256667 | 0.386667 | 0.313333 | 0.264167 | 0.229333 | 0.386667 | 0.516666 | 0.341667 | 0.266667 | 0.2225 |
| medium | 0.156667 | 0.186667 | 0.302222 | 0.280833 | 0.256 | 0.186667 | 0.216667 | 0.375 | 0.322222 | 0.280833 |
| high | 0.09 | 0.103333 | 0.124444 | 0.2225 | 0.229333 | 0.103333 | 0.116667 | 0.141667 | 0.266667 | 0.264167 |
| very high | 0.04 | 0.045 | 0.052222 | 0.064167 | 0.142667 | 0.045 | 0.05 | 0.058333 | 0.072222 | 0.168333 |

| Chemical_Cate | very low | | | | | | | | | |
| --- | --- | --- | --- | --- | --- | --- | --- | --- | --- | --- |
| Tox_fish_chem | very high | | | | | | | | | |
| Tox_fish_oth_s | medium | | | | | high | | | | |
| Tox_emb_lev | very low | low | medium | high | very high | very low | low | medium | high | very high |
| very low | 0.207778 | 0.083333 | 0.066667 | 0.058333 | 0.052222 | 0.168333 | 0.072222 | 0.058333 | 0.05 | 0.045 |
| low | 0.313333 | 0.341667 | 0.166667 | 0.141667 | 0.124444 | 0.264167 | 0.266667 | 0.141667 | 0.116667 | 0.103333 |
| medium | 0.302222 | 0.375 | 0.533333 | 0.375 | 0.302222 | 0.280833 | 0.322222 | 0.375 | 0.216667 | 0.186667 |
| high | 0.124444 | 0.141667 | 0.166667 | 0.341667 | 0.313333 | 0.2225 | 0.266667 | 0.341667 | 0.516666 | 0.386667 |
| very high | 0.052222 | 0.058333 | 0.066667 | 0.083333 | 0.207778 | 0.064167 | 0.072222 | 0.083333 | 0.1 | 0.278333 |

| Chemical_Cate | very low | | | | | low | | | | |
| --- | --- | --- | --- | --- | --- | --- | --- | --- | --- | --- |
| Tox_fish_chem | very high | | | | | very low | | | | |
| Tox_fish_oth_s | very high | | | | | very low | | | | |
| Tox_emb_lev | very low | low | medium | high | very high | very low | low | medium | high | very high |
| very low | 0.142667 | 0.064167 | 0.052222 | 0.045 | 0.04 | 0.75 | 0.5 | 0.388889 | 0.322917 | 0.278333 |
| low | 0.229333 | 0.2225 | 0.124444 | 0.103333 | 0.09 | 0.25 | 0.5 | 0.472222 | 0.427083 | 0.386667 |
| medium | 0.256 | 0.280833 | 0.302222 | 0.186667 | 0.156667 | 0 | 0 | 0.138889 | 0.177083 | 0.186667 |
| high | 0.229333 | 0.264167 | 0.313333 | 0.386667 | 0.256667 | 0 | 0 | 0 | 0.072917 | 0.103333 |
| very high | 0.142667 | 0.168333 | 0.207778 | 0.278333 | 0.456667 | 0 | 0 | 0 | 0 | 0.045 |

| Chemical_Cate | low | | | | | | | | | |
| --- | --- | --- | --- | --- | --- | --- | --- | --- | --- | --- |
| Tox_fish_chem | very low | | | | | | | | | |
| Tox_fish_oth_s | low | | | | | medium | | | | |
| Tox_emb_lev | very low | low | medium | high | very high | very low | low | medium | high | very high |
| very low | 0.5 | 0.25 | 0.208333 | 0.180556 | 0.160417 | 0.388889 | 0.208333 | 0.166667 | 0.145833 | 0.130556 |
| low | 0.5 | 0.75 | 0.583334 | 0.486111 | 0.420833 | 0.472222 | 0.583334 | 0.416667 | 0.354167 | 0.311111 |
| medium | 0 | 0 | 0.208333 | 0.236111 | 0.233333 | 0.138889 | 0.208333 | 0.416667 | 0.354167 | 0.311111 |
| high | 0 | 0 | 0 | 0.097222 | 0.129167 | 0 | 0 | 0 | 0.145833 | 0.172222 |
| very high | 0 | 0 | 0 | 0 | 0.05625 | 0 | 0 | 0 | 0 | 0.075 |

| Chemical_Cate | low | | | | | | | | | |
| --- | --- | --- | --- | --- | --- | --- | --- | --- | --- | --- |
| Tox_fish_chem | very low | | | | | | | | | |
| Tox_fish_oth_s | high | | | | | very high | | | | |
| Tox_emb_lev | very low | low | medium | high | very high | very low | low | medium | high | very high |
| very low | 0.322917 | 0.180556 | 0.145833 | 0.125 | 0.1125 | 0.278333 | 0.160417 | 0.130556 | 0.1125 | 0.1 |
| low | 0.427083 | 0.486111 | 0.354167 | 0.291667 | 0.258333 | 0.386667 | 0.420833 | 0.311111 | 0.258333 | 0.225 |
| medium | 0.177083 | 0.236111 | 0.354167 | 0.291667 | 0.258333 | 0.186667 | 0.233333 | 0.311111 | 0.258333 | 0.225 |
| high | 0.072917 | 0.097222 | 0.145833 | 0.291667 | 0.258333 | 0.103333 | 0.129167 | 0.172222 | 0.258333 | 0.225 |
| very high | 0 | 0 | 0 | 0 | 0.1125 | 0.045 | 0.05625 | 0.075 | 0.1125 | 0.225 |

| Chemical_Cate | low | | | | | | | | | |
| --- | --- | --- | --- | --- | --- | --- | --- | --- | --- | --- |
| Tox_fish_chem | low | | | | | | | | | |
| Tox_fish_oth_s | very low | | | | | low | | | | |
| Tox_emb_lev | very low | low | medium | high | very high | very low | low | medium | high | very high |
| very low | 0.5 | 0.25 | 0.166667 | 0.125 | 0.1 | 0.25 | 0 | 0 | 0 | 0 |
| low | 0.5 | 0.75 | 0.666666 | 0.583333 | 0.516666 | 0.75 | 1 | 0.75 | 0.611111 | 0.520834 |
| medium | 0 | 0 | 0.166667 | 0.208333 | 0.216667 | 0 | 0 | 0.25 | 0.277778 | 0.270833 |
| high | 0 | 0 | 0 | 0.083334 | 0.116667 | 0 | 0 | 0 | 0.111111 | 0.145833 |
| very high | 0 | 0 | 0 | 0 | 0.05 | 0 | 0 | 0 | 0 | 0.0625 |

| Chemical_Cate | low | | | | | | | | | |
| --- | --- | --- | --- | --- | --- | --- | --- | --- | --- | --- |
| Tox_fish_chem | low | | | | | | | | | |
| Tox_fish_oth_s | medium | | | | | high | | | | |
| Tox_emb_lev | very low | low | medium | high | very high | very low | low | medium | high | very high |
| very low | 0.166667 | 0 | 0 | 0 | 0 | 0.125 | 0 | 0 | 0 | 0 |
| low | 0.666666 | 0.75 | 0.5 | 0.416667 | 0.361111 | 0.583333 | 0.611111 | 0.416667 | 0.333333 | 0.291667 |
| medium | 0.166667 | 0.25 | 0.5 | 0.416667 | 0.361111 | 0.208333 | 0.277778 | 0.416667 | 0.333333 | 0.291667 |
| high | 0 | 0 | 0 | 0.166667 | 0.194444 | 0.083334 | 0.111111 | 0.166667 | 0.333333 | 0.291667 |
| very high | 0 | 0 | 0 | 0 | 0.083334 | 0 | 0 | 0 | 0 | 0.125 |

| Chemical_Cate | low | | | | | | | | | |
| --- | --- | --- | --- | --- | --- | --- | --- | --- | --- | --- |
| Tox_fish_chem | low | | | | | medium | | | | |
| Tox_fish_oth_s | very high | | | | | very low | | | | |
| Tox_emb_lev | very low | low | medium | high | very high | very low | low | medium | high | very high |
| very low | 0.1 | 0 | 0 | 0 | 0 | 0.416667 | 0.208333 | 0.138889 | 0.104167 | 0.083333 |
| low | 0.516666 | 0.520834 | 0.361111 | 0.291667 | 0.25 | 0.416667 | 0.583334 | 0.472222 | 0.395833 | 0.341667 |
| medium | 0.216667 | 0.270833 | 0.361111 | 0.291667 | 0.25 | 0.166667 | 0.208333 | 0.388889 | 0.395833 | 0.375 |
| high | 0.116667 | 0.145833 | 0.194444 | 0.291667 | 0.25 | 0 | 0 | 0 | 0.104167 | 0.141667 |
| very high | 0.05 | 0.0625 | 0.083334 | 0.125 | 0.25 | 0 | 0 | 0 | 0 | 0.058333 |

| Chemical_Cate | low | | | | | | | | | |
| --- | --- | --- | --- | --- | --- | --- | --- | --- | --- | --- |
| Tox_fish_chem | medium | | | | | | | | | |
| Tox_fish_oth_s | low | | | | | medium | | | | |
| Tox_emb_lev | very low | low | medium | high | very high | very low | low | medium | high | very high |
| very low | 0.208333 | 0 | 0 | 0 | 0 | 0.138889 | 0 | 0 | 0 | 0 |
| low | 0.583334 | 0.75 | 0.5 | 0.388889 | 0.322917 | 0.472222 | 0.5 | 0.25 | 0.208333 | 0.180556 |
| medium | 0.208333 | 0.25 | 0.5 | 0.472222 | 0.427083 | 0.388889 | 0.5 | 0.75 | 0.583334 | 0.486111 |
| high | 0 | 0 | 0 | 0.138889 | 0.177083 | 0 | 0 | 0 | 0.208333 | 0.236111 |
| very high | 0 | 0 | 0 | 0 | 0.072917 | 0 | 0 | 0 | 0 | 0.097222 |

| Chemical_Cate | low | | | | | | | | | |
| --- | --- | --- | --- | --- | --- | --- | --- | --- | --- | --- |
| Tox_fish_chem | medium | | | | | | | | | |
| Tox_fish_oth_s | high | | | | | very high | | | | |
| Tox_emb_lev | very low | low | medium | high | very high | very low | low | medium | high | very high |
| very low | 0.104167 | 0 | 0 | 0 | 0 | 0.083333 | 0 | 0 | 0 | 0 |
| low | 0.395833 | 0.388889 | 0.208333 | 0.166667 | 0.145833 | 0.341667 | 0.322917 | 0.180556 | 0.145833 | 0.125 |
| medium | 0.395833 | 0.472222 | 0.583334 | 0.416667 | 0.354167 | 0.375 | 0.427083 | 0.486111 | 0.354167 | 0.291667 |
| high | 0.104167 | 0.138889 | 0.208333 | 0.416667 | 0.354167 | 0.141667 | 0.177083 | 0.236111 | 0.354167 | 0.291667 |
| very high | 0 | 0 | 0 | 0 | 0.145833 | 0.058333 | 0.072917 | 0.097222 | 0.145833 | 0.291667 |

| Chemical_Cate | low | | | | | | | | | |
| --- | --- | --- | --- | --- | --- | --- | --- | --- | --- | --- |
| Tox_fish_chem | high | | | | | | | | | |
| Tox_fish_oth_s | very low | | | | | low | | | | |
| Tox_emb_lev | very low | low | medium | high | very high | very low | low | medium | high | very high |
| very low | 0.361111 | 0.180556 | 0.12037 | 0.090278 | 0.072222 | 0.180556 | 0 | 0 | 0 | 0 |
| low | 0.361111 | 0.486111 | 0.37963 | 0.3125 | 0.266667 | 0.486111 | 0.611111 | 0.388889 | 0.296296 | 0.243056 |
| medium | 0.194444 | 0.236111 | 0.37963 | 0.354167 | 0.322222 | 0.236111 | 0.277778 | 0.472222 | 0.407407 | 0.354167 |
| high | 0.083334 | 0.097222 | 0.12037 | 0.243056 | 0.266667 | 0.097222 | 0.111111 | 0.138889 | 0.296296 | 0.3125 |
| very high | 0 | 0 | 0 | 0 | 0.072222 | 0 | 0 | 0 | 0 | 0.090278 |

| Chemical_Cate | low | | | | | | | | | |
| --- | --- | --- | --- | --- | --- | --- | --- | --- | --- | --- |
| Tox_fish_chem | high | | | | | | | | | |
| Tox_fish_oth_s | medium | | | | | high | | | | |
| Tox_emb_lev | very low | low | medium | high | very high | very low | low | medium | high | very high |
| very low | 0.12037 | 0 | 0 | 0 | 0 | 0.090278 | 0 | 0 | 0 | 0 |
| low | 0.37963 | 0.388889 | 0.166667 | 0.138889 | 0.12037 | 0.3125 | 0.296296 | 0.138889 | 0.111111 | 0.097222 |
| medium | 0.37963 | 0.472222 | 0.666666 | 0.472222 | 0.37963 | 0.354167 | 0.407407 | 0.472222 | 0.277778 | 0.236111 |
| high | 0.12037 | 0.138889 | 0.166667 | 0.388889 | 0.37963 | 0.243056 | 0.296296 | 0.388889 | 0.611111 | 0.486111 |
| very high | 0 | 0 | 0 | 0 | 0.12037 | 0 | 0 | 0 | 0 | 0.180556 |

| Chemical_Cate | low | | | | | | | | | |
| --- | --- | --- | --- | --- | --- | --- | --- | --- | --- | --- |
| Tox_fish_chem | high | | | | | very high | | | | |
| Tox_fish_oth_s | very high | | | | | very low | | | | |
| Tox_emb_lev | very low | low | medium | high | very high | very low | low | medium | high | very high |
| very low | 0.072222 | 0 | 0 | 0 | 0 | 0.320833 | 0.160417 | 0.106944 | 0.080208 | 0.064167 |
| low | 0.266667 | 0.243056 | 0.12037 | 0.097222 | 0.083334 | 0.320833 | 0.420833 | 0.322222 | 0.2625 | 0.2225 |
| medium | 0.322222 | 0.354167 | 0.37963 | 0.236111 | 0.194444 | 0.195833 | 0.233333 | 0.35 | 0.314583 | 0.280833 |
| high | 0.266667 | 0.3125 | 0.37963 | 0.486111 | 0.361111 | 0.1125 | 0.129167 | 0.155556 | 0.2625 | 0.264167 |
| very high | 0.072222 | 0.090278 | 0.12037 | 0.180556 | 0.361111 | 0.05 | 0.05625 | 0.065278 | 0.080208 | 0.168333 |

| Chemical_Cate | low | | | | | | | | | |
| --- | --- | --- | --- | --- | --- | --- | --- | --- | --- | --- |
| Tox_fish_chem | very high | | | | | | | | | |
| Tox_fish_oth_s | low | | | | | medium | | | | |
| Tox_emb_lev | very low | low | medium | high | very high | very low | low | medium | high | very high |
| very low | 0.160417 | 0 | 0 | 0 | 0 | 0.106944 | 0 | 0 | 0 | 0 |
| low | 0.420833 | 0.520834 | 0.322917 | 0.243056 | 0.197917 | 0.322222 | 0.322917 | 0.125 | 0.104167 | 0.090278 |
| medium | 0.233333 | 0.270833 | 0.427083 | 0.354167 | 0.302083 | 0.35 | 0.427083 | 0.583333 | 0.395833 | 0.3125 |
| high | 0.129167 | 0.145833 | 0.177083 | 0.3125 | 0.302083 | 0.155556 | 0.177083 | 0.208333 | 0.395833 | 0.354167 |
| very high | 0.05625 | 0.0625 | 0.072917 | 0.090278 | 0.197917 | 0.065278 | 0.072917 | 0.083334 | 0.104167 | 0.243056 |

| Chemical_Cate | low | | | | | | | | | |
| --- | --- | --- | --- | --- | --- | --- | --- | --- | --- | --- |
| Tox_fish_chem | very high | | | | | | | | | |
| Tox_fish_oth_s | high | | | | | very high | | | | |
| Tox_emb_lev | very low | low | medium | high | very high | very low | low | medium | high | very high |
| very low | 0.080208 | 0 | 0 | 0 | 0 | 0.064167 | 0 | 0 | 0 | 0 |
| low | 0.2625 | 0.243056 | 0.104167 | 0.083334 | 0.072917 | 0.2225 | 0.197917 | 0.090278 | 0.072917 | 0.0625 |
| medium | 0.314583 | 0.354167 | 0.395833 | 0.208333 | 0.177083 | 0.280833 | 0.302083 | 0.3125 | 0.177083 | 0.145833 |
| high | 0.2625 | 0.3125 | 0.395833 | 0.583333 | 0.427083 | 0.264167 | 0.302083 | 0.354167 | 0.427083 | 0.270833 |
| very high | 0.080208 | 0.090278 | 0.104167 | 0.125 | 0.322917 | 0.168333 | 0.197917 | 0.243056 | 0.322917 | 0.520834 |

| Chemical_Cate | medium | | | | | | | | | |
| --- | --- | --- | --- | --- | --- | --- | --- | --- | --- | --- |
| Tox_fish_chem | very low | | | | | | | | | |
| Tox_fish_oth_s | very low | | | | | low | | | | |
| Tox_emb_lev | very low | low | medium | high | very high | very low | low | medium | high | very high |
| very low | 0.611111 | 0.388889 | 0.296296 | 0.243056 | 0.207778 | 0.388889 | 0.166667 | 0.138889 | 0.12037 | 0.106944 |
| low | 0.277778 | 0.472222 | 0.407407 | 0.354167 | 0.313333 | 0.472222 | 0.666666 | 0.472222 | 0.37963 | 0.322222 |
| medium | 0.111111 | 0.138889 | 0.296296 | 0.3125 | 0.302222 | 0.138889 | 0.166667 | 0.388889 | 0.37963 | 0.35 |
| high | 0 | 0 | 0 | 0.090278 | 0.124444 | 0 | 0 | 0 | 0.12037 | 0.155556 |
| very high | 0 | 0 | 0 | 0 | 0.052222 | 0 | 0 | 0 | 0 | 0.065278 |

| Chemical_Cate | medium | | | | | | | | | |
| --- | --- | --- | --- | --- | --- | --- | --- | --- | --- | --- |
| Tox_fish_chem | very low | | | | | | | | | |
| Tox_fish_oth_s | medium | | | | | high | | | | |
| Tox_emb_lev | very low | low | medium | high | very high | very low | low | medium | high | very high |
| very low | 0.296296 | 0.138889 | 0.111111 | 0.097222 | 0.087037 | 0.243056 | 0.12037 | 0.097222 | 0.083334 | 0.075 |
| low | 0.407407 | 0.472222 | 0.277778 | 0.236111 | 0.207407 | 0.354167 | 0.37963 | 0.236111 | 0.194444 | 0.172222 |
| medium | 0.296296 | 0.388889 | 0.611111 | 0.486111 | 0.411111 | 0.3125 | 0.37963 | 0.486111 | 0.361111 | 0.311111 |
| high | 0 | 0 | 0 | 0.180556 | 0.207407 | 0.090278 | 0.12037 | 0.180556 | 0.361111 | 0.311111 |
| very high | 0 | 0 | 0 | 0 | 0.087037 | 0 | 0 | 0 | 0 | 0.130556 |

| Chemical_Cate | medium | | | | | | | | | |
| --- | --- | --- | --- | --- | --- | --- | --- | --- | --- | --- |
| Tox_fish_chem | very low | | | | | low | | | | |
| Tox_fish_oth_s | very high | | | | | very low | | | | |
| Tox_emb_lev | very low | low | medium | high | very high | very low | low | medium | high | very high |
| very low | 0.207778 | 0.106944 | 0.087037 | 0.075 | 0.066667 | 0.416667 | 0.208333 | 0.138889 | 0.104167 | 0.083333 |
| low | 0.313333 | 0.322222 | 0.207407 | 0.172222 | 0.15 | 0.416667 | 0.583334 | 0.472222 | 0.395833 | 0.341667 |
| medium | 0.302222 | 0.35 | 0.411111 | 0.311111 | 0.261111 | 0.166667 | 0.208333 | 0.388889 | 0.395833 | 0.375 |
| high | 0.124444 | 0.155556 | 0.207407 | 0.311111 | 0.261111 | 0 | 0 | 0 | 0.104167 | 0.141667 |
| very high | 0.052222 | 0.065278 | 0.087037 | 0.130556 | 0.261111 | 0 | 0 | 0 | 0 | 0.058333 |

| Chemical_Cate | medium | | | | | | | | | |
| --- | --- | --- | --- | --- | --- | --- | --- | --- | --- | --- |
| Tox_fish_chem | low | | | | | | | | | |
| Tox_fish_oth_s | low | | | | | medium | | | | |
| Tox_emb_lev | very low | low | medium | high | very high | very low | low | medium | high | very high |
| very low | 0.208333 | 0 | 0 | 0 | 0 | 0.138889 | 0 | 0 | 0 | 0 |
| low | 0.583334 | 0.75 | 0.5 | 0.388889 | 0.322917 | 0.472222 | 0.5 | 0.25 | 0.208333 | 0.180556 |
| medium | 0.208333 | 0.25 | 0.5 | 0.472222 | 0.427083 | 0.388889 | 0.5 | 0.75 | 0.583334 | 0.486111 |
| high | 0 | 0 | 0 | 0.138889 | 0.177083 | 0 | 0 | 0 | 0.208333 | 0.236111 |
| very high | 0 | 0 | 0 | 0 | 0.072917 | 0 | 0 | 0 | 0 | 0.097222 |

| Chemical_Cate | medium | | | | | | | | | |
| --- | --- | --- | --- | --- | --- | --- | --- | --- | --- | --- |
| Tox_fish_chem | low | | | | | | | | | |
| Tox_fish_oth_s | high | | | | | very high | | | | |
| Tox_emb_lev | very low | low | medium | high | very high | very low | low | medium | high | very high |
| very low | 0.104167 | 0 | 0 | 0 | 0 | 0.083333 | 0 | 0 | 0 | 0 |
| low | 0.395833 | 0.388889 | 0.208333 | 0.166667 | 0.145833 | 0.341667 | 0.322917 | 0.180556 | 0.145833 | 0.125 |
| medium | 0.395833 | 0.472222 | 0.583334 | 0.416667 | 0.354167 | 0.375 | 0.427083 | 0.486111 | 0.354167 | 0.291667 |
| high | 0.104167 | 0.138889 | 0.208333 | 0.416667 | 0.354167 | 0.141667 | 0.177083 | 0.236111 | 0.354167 | 0.291667 |
| very high | 0 | 0 | 0 | 0 | 0.145833 | 0.058333 | 0.072917 | 0.097222 | 0.145833 | 0.291667 |

| Chemical_Cate | medium | | | | | | | | | |
| --- | --- | --- | --- | --- | --- | --- | --- | --- | --- | --- |
| Tox_fish_chem | medium | | | | | | | | | |
| Tox_fish_oth_s | very low | | | | | low | | | | |
| Tox_emb_lev | very low | low | medium | high | very high | very low | low | medium | high | very high |
| very low | 0.333333 | 0.166667 | 0.111111 | 0.083334 | 0.066667 | 0.166667 | 0 | 0 | 0 | 0 |
| low | 0.333333 | 0.416667 | 0.277778 | 0.208333 | 0.166667 | 0.416667 | 0.5 | 0.25 | 0.166667 | 0.125 |
| medium | 0.333333 | 0.416667 | 0.611111 | 0.583333 | 0.533333 | 0.416667 | 0.5 | 0.75 | 0.666666 | 0.583333 |
| high | 0 | 0 | 0 | 0.125 | 0.166667 | 0 | 0 | 0 | 0.166667 | 0.208333 |
| very high | 0 | 0 | 0 | 0 | 0.066667 | 0 | 0 | 0 | 0 | 0.083334 |

| Chemical_Cate | medium | | | | | | | | | |
| --- | --- | --- | --- | --- | --- | --- | --- | --- | --- | --- |
| Tox_fish_chem | medium | | | | | | | | | |
| Tox_fish_oth_s | medium | | | | | high | | | | |
| Tox_emb_lev | very low | low | medium | high | very high | very low | low | medium | high | very high |
| very low | 0.111111 | 0 | 0 | 0 | 0 | 0.083334 | 0 | 0 | 0 | 0 |
| low | 0.277778 | 0.25 | 0 | 0 | 0 | 0.208333 | 0.166667 | 0 | 0 | 0 |
| medium | 0.611111 | 0.75 | 1 | 0.75 | 0.611111 | 0.583333 | 0.666666 | 0.75 | 0.5 | 0.416667 |
| high | 0 | 0 | 0 | 0.25 | 0.277778 | 0.125 | 0.166667 | 0.25 | 0.5 | 0.416667 |
| very high | 0 | 0 | 0 | 0 | 0.111111 | 0 | 0 | 0 | 0 | 0.166667 |

| Chemical_Cate | medium | | | | | | | | | |
| --- | --- | --- | --- | --- | --- | --- | --- | --- | --- | --- |
| Tox_fish_chem | medium | | | | | high | | | | |
| Tox_fish_oth_s | very high | | | | | very low | | | | |
| Tox_emb_lev | very low | low | medium | high | very high | very low | low | medium | high | very high |
| very low | 0.066667 | 0 | 0 | 0 | 0 | 0.291667 | 0.145833 | 0.097222 | 0.072917 | 0.058333 |
| low | 0.166667 | 0.125 | 0 | 0 | 0 | 0.291667 | 0.354167 | 0.236111 | 0.177083 | 0.141667 |
| medium | 0.533333 | 0.583333 | 0.611111 | 0.416667 | 0.333333 | 0.291667 | 0.354167 | 0.486111 | 0.427083 | 0.375 |
| high | 0.166667 | 0.208333 | 0.277778 | 0.416667 | 0.333333 | 0.125 | 0.145833 | 0.180556 | 0.322917 | 0.341667 |
| very high | 0.066667 | 0.083334 | 0.111111 | 0.166667 | 0.333333 | 0 | 0 | 0 | 0 | 0.083333 |

| Chemical_Cate | medium | | | | | | | | | |
| --- | --- | --- | --- | --- | --- | --- | --- | --- | --- | --- |
| Tox_fish_chem | high | | | | | | | | | |
| Tox_fish_oth_s | low | | | | | medium | | | | |
| Tox_emb_lev | very low | low | medium | high | very high | very low | low | medium | high | very high |
| very low | 0.145833 | 0 | 0 | 0 | 0 | 0.097222 | 0 | 0 | 0 | 0 |
| low | 0.354167 | 0.416667 | 0.208333 | 0.138889 | 0.104167 | 0.236111 | 0.208333 | 0 | 0 | 0 |
| medium | 0.354167 | 0.416667 | 0.583334 | 0.472222 | 0.395833 | 0.486111 | 0.583334 | 0.75 | 0.5 | 0.388889 |
| high | 0.145833 | 0.166667 | 0.208333 | 0.388889 | 0.395833 | 0.180556 | 0.208333 | 0.25 | 0.5 | 0.472222 |
| very high | 0 | 0 | 0 | 0 | 0.104167 | 0 | 0 | 0 | 0 | 0.138889 |

| Chemical_Cate | medium | | | | | | | | | |
| --- | --- | --- | --- | --- | --- | --- | --- | --- | --- | --- |
| Tox_fish_chem | high | | | | | | | | | |
| Tox_fish_oth_s | high | | | | | very high | | | | |
| Tox_emb_lev | very low | low | medium | high | very high | very low | low | medium | high | very high |
| very low | 0.072917 | 0 | 0 | 0 | 0 | 0.058333 | 0 | 0 | 0 | 0 |
| low | 0.177083 | 0.138889 | 0 | 0 | 0 | 0.141667 | 0.104167 | 0 | 0 | 0 |
| medium | 0.427083 | 0.472222 | 0.5 | 0.25 | 0.208333 | 0.375 | 0.395833 | 0.388889 | 0.208333 | 0.166667 |
| high | 0.322917 | 0.388889 | 0.5 | 0.75 | 0.583334 | 0.341667 | 0.395833 | 0.472222 | 0.583334 | 0.416667 |
| very high | 0 | 0 | 0 | 0 | 0.208333 | 0.083333 | 0.104167 | 0.138889 | 0.208333 | 0.416667 |

| Chemical_Cate | medium | | | | | | | | | |
| --- | --- | --- | --- | --- | --- | --- | --- | --- | --- | --- |
| Tox_fish_chem | very high | | | | | | | | | |
| Tox_fish_oth_s | very low | | | | | low | | | | |
| Tox_emb_lev | very low | low | medium | high | very high | very low | low | medium | high | very high |
| very low | 0.261111 | 0.130556 | 0.087037 | 0.065278 | 0.052222 | 0.130556 | 0 | 0 | 0 | 0 |
| low | 0.261111 | 0.311111 | 0.207407 | 0.155556 | 0.124444 | 0.311111 | 0.361111 | 0.180556 | 0.12037 | 0.090278 |
| medium | 0.261111 | 0.311111 | 0.411111 | 0.35 | 0.302222 | 0.311111 | 0.361111 | 0.486111 | 0.37963 | 0.3125 |
| high | 0.15 | 0.172222 | 0.207407 | 0.322222 | 0.313333 | 0.172222 | 0.194444 | 0.236111 | 0.37963 | 0.354167 |
| very high | 0.066667 | 0.075 | 0.087037 | 0.106944 | 0.207778 | 0.075 | 0.083334 | 0.097222 | 0.12037 | 0.243056 |

| Chemical_Cate | medium | | | | | | | | | |
| --- | --- | --- | --- | --- | --- | --- | --- | --- | --- | --- |
| Tox_fish_chem | very high | | | | | | | | | |
| Tox_fish_oth_s | medium | | | | | high | | | | |
| Tox_emb_lev | very low | low | medium | high | very high | very low | low | medium | high | very high |
| very low | 0.087037 | 0 | 0 | 0 | 0 | 0.065278 | 0 | 0 | 0 | 0 |
| low | 0.207407 | 0.180556 | 0 | 0 | 0 | 0.155556 | 0.12037 | 0 | 0 | 0 |
| medium | 0.411111 | 0.486111 | 0.611111 | 0.388889 | 0.296296 | 0.35 | 0.37963 | 0.388889 | 0.166667 | 0.138889 |
| high | 0.207407 | 0.236111 | 0.277778 | 0.472222 | 0.407407 | 0.322222 | 0.37963 | 0.472222 | 0.666666 | 0.472222 |
| very high | 0.087037 | 0.097222 | 0.111111 | 0.138889 | 0.296296 | 0.106944 | 0.12037 | 0.138889 | 0.166667 | 0.388889 |

| Chemical_Cate | medium | | | | | high | | | | |
| --- | --- | --- | --- | --- | --- | --- | --- | --- | --- | --- |
| Tox_fish_chem | very high | | | | | very low | | | | |
| Tox_fish_oth_s | very high | | | | | very low | | | | |
| Tox_emb_lev | very low | low | medium | high | very high | very low | low | medium | high | very high |
| very low | 0.052222 | 0 | 0 | 0 | 0 | 0.520834 | 0.322917 | 0.243056 | 0.197917 | 0.168333 |
| low | 0.124444 | 0.090278 | 0 | 0 | 0 | 0.270833 | 0.427083 | 0.354167 | 0.302083 | 0.264167 |
| medium | 0.302222 | 0.3125 | 0.296296 | 0.138889 | 0.111111 | 0.145833 | 0.177083 | 0.3125 | 0.302083 | 0.280833 |
| high | 0.313333 | 0.354167 | 0.407407 | 0.472222 | 0.277778 | 0.0625 | 0.072917 | 0.090278 | 0.197917 | 0.2225 |
| very high | 0.207778 | 0.243056 | 0.296296 | 0.388889 | 0.611111 | 0 | 0 | 0 | 0 | 0.064167 |

| Chemical_Cate | high | | | | | | | | | |
| --- | --- | --- | --- | --- | --- | --- | --- | --- | --- | --- |
| Tox_fish_chem | very low | | | | | | | | | |
| Tox_fish_oth_s | low | | | | | medium | | | | |
| Tox_emb_lev | very low | low | medium | high | very high | very low | low | medium | high | very high |
| very low | 0.322917 | 0.125 | 0.104167 | 0.090278 | 0.080208 | 0.243056 | 0.104167 | 0.083334 | 0.072917 | 0.065278 |
| low | 0.427083 | 0.583333 | 0.395833 | 0.3125 | 0.2625 | 0.354167 | 0.395833 | 0.208333 | 0.177083 | 0.155556 |
| medium | 0.177083 | 0.208333 | 0.395833 | 0.354167 | 0.314583 | 0.3125 | 0.395833 | 0.583333 | 0.427083 | 0.35 |
| high | 0.072917 | 0.083334 | 0.104167 | 0.243056 | 0.2625 | 0.090278 | 0.104167 | 0.125 | 0.322917 | 0.322222 |
| very high | 0 | 0 | 0 | 0 | 0.080208 | 0 | 0 | 0 | 0 | 0.106944 |

| Chemical_Cate | high | | | | | | | | | |
| --- | --- | --- | --- | --- | --- | --- | --- | --- | --- | --- |
| Tox_fish_chem | very low | | | | | | | | | |
| Tox_fish_oth_s | high | | | | | very high | | | | |
| Tox_emb_lev | very low | low | medium | high | very high | very low | low | medium | high | very high |
| very low | 0.197917 | 0.090278 | 0.072917 | 0.0625 | 0.05625 | 0.168333 | 0.080208 | 0.065278 | 0.05625 | 0.05 |
| low | 0.302083 | 0.3125 | 0.177083 | 0.145833 | 0.129167 | 0.264167 | 0.2625 | 0.155556 | 0.129167 | 0.1125 |
| medium | 0.302083 | 0.354167 | 0.427083 | 0.270833 | 0.233333 | 0.280833 | 0.314583 | 0.35 | 0.233333 | 0.195833 |
| high | 0.197917 | 0.243056 | 0.322917 | 0.520834 | 0.420833 | 0.2225 | 0.2625 | 0.322222 | 0.420833 | 0.320833 |
| very high | 0 | 0 | 0 | 0 | 0.160417 | 0.064167 | 0.080208 | 0.106944 | 0.160417 | 0.320833 |

| Chemical_Cate | high | | | | | | | | | |
| --- | --- | --- | --- | --- | --- | --- | --- | --- | --- | --- |
| Tox_fish_chem | low | | | | | | | | | |
| Tox_fish_oth_s | very low | | | | | low | | | | |
| Tox_emb_lev | very low | low | medium | high | very high | very low | low | medium | high | very high |
| very low | 0.361111 | 0.180556 | 0.12037 | 0.090278 | 0.072222 | 0.180556 | 0 | 0 | 0 | 0 |
| low | 0.361111 | 0.486111 | 0.37963 | 0.3125 | 0.266667 | 0.486111 | 0.611111 | 0.388889 | 0.296296 | 0.243056 |
| medium | 0.194444 | 0.236111 | 0.37963 | 0.354167 | 0.322222 | 0.236111 | 0.277778 | 0.472222 | 0.407407 | 0.354167 |
| high | 0.083334 | 0.097222 | 0.12037 | 0.243056 | 0.266667 | 0.097222 | 0.111111 | 0.138889 | 0.296296 | 0.3125 |
| very high | 0 | 0 | 0 | 0 | 0.072222 | 0 | 0 | 0 | 0 | 0.090278 |

| Chemical_Cate | high | | | | | | | | | |
| --- | --- | --- | --- | --- | --- | --- | --- | --- | --- | --- |
| Tox_fish_chem | low | | | | | | | | | |
| Tox_fish_oth_s | medium | | | | | high | | | | |
| Tox_emb_lev | very low | low | medium | high | very high | very low | low | medium | high | very high |
| very low | 0.12037 | 0 | 0 | 0 | 0 | 0.090278 | 0 | 0 | 0 | 0 |
| low | 0.37963 | 0.388889 | 0.166667 | 0.138889 | 0.12037 | 0.3125 | 0.296296 | 0.138889 | 0.111111 | 0.097222 |
| medium | 0.37963 | 0.472222 | 0.666666 | 0.472222 | 0.37963 | 0.354167 | 0.407407 | 0.472222 | 0.277778 | 0.236111 |
| high | 0.12037 | 0.138889 | 0.166667 | 0.388889 | 0.37963 | 0.243056 | 0.296296 | 0.388889 | 0.611111 | 0.486111 |
| very high | 0 | 0 | 0 | 0 | 0.12037 | 0 | 0 | 0 | 0 | 0.180556 |

| Chemical_Cate | high | | | | | | | | | |
| --- | --- | --- | --- | --- | --- | --- | --- | --- | --- | --- |
| Tox_fish_chem | low | | | | | medium | | | | |
| Tox_fish_oth_s | very high | | | | | very low | | | | |
| Tox_emb_lev | very low | low | medium | high | very high | very low | low | medium | high | very high |
| very low | 0.072222 | 0 | 0 | 0 | 0 | 0.291667 | 0.145833 | 0.097222 | 0.072917 | 0.058333 |
| low | 0.266667 | 0.243056 | 0.12037 | 0.097222 | 0.083334 | 0.291667 | 0.354167 | 0.236111 | 0.177083 | 0.141667 |
| medium | 0.322222 | 0.354167 | 0.37963 | 0.236111 | 0.194444 | 0.291667 | 0.354167 | 0.486111 | 0.427083 | 0.375 |
| high | 0.266667 | 0.3125 | 0.37963 | 0.486111 | 0.361111 | 0.125 | 0.145833 | 0.180556 | 0.322917 | 0.341667 |
| very high | 0.072222 | 0.090278 | 0.12037 | 0.180556 | 0.361111 | 0 | 0 | 0 | 0 | 0.083333 |

| Chemical_Cate | high | | | | | | | | | |
| --- | --- | --- | --- | --- | --- | --- | --- | --- | --- | --- |
| Tox_fish_chem | medium | | | | | | | | | |
| Tox_fish_oth_s | low | | | | | medium | | | | |
| Tox_emb_lev | very low | low | medium | high | very high | very low | low | medium | high | very high |
| very low | 0.145833 | 0 | 0 | 0 | 0 | 0.097222 | 0 | 0 | 0 | 0 |
| low | 0.354167 | 0.416667 | 0.208333 | 0.138889 | 0.104167 | 0.236111 | 0.208333 | 0 | 0 | 0 |
| medium | 0.354167 | 0.416667 | 0.583334 | 0.472222 | 0.395833 | 0.486111 | 0.583334 | 0.75 | 0.5 | 0.388889 |
| high | 0.145833 | 0.166667 | 0.208333 | 0.388889 | 0.395833 | 0.180556 | 0.208333 | 0.25 | 0.5 | 0.472222 |
| very high | 0 | 0 | 0 | 0 | 0.104167 | 0 | 0 | 0 | 0 | 0.138889 |

| Chemical_Cate | high | | | | | | | | | |
| --- | --- | --- | --- | --- | --- | --- | --- | --- | --- | --- |
| Tox_fish_chem | medium | | | | | | | | | |
| Tox_fish_oth_s | high | | | | | very high | | | | |
| Tox_emb_lev | very low | low | medium | high | very high | very low | low | medium | high | very high |
| very low | 0.072917 | 0 | 0 | 0 | 0 | 0.058333 | 0 | 0 | 0 | 0 |
| low | 0.177083 | 0.138889 | 0 | 0 | 0 | 0.141667 | 0.104167 | 0 | 0 | 0 |
| medium | 0.427083 | 0.472222 | 0.5 | 0.25 | 0.208333 | 0.375 | 0.395833 | 0.388889 | 0.208333 | 0.166667 |
| high | 0.322917 | 0.388889 | 0.5 | 0.75 | 0.583334 | 0.341667 | 0.395833 | 0.472222 | 0.583334 | 0.416667 |
| very high | 0 | 0 | 0 | 0 | 0.208333 | 0.083333 | 0.104167 | 0.138889 | 0.208333 | 0.416667 |

| Chemical_Cate | high | | | | | | | | | |
| --- | --- | --- | --- | --- | --- | --- | --- | --- | --- | --- |
| Tox_fish_chem | high | | | | | | | | | |
| Tox_fish_oth_s | very low | | | | | low | | | | |
| Tox_emb_lev | very low | low | medium | high | very high | very low | low | medium | high | very high |
| very low | 0.25 | 0.125 | 0.083334 | 0.0625 | 0.05 | 0.125 | 0 | 0 | 0 | 0 |
| low | 0.25 | 0.291667 | 0.194444 | 0.145833 | 0.116667 | 0.291667 | 0.333333 | 0.166667 | 0.111111 | 0.083334 |
| medium | 0.25 | 0.291667 | 0.361111 | 0.270833 | 0.216667 | 0.291667 | 0.333333 | 0.416667 | 0.277778 | 0.208333 |
| high | 0.25 | 0.291667 | 0.361111 | 0.520834 | 0.516666 | 0.291667 | 0.333333 | 0.416667 | 0.611111 | 0.583333 |
| very high | 0 | 0 | 0 | 0 | 0.1 | 0 | 0 | 0 | 0 | 0.125 |

| Chemical_Cate | high | | | | | | | | | |
| --- | --- | --- | --- | --- | --- | --- | --- | --- | --- | --- |
| Tox_fish_chem | high | | | | | | | | | |
| Tox_fish_oth_s | medium | | | | | high | | | | |
| Tox_emb_lev | very low | low | medium | high | very high | very low | low | medium | high | very high |
| very low | 0.083334 | 0 | 0 | 0 | 0 | 0.0625 | 0 | 0 | 0 | 0 |
| low | 0.194444 | 0.166667 | 0 | 0 | 0 | 0.145833 | 0.111111 | 0 | 0 | 0 |
| medium | 0.361111 | 0.416667 | 0.5 | 0.25 | 0.166667 | 0.270833 | 0.277778 | 0.25 | 0 | 0 |
| high | 0.361111 | 0.416667 | 0.5 | 0.75 | 0.666666 | 0.520834 | 0.611111 | 0.75 | 1 | 0.75 |
| very high | 0 | 0 | 0 | 0 | 0.166667 | 0 | 0 | 0 | 0 | 0.25 |

| Chemical_Cate | high | | | | | | | | | |
| --- | --- | --- | --- | --- | --- | --- | --- | --- | --- | --- |
| Tox_fish_chem | high | | | | | very high | | | | |
| Tox_fish_oth_s | very high | | | | | very low | | | | |
| Tox_emb_lev | very low | low | medium | high | very high | very low | low | medium | high | very high |
| very low | 0.05 | 0 | 0 | 0 | 0 | 0.225 | 0.1125 | 0.075 | 0.05625 | 0.045 |
| low | 0.116667 | 0.083334 | 0 | 0 | 0 | 0.225 | 0.258333 | 0.172222 | 0.129167 | 0.103333 |
| medium | 0.216667 | 0.208333 | 0.166667 | 0 | 0 | 0.225 | 0.258333 | 0.311111 | 0.233333 | 0.186667 |
| high | 0.516666 | 0.583333 | 0.666666 | 0.75 | 0.5 | 0.225 | 0.258333 | 0.311111 | 0.420833 | 0.386667 |
| very high | 0.1 | 0.125 | 0.166667 | 0.25 | 0.5 | 0.1 | 0.1125 | 0.130556 | 0.160417 | 0.278333 |

| Chemical_Cate | high | | | | | | | | | |
| --- | --- | --- | --- | --- | --- | --- | --- | --- | --- | --- |
| Tox_fish_chem | very high | | | | | | | | | |
| Tox_fish_oth_s | low | | | | | medium | | | | |
| Tox_emb_lev | very low | low | medium | high | very high | very low | low | medium | high | very high |
| very low | 0.1125 | 0 | 0 | 0 | 0 | 0.075 | 0 | 0 | 0 | 0 |
| low | 0.258333 | 0.291667 | 0.145833 | 0.097222 | 0.072917 | 0.172222 | 0.145833 | 0 | 0 | 0 |
| medium | 0.258333 | 0.291667 | 0.354167 | 0.236111 | 0.177083 | 0.311111 | 0.354167 | 0.416667 | 0.208333 | 0.138889 |
| high | 0.258333 | 0.291667 | 0.354167 | 0.486111 | 0.427083 | 0.311111 | 0.354167 | 0.416667 | 0.583334 | 0.472222 |
| very high | 0.1125 | 0.125 | 0.145833 | 0.180556 | 0.322917 | 0.130556 | 0.145833 | 0.166667 | 0.208333 | 0.388889 |

| Chemical_Cate | high | | | | | | | | | |
| --- | --- | --- | --- | --- | --- | --- | --- | --- | --- | --- |
| Tox_fish_chem | very high | | | | | | | | | |
| Tox_fish_oth_s | high | | | | | very high | | | | |
| Tox_emb_lev | very low | low | medium | high | very high | very low | low | medium | high | very high |
| very low | 0.05625 | 0 | 0 | 0 | 0 | 0.045 | 0 | 0 | 0 | 0 |
| low | 0.129167 | 0.097222 | 0 | 0 | 0 | 0.103333 | 0.072917 | 0 | 0 | 0 |
| medium | 0.233333 | 0.236111 | 0.208333 | 0 | 0 | 0.186667 | 0.177083 | 0.138889 | 0 | 0 |
| high | 0.420833 | 0.486111 | 0.583334 | 0.75 | 0.5 | 0.386667 | 0.427083 | 0.472222 | 0.5 | 0.25 |
| very high | 0.160417 | 0.180556 | 0.208333 | 0.25 | 0.5 | 0.278333 | 0.322917 | 0.388889 | 0.5 | 0.75 |

| Chemical_Cate | very high | | | | | | | | | |
| --- | --- | --- | --- | --- | --- | --- | --- | --- | --- | --- |
| Tox_fish_chem | very low | | | | | | | | | |
| Tox_fish_oth_s | very low | | | | | low | | | | |
| Tox_emb_lev | very low | low | medium | high | very high | very low | low | medium | high | very high |
| very low | 0.456667 | 0.278333 | 0.207778 | 0.168333 | 0.142667 | 0.278333 | 0.1 | 0.083333 | 0.072222 | 0.064167 |
| low | 0.256667 | 0.386667 | 0.313333 | 0.264167 | 0.229333 | 0.386667 | 0.516666 | 0.341667 | 0.266667 | 0.2225 |
| medium | 0.156667 | 0.186667 | 0.302222 | 0.280833 | 0.256 | 0.186667 | 0.216667 | 0.375 | 0.322222 | 0.280833 |
| high | 0.09 | 0.103333 | 0.124444 | 0.2225 | 0.229333 | 0.103333 | 0.116667 | 0.141667 | 0.266667 | 0.264167 |
| very high | 0.04 | 0.045 | 0.052222 | 0.064167 | 0.142667 | 0.045 | 0.05 | 0.058333 | 0.072222 | 0.168333 |

| Chemical_Cate | very high | | | | | | | | | |
| --- | --- | --- | --- | --- | --- | --- | --- | --- | --- | --- |
| Tox_fish_chem | very low | | | | | | | | | |
| Tox_fish_oth_s | medium | | | | | high | | | | |
| Tox_emb_lev | very low | low | medium | high | very high | very low | low | medium | high | very high |
| very low | 0.207778 | 0.083333 | 0.066667 | 0.058333 | 0.052222 | 0.168333 | 0.072222 | 0.058333 | 0.05 | 0.045 |
| low | 0.313333 | 0.341667 | 0.166667 | 0.141667 | 0.124444 | 0.264167 | 0.266667 | 0.141667 | 0.116667 | 0.103333 |
| medium | 0.302222 | 0.375 | 0.533333 | 0.375 | 0.302222 | 0.280833 | 0.322222 | 0.375 | 0.216667 | 0.186667 |
| high | 0.124444 | 0.141667 | 0.166667 | 0.341667 | 0.313333 | 0.2225 | 0.266667 | 0.341667 | 0.516666 | 0.386667 |
| very high | 0.052222 | 0.058333 | 0.066667 | 0.083333 | 0.207778 | 0.064167 | 0.072222 | 0.083333 | 0.1 | 0.278333 |

| Chemical_Cate | very high | | | | | | | | | |
| --- | --- | --- | --- | --- | --- | --- | --- | --- | --- | --- |
| Tox_fish_chem | very low | | | | | low | | | | |
| Tox_fish_oth_s | very high | | | | | very low | | | | |
| Tox_emb_lev | very low | low | medium | high | very high | very low | low | medium | high | very high |
| very low | 0.142667 | 0.064167 | 0.052222 | 0.045 | 0.04 | 0.320833 | 0.160417 | 0.106944 | 0.080208 | 0.064167 |
| low | 0.229333 | 0.2225 | 0.124444 | 0.103333 | 0.09 | 0.320833 | 0.420833 | 0.322222 | 0.2625 | 0.2225 |
| medium | 0.256 | 0.280833 | 0.302222 | 0.186667 | 0.156667 | 0.195833 | 0.233333 | 0.35 | 0.314583 | 0.280833 |
| high | 0.229333 | 0.264167 | 0.313333 | 0.386667 | 0.256667 | 0.1125 | 0.129167 | 0.155556 | 0.2625 | 0.264167 |
| very high | 0.142667 | 0.168333 | 0.207778 | 0.278333 | 0.456667 | 0.05 | 0.05625 | 0.065278 | 0.080208 | 0.168333 |

| Chemical_Cate | very high | | | | | | | | | |
| --- | --- | --- | --- | --- | --- | --- | --- | --- | --- | --- |
| Tox_fish_chem | low | | | | | | | | | |
| Tox_fish_oth_s | low | | | | | medium | | | | |
| Tox_emb_lev | very low | low | medium | high | very high | very low | low | medium | high | very high |
| very low | 0.160417 | 0 | 0 | 0 | 0 | 0.106944 | 0 | 0 | 0 | 0 |
| low | 0.420833 | 0.520834 | 0.322917 | 0.243056 | 0.197917 | 0.322222 | 0.322917 | 0.125 | 0.104167 | 0.090278 |
| medium | 0.233333 | 0.270833 | 0.427083 | 0.354167 | 0.302083 | 0.35 | 0.427083 | 0.583333 | 0.395833 | 0.3125 |
| high | 0.129167 | 0.145833 | 0.177083 | 0.3125 | 0.302083 | 0.155556 | 0.177083 | 0.208333 | 0.395833 | 0.354167 |
| very high | 0.05625 | 0.0625 | 0.072917 | 0.090278 | 0.197917 | 0.065278 | 0.072917 | 0.083334 | 0.104167 | 0.243056 |

| Chemical_Cate | very high | | | | | | | | | |
| --- | --- | --- | --- | --- | --- | --- | --- | --- | --- | --- |
| Tox_fish_chem | low | | | | | | | | | |
| Tox_fish_oth_s | high | | | | | very high | | | | |
| Tox_emb_lev | very low | low | medium | high | very high | very low | low | medium | high | very high |
| very low | 0.080208 | 0 | 0 | 0 | 0 | 0.064167 | 0 | 0 | 0 | 0 |
| low | 0.2625 | 0.243056 | 0.104167 | 0.083334 | 0.072917 | 0.2225 | 0.197917 | 0.090278 | 0.072917 | 0.0625 |
| medium | 0.314583 | 0.354167 | 0.395833 | 0.208333 | 0.177083 | 0.280833 | 0.302083 | 0.3125 | 0.177083 | 0.145833 |
| high | 0.2625 | 0.3125 | 0.395833 | 0.583333 | 0.427083 | 0.264167 | 0.302083 | 0.354167 | 0.427083 | 0.270833 |
| very high | 0.080208 | 0.090278 | 0.104167 | 0.125 | 0.322917 | 0.168333 | 0.197917 | 0.243056 | 0.322917 | 0.520834 |

| Chemical_Cate | very high | | | | | | | | | |
| --- | --- | --- | --- | --- | --- | --- | --- | --- | --- | --- |
| Tox_fish_chem | medium | | | | | | | | | |
| Tox_fish_oth_s | very low | | | | | low | | | | |
| Tox_emb_lev | very low | low | medium | high | very high | very low | low | medium | high | very high |
| very low | 0.261111 | 0.130556 | 0.087037 | 0.065278 | 0.052222 | 0.130556 | 0 | 0 | 0 | 0 |
| low | 0.261111 | 0.311111 | 0.207407 | 0.155556 | 0.124444 | 0.311111 | 0.361111 | 0.180556 | 0.12037 | 0.090278 |
| medium | 0.261111 | 0.311111 | 0.411111 | 0.35 | 0.302222 | 0.311111 | 0.361111 | 0.486111 | 0.37963 | 0.3125 |
| high | 0.15 | 0.172222 | 0.207407 | 0.322222 | 0.313333 | 0.172222 | 0.194444 | 0.236111 | 0.37963 | 0.354167 |
| very high | 0.066667 | 0.075 | 0.087037 | 0.106944 | 0.207778 | 0.075 | 0.083334 | 0.097222 | 0.12037 | 0.243056 |

| Chemical_Cate | very high | | | | | | | | | |
| --- | --- | --- | --- | --- | --- | --- | --- | --- | --- | --- |
| Tox_fish_chem | medium | | | | | | | | | |
| Tox_fish_oth_s | medium | | | | | high | | | | |
| Tox_emb_lev | very low | low | medium | high | very high | very low | low | medium | high | very high |
| very low | 0.087037 | 0 | 0 | 0 | 0 | 0.065278 | 0 | 0 | 0 | 0 |
| low | 0.207407 | 0.180556 | 0 | 0 | 0 | 0.155556 | 0.12037 | 0 | 0 | 0 |
| medium | 0.411111 | 0.486111 | 0.611111 | 0.388889 | 0.296296 | 0.35 | 0.37963 | 0.388889 | 0.166667 | 0.138889 |
| high | 0.207407 | 0.236111 | 0.277778 | 0.472222 | 0.407407 | 0.322222 | 0.37963 | 0.472222 | 0.666666 | 0.472222 |
| very high | 0.087037 | 0.097222 | 0.111111 | 0.138889 | 0.296296 | 0.106944 | 0.12037 | 0.138889 | 0.166667 | 0.388889 |

| Chemical_Cate | very high | | | | | | | | | |
| --- | --- | --- | --- | --- | --- | --- | --- | --- | --- | --- |
| Tox_fish_chem | medium | | | | | high | | | | |
| Tox_fish_oth_s | very high | | | | | very low | | | | |
| Tox_emb_lev | very low | low | medium | high | very high | very low | low | medium | high | very high |
| very low | 0.052222 | 0 | 0 | 0 | 0 | 0.225 | 0.1125 | 0.075 | 0.05625 | 0.045 |
| low | 0.124444 | 0.090278 | 0 | 0 | 0 | 0.225 | 0.258333 | 0.172222 | 0.129167 | 0.103333 |
| medium | 0.302222 | 0.3125 | 0.296296 | 0.138889 | 0.111111 | 0.225 | 0.258333 | 0.311111 | 0.233333 | 0.186667 |
| high | 0.313333 | 0.354167 | 0.407407 | 0.472222 | 0.277778 | 0.225 | 0.258333 | 0.311111 | 0.420833 | 0.386667 |
| very high | 0.207778 | 0.243056 | 0.296296 | 0.388889 | 0.611111 | 0.1 | 0.1125 | 0.130556 | 0.160417 | 0.278333 |

| Chemical_Cate | very high | | | | | | | | | |
| --- | --- | --- | --- | --- | --- | --- | --- | --- | --- | --- |
| Tox_fish_chem | high | | | | | | | | | |
| Tox_fish_oth_s | low | | | | | medium | | | | |
| Tox_emb_lev | very low | low | medium | high | very high | very low | low | medium | high | very high |
| very low | 0.1125 | 0 | 0 | 0 | 0 | 0.075 | 0 | 0 | 0 | 0 |
| low | 0.258333 | 0.291667 | 0.145833 | 0.097222 | 0.072917 | 0.172222 | 0.145833 | 0 | 0 | 0 |
| medium | 0.258333 | 0.291667 | 0.354167 | 0.236111 | 0.177083 | 0.311111 | 0.354167 | 0.416667 | 0.208333 | 0.138889 |
| high | 0.258333 | 0.291667 | 0.354167 | 0.486111 | 0.427083 | 0.311111 | 0.354167 | 0.416667 | 0.583334 | 0.472222 |
| very high | 0.1125 | 0.125 | 0.145833 | 0.180556 | 0.322917 | 0.130556 | 0.145833 | 0.166667 | 0.208333 | 0.388889 |

| Chemical_Cate | very high | | | | | | | | | |
| --- | --- | --- | --- | --- | --- | --- | --- | --- | --- | --- |
| Tox_fish_chem | high | | | | | | | | | |
| Tox_fish_oth_s | high | | | | | very high | | | | |
| Tox_emb_lev | very low | low | medium | high | very high | very low | low | medium | high | very high |
| very low | 0.05625 | 0 | 0 | 0 | 0 | 0.045 | 0 | 0 | 0 | 0 |
| low | 0.129167 | 0.097222 | 0 | 0 | 0 | 0.103333 | 0.072917 | 0 | 0 | 0 |
| medium | 0.233333 | 0.236111 | 0.208333 | 0 | 0 | 0.186667 | 0.177083 | 0.138889 | 0 | 0 |
| high | 0.420833 | 0.486111 | 0.583334 | 0.75 | 0.5 | 0.386667 | 0.427083 | 0.472222 | 0.5 | 0.25 |
| very high | 0.160417 | 0.180556 | 0.208333 | 0.25 | 0.5 | 0.278333 | 0.322917 | 0.388889 | 0.5 | 0.75 |

| Chemical_Cate | very high | | | | | | | | | |
| --- | --- | --- | --- | --- | --- | --- | --- | --- | --- | --- |
| Tox_fish_chem | very high | | | | | | | | | |
| Tox_fish_oth_s | very low | | | | | low | | | | |
| Tox_emb_lev | very low | low | medium | high | very high | very low | low | medium | high | very high |
| very low | 0.2 | 0.1 | 0.066667 | 0.05 | 0.04 | 0.1 | 0 | 0 | 0 | 0 |
| low | 0.2 | 0.225 | 0.15 | 0.1125 | 0.09 | 0.225 | 0.25 | 0.125 | 0.083334 | 0.0625 |
| medium | 0.2 | 0.225 | 0.261111 | 0.195833 | 0.156667 | 0.225 | 0.25 | 0.291667 | 0.194444 | 0.145833 |
| high | 0.2 | 0.225 | 0.261111 | 0.320833 | 0.256667 | 0.225 | 0.25 | 0.291667 | 0.361111 | 0.270833 |
| very high | 0.2 | 0.225 | 0.261111 | 0.320833 | 0.456667 | 0.225 | 0.25 | 0.291667 | 0.361111 | 0.520834 |

| Chemical_Cate | very high | | | | | | | | | |
| --- | --- | --- | --- | --- | --- | --- | --- | --- | --- | --- |
| Tox_fish_chem | very high | | | | | | | | | |
| Tox_fish_oth_s | medium | | | | | high | | | | |
| Tox_emb_lev | very low | low | medium | high | very high | very low | low | medium | high | very high |
| very low | 0.066667 | 0 | 0 | 0 | 0 | 0.05 | 0 | 0 | 0 | 0 |
| low | 0.15 | 0.125 | 0 | 0 | 0 | 0.1125 | 0.083334 | 0 | 0 | 0 |
| medium | 0.261111 | 0.291667 | 0.333333 | 0.166667 | 0.111111 | 0.195833 | 0.194444 | 0.166667 | 0 | 0 |
| high | 0.261111 | 0.291667 | 0.333333 | 0.416667 | 0.277778 | 0.320833 | 0.361111 | 0.416667 | 0.5 | 0.25 |
| very high | 0.261111 | 0.291667 | 0.333333 | 0.416667 | 0.611111 | 0.320833 | 0.361111 | 0.416667 | 0.5 | 0.75 |

| Chemical_Cate | very high | | | | | unknown | | | | |
| --- | --- | --- | --- | --- | --- | --- | --- | --- | --- | --- |
| Tox_fish_chem | very high | | | | | very low | | | | |
| Tox_fish_oth_s | very high | | | | | very low | | | | |
| Tox_emb_lev | very low | low | medium | high | very high | very low | low | medium | high | very high |
| very low | 0.04 | 0 | 0 | 0 | 0 | 1 | 0.666667 | 0.5 | 0.4 | 0.2 |
| low | 0.09 | 0.0625 | 0 | 0 | 0 | 0 | 0.333333 | 0.25 | 0.2 | 0.2 |
| medium | 0.156667 | 0.145833 | 0.111111 | 0 | 0 | 0 | 0 | 0.25 | 0.2 | 0.2 |
| high | 0.256667 | 0.270833 | 0.277778 | 0.25 | 0 | 0 | 0 | 0 | 0.2 | 0.2 |
| very high | 0.456667 | 0.520834 | 0.611111 | 0.75 | 1 | 0 | 0 | 0 | 0 | 0.2 |

| Chemical_Cate | unknown | | | | | | | | | |
| --- | --- | --- | --- | --- | --- | --- | --- | --- | --- | --- |
| Tox_fish_chem | very low | | | | | | | | | |
| Tox_fish_oth_s | low | | | | | medium | | | | |
| Tox_emb_lev | very low | low | medium | high | very high | very low | low | medium | high | very high |
| very low | 0.666667 | 0.333333 | 0.333333 | 0.25 | 0.2 | 0.5 | 0.333333 | 0.25 | 0.25 | 0.2 |
| low | 0.333333 | 0.666667 | 0.333333 | 0.25 | 0.2 | 0.25 | 0.333333 | 0.25 | 0.25 | 0.2 |
| medium | 0 | 0 | 0.333333 | 0.25 | 0.2 | 0.25 | 0.333333 | 0.5 | 0.25 | 0.2 |
| high | 0 | 0 | 0 | 0.25 | 0.2 | 0 | 0 | 0 | 0.25 | 0.2 |
| very high | 0 | 0 | 0 | 0 | 0.2 | 0 | 0 | 0 | 0 | 0.2 |

| Chemical_Cate | unknown | | | | | | | | | |
| --- | --- | --- | --- | --- | --- | --- | --- | --- | --- | --- |
| Tox_fish_chem | very low | | | | | | | | | |
| Tox_fish_oth_s | high | | | | | very high | | | | |
| Tox_emb_lev | very low | low | medium | high | very high | very low | low | medium | high | very high |
| very low | 0.4 | 0.25 | 0.25 | 0.2 | 0.2 | 0.2 | 0.2 | 0.2 | 0.2 | 0.2 |
| low | 0.2 | 0.25 | 0.25 | 0.2 | 0.2 | 0.2 | 0.2 | 0.2 | 0.2 | 0.2 |
| medium | 0.2 | 0.25 | 0.25 | 0.2 | 0.2 | 0.2 | 0.2 | 0.2 | 0.2 | 0.2 |
| high | 0.2 | 0.25 | 0.25 | 0.4 | 0.2 | 0.2 | 0.2 | 0.2 | 0.2 | 0.2 |
| very high | 0 | 0 | 0 | 0 | 0.2 | 0.2 | 0.2 | 0.2 | 0.2 | 0.2 |

| Chemical_Cate | unknown | | | | | | | | | |
| --- | --- | --- | --- | --- | --- | --- | --- | --- | --- | --- |
| Tox_fish_chem | low | | | | | | | | | |
| Tox_fish_oth_s | very low | | | | | low | | | | |
| Tox_emb_lev | very low | low | medium | high | very high | very low | low | medium | high | very high |
| very low | 0.666667 | 0.333333 | 0.333333 | 0.25 | 0.2 | 0.333333 | 0 | 0 | 0 | 0 |
| low | 0.333333 | 0.666667 | 0.333333 | 0.25 | 0.2 | 0.666667 | 1 | 0.666667 | 0.5 | 0.4 |
| medium | 0 | 0 | 0.333333 | 0.25 | 0.2 | 0 | 0 | 0.333333 | 0.25 | 0.2 |
| high | 0 | 0 | 0 | 0.25 | 0.2 | 0 | 0 | 0 | 0.25 | 0.2 |
| very high | 0 | 0 | 0 | 0 | 0.2 | 0 | 0 | 0 | 0 | 0.2 |

| Chemical_Cate | unknown | | | | | | | | | |
| --- | --- | --- | --- | --- | --- | --- | --- | --- | --- | --- |
| Tox_fish_chem | low | | | | | | | | | |
| Tox_fish_oth_s | medium | | | | | high | | | | |
| Tox_emb_lev | very low | low | medium | high | very high | very low | low | medium | high | very high |
| very low | 0.333333 | 0 | 0 | 0 | 0 | 0.25 | 0 | 0 | 0 | 0 |
| low | 0.333333 | 0.666667 | 0.333333 | 0.333333 | 0.25 | 0.25 | 0.5 | 0.333333 | 0.25 | 0.25 |
| medium | 0.333333 | 0.333333 | 0.666667 | 0.333333 | 0.25 | 0.25 | 0.25 | 0.333333 | 0.25 | 0.25 |
| high | 0 | 0 | 0 | 0.333333 | 0.25 | 0.25 | 0.25 | 0.333333 | 0.5 | 0.25 |
| very high | 0 | 0 | 0 | 0 | 0.25 | 0 | 0 | 0 | 0 | 0.25 |

| Chemical_Cate | unknown | | | | | | | | | |
| --- | --- | --- | --- | --- | --- | --- | --- | --- | --- | --- |
| Tox_fish_chem | low | | | | | medium | | | | |
| Tox_fish_oth_s | very high | | | | | very low | | | | |
| Tox_emb_lev | very low | low | medium | high | very high | very low | low | medium | high | very high |
| very low | 0.2 | 0 | 0 | 0 | 0 | 0.5 | 0.333333 | 0.25 | 0.25 | 0.2 |
| low | 0.2 | 0.4 | 0.25 | 0.25 | 0.2 | 0.25 | 0.333333 | 0.25 | 0.25 | 0.2 |
| medium | 0.2 | 0.2 | 0.25 | 0.25 | 0.2 | 0.25 | 0.333333 | 0.5 | 0.25 | 0.2 |
| high | 0.2 | 0.2 | 0.25 | 0.25 | 0.2 | 0 | 0 | 0 | 0.25 | 0.2 |
| very high | 0.2 | 0.2 | 0.25 | 0.25 | 0.4 | 0 | 0 | 0 | 0 | 0.2 |

| Chemical_Cate | unknown | | | | | | | | | |
| --- | --- | --- | --- | --- | --- | --- | --- | --- | --- | --- |
| Tox_fish_chem | medium | | | | | | | | | |
| Tox_fish_oth_s | low | | | | | medium | | | | |
| Tox_emb_lev | very low | low | medium | high | very high | very low | low | medium | high | very high |
| very low | 0.333333 | 0 | 0 | 0 | 0 | 0.25 | 0 | 0 | 0 | 0 |
| low | 0.333333 | 0.666667 | 0.333333 | 0.333333 | 0.25 | 0.25 | 0.333333 | 0 | 0 | 0 |
| medium | 0.333333 | 0.333333 | 0.666667 | 0.333333 | 0.25 | 0.5 | 0.666667 | 1 | 0.666667 | 0.5 |
| high | 0 | 0 | 0 | 0.333333 | 0.25 | 0 | 0 | 0 | 0.333333 | 0.25 |
| very high | 0 | 0 | 0 | 0 | 0.25 | 0 | 0 | 0 | 0 | 0.25 |

| Chemical_Cate | unknown | | | | | | | | | |
| --- | --- | --- | --- | --- | --- | --- | --- | --- | --- | --- |
| Tox_fish_chem | medium | | | | | | | | | |
| Tox_fish_oth_s | high | | | | | very high | | | | |
| Tox_emb_lev | very low | low | medium | high | very high | very low | low | medium | high | very high |
| very low | 0.25 | 0 | 0 | 0 | 0 | 0.2 | 0 | 0 | 0 | 0 |
| low | 0.25 | 0.333333 | 0 | 0 | 0 | 0.2 | 0.25 | 0 | 0 | 0 |
| medium | 0.25 | 0.333333 | 0.666667 | 0.333333 | 0.333333 | 0.2 | 0.25 | 0.5 | 0.333333 | 0.25 |
| high | 0.25 | 0.333333 | 0.333333 | 0.666667 | 0.333333 | 0.2 | 0.25 | 0.25 | 0.333333 | 0.25 |
| very high | 0 | 0 | 0 | 0 | 0.333333 | 0.2 | 0.25 | 0.25 | 0.333333 | 0.5 |

| Chemical_Cate | unknown | | | | | | | | | |
| --- | --- | --- | --- | --- | --- | --- | --- | --- | --- | --- |
| Tox_fish_chem | high | | | | | | | | | |
| Tox_fish_oth_s | very low | | | | | low | | | | |
| Tox_emb_lev | very low | low | medium | high | very high | very low | low | medium | high | very high |
| very low | 0.4 | 0.25 | 0.25 | 0.2 | 0.2 | 0.25 | 0 | 0 | 0 | 0 |
| low | 0.2 | 0.25 | 0.25 | 0.2 | 0.2 | 0.25 | 0.5 | 0.333333 | 0.25 | 0.25 |
| medium | 0.2 | 0.25 | 0.25 | 0.2 | 0.2 | 0.25 | 0.25 | 0.333333 | 0.25 | 0.25 |
| high | 0.2 | 0.25 | 0.25 | 0.4 | 0.2 | 0.25 | 0.25 | 0.333333 | 0.5 | 0.25 |
| very high | 0 | 0 | 0 | 0 | 0.2 | 0 | 0 | 0 | 0 | 0.25 |

| Chemical_Cate | unknown | | | | | | | | | |
| --- | --- | --- | --- | --- | --- | --- | --- | --- | --- | --- |
| Tox_fish_chem | high | | | | | | | | | |
| Tox_fish_oth_s | medium | | | | | high | | | | |
| Tox_emb_lev | very low | low | medium | high | very high | very low | low | medium | high | very high |
| very low | 0.25 | 0 | 0 | 0 | 0 | 0.2 | 0 | 0 | 0 | 0 |
| low | 0.25 | 0.333333 | 0 | 0 | 0 | 0.2 | 0.25 | 0 | 0 | 0 |
| medium | 0.25 | 0.333333 | 0.666667 | 0.333333 | 0.333333 | 0.2 | 0.25 | 0.333333 | 0 | 0 |
| high | 0.25 | 0.333333 | 0.333333 | 0.666667 | 0.333333 | 0.4 | 0.5 | 0.666667 | 1 | 0.666667 |
| very high | 0 | 0 | 0 | 0 | 0.333333 | 0 | 0 | 0 | 0 | 0.333333 |

| Chemical_Cate | unknown | | | | | | | | | |
| --- | --- | --- | --- | --- | --- | --- | --- | --- | --- | --- |
| Tox_fish_chem | high | | | | | very high | | | | |
| Tox_fish_oth_s | very high | | | | | very low | | | | |
| Tox_emb_lev | very low | low | medium | high | very high | very low | low | medium | high | very high |
| very low | 0.2 | 0 | 0 | 0 | 0 | 0.2 | 0.2 | 0.2 | 0.2 | 0.2 |
| low | 0.2 | 0.25 | 0 | 0 | 0 | 0.2 | 0.2 | 0.2 | 0.2 | 0.2 |
| medium | 0.2 | 0.25 | 0.333333 | 0 | 0 | 0.2 | 0.2 | 0.2 | 0.2 | 0.2 |
| high | 0.2 | 0.25 | 0.333333 | 0.666667 | 0.333333 | 0.2 | 0.2 | 0.2 | 0.2 | 0.2 |
| very high | 0.2 | 0.25 | 0.333333 | 0.333333 | 0.666667 | 0.2 | 0.2 | 0.2 | 0.2 | 0.2 |

| Chemical_Cate | unknown | | | | | | | | | |
| --- | --- | --- | --- | --- | --- | --- | --- | --- | --- | --- |
| Tox_fish_chem | very high | | | | | | | | | |
| Tox_fish_oth_s | low | | | | | medium | | | | |
| Tox_emb_lev | very low | low | medium | high | very high | very low | low | medium | high | very high |
| very low | 0.2 | 0 | 0 | 0 | 0 | 0.2 | 0 | 0 | 0 | 0 |
| low | 0.2 | 0.4 | 0.25 | 0.25 | 0.2 | 0.2 | 0.25 | 0 | 0 | 0 |
| medium | 0.2 | 0.2 | 0.25 | 0.25 | 0.2 | 0.2 | 0.25 | 0.5 | 0.333333 | 0.25 |
| high | 0.2 | 0.2 | 0.25 | 0.25 | 0.2 | 0.2 | 0.25 | 0.25 | 0.333333 | 0.25 |
| very high | 0.2 | 0.2 | 0.25 | 0.25 | 0.4 | 0.2 | 0.25 | 0.25 | 0.333333 | 0.5 |

| Chemical_Cate | unknown | | | | | | | | | |
| --- | --- | --- | --- | --- | --- | --- | --- | --- | --- | --- |
| Tox_fish_chem | very high | | | | | | | | | |
| Tox_fish_oth_s | high | | | | | very high | | | | |
| Tox_emb_lev | very low | low | medium | high | very high | very low | low | medium | high | very high |
| very low | 0.2 | 0 | 0 | 0 | 0 | 0.2 | 0 | 0 | 0 | 0 |
| low | 0.2 | 0.25 | 0 | 0 | 0 | 0.2 | 0.2 | 0 | 0 | 0 |
| medium | 0.2 | 0.25 | 0.333333 | 0 | 0 | 0.2 | 0.2 | 0.25 | 0 | 0 |
| high | 0.2 | 0.25 | 0.333333 | 0.666667 | 0.333333 | 0.2 | 0.2 | 0.25 | 0.333333 | 0 |
| very high | 0.2 | 0.25 | 0.333333 | 0.333333 | 0.666667 | 0.2 | 0.4 | 0.5 | 0.666667 | 1 |
