## Supplementary figures for "Development of a hybrid Bayesian network model for predicting acute fish toxicity using multiple lines of evidence"

This supplementary file contains pictures (Figurea S.1-2) from the web interface, "A Bayesian network model to predict fish acute toxicity from multiple lines of evidence" (https://demo.hugin.com/example/FET).

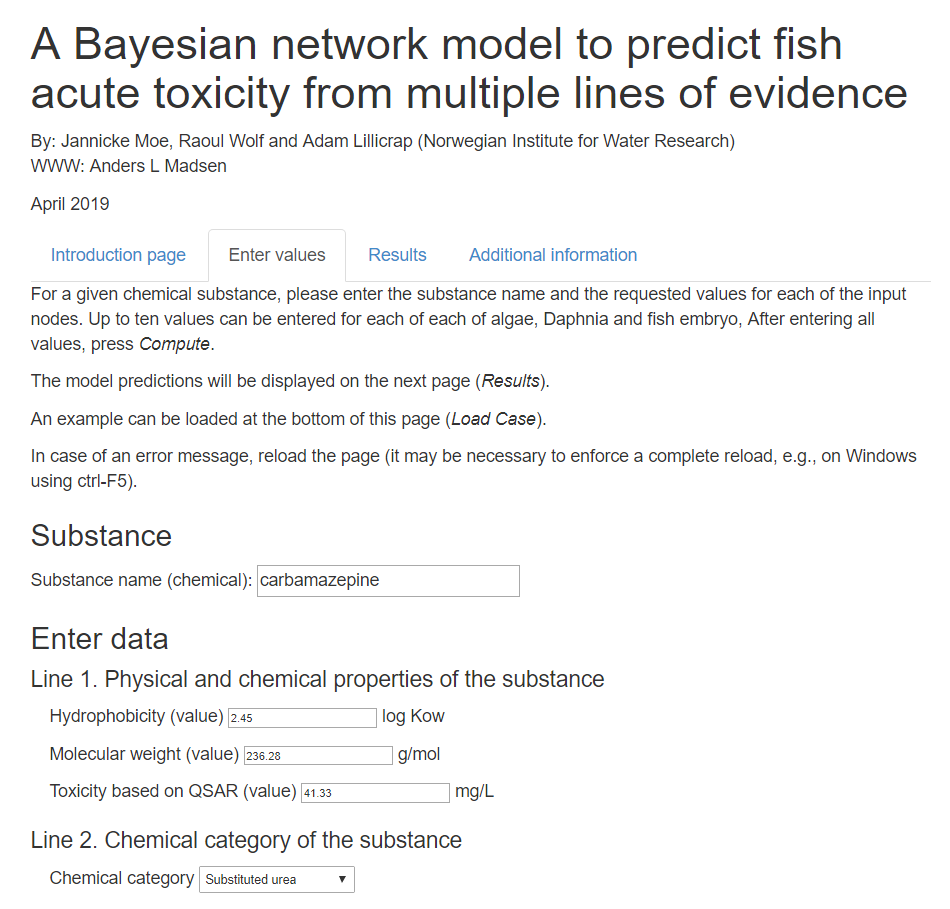

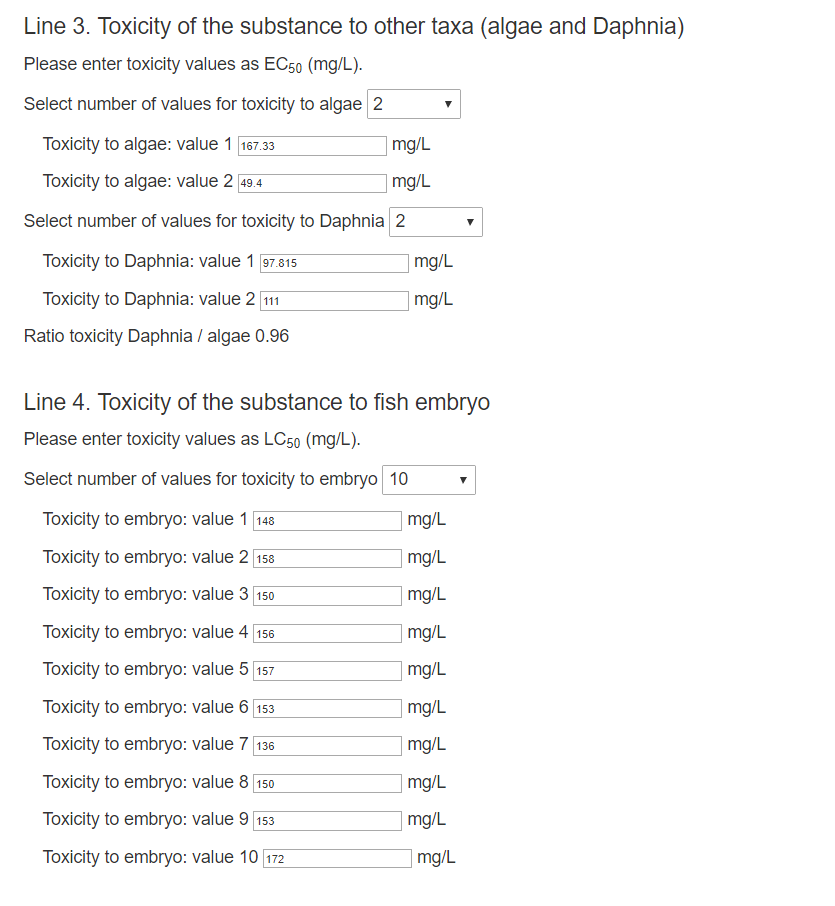

**Figure S.1.** Tab no. 2: "Enter Values". The values shown are entered by clicking the button "Load Case" further down on the page.

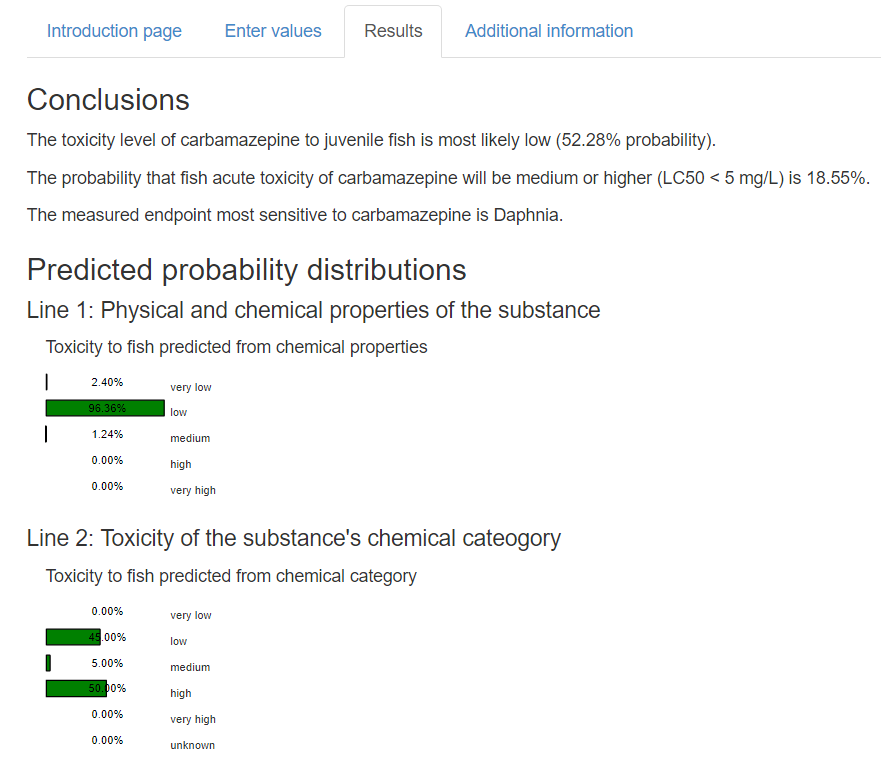

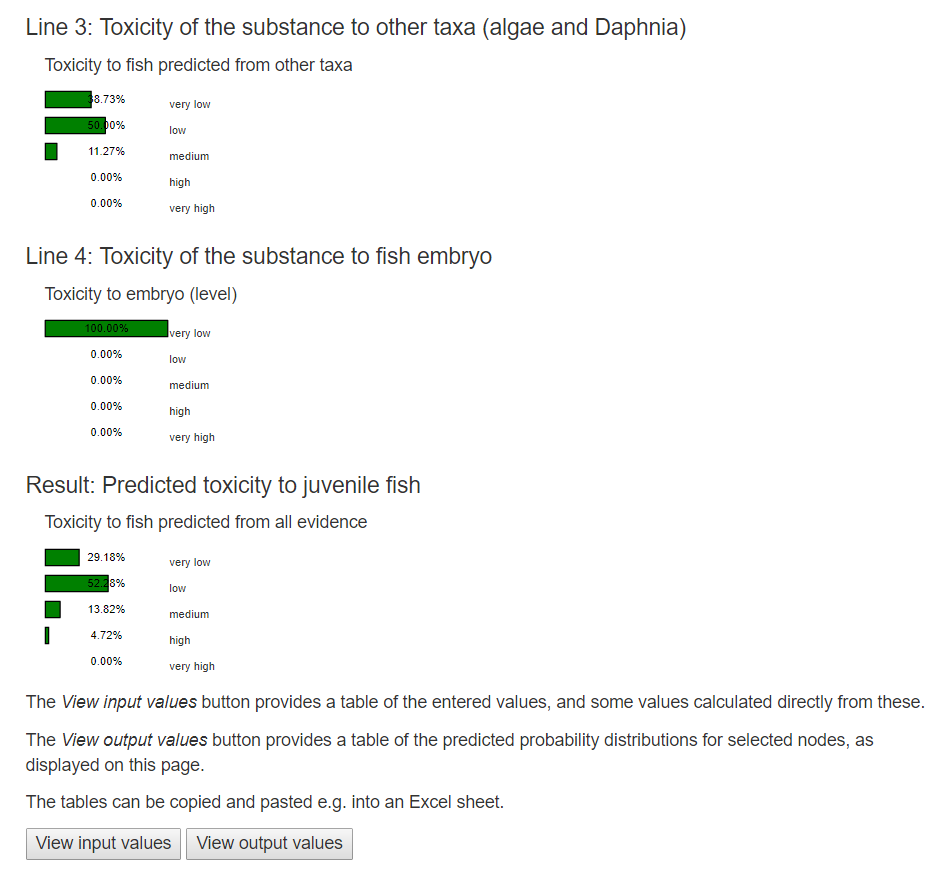

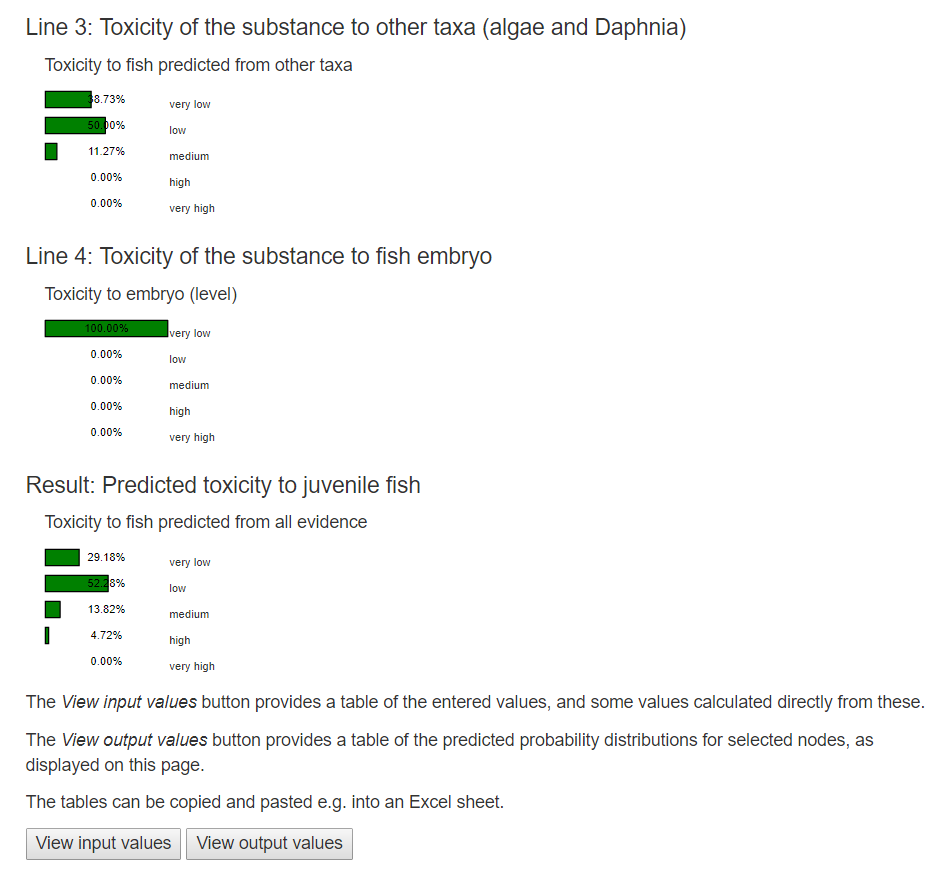

**Figure S.2.** Tab no. 3: "Results". The results are displayed as posterior probability distributions for the four lines of evidence separately and combined. The buttons "View Input Value" and "View Output Value" generate tables corresponding to Table 4a-b.
